# *Mesembryanthemum crystallinum* plasma membrane root aquaporins are regulated via clathrin-coated vesicles in response to salt stress

**DOI:** 10.1101/2022.03.16.484672

**Authors:** María Fernanda Gómez-Méndez, Rosario Vera-Estrella, Julio César Amezcua-Romero, Paul Rosas-Santiago, Eric Edmundo Hernández-Domínguez, Luis Alberto de Luna-Valdez, Omar Pantoja

## Abstract

The regulation of PIP-type aquaporins in the roots of plants has been identified as an important aspect in salinity tolerance. However, the molecular and cellular details underlying this process in halophytes remain unclear. Using free flow electrophoresis and label-free proteomics, we report that the increased abundance of PIPs at the plasma membrane of *Mesembryanthemum crystallinum* roots under salinity conditions is regulated by clathrin-coated vesicles (CCVs). To understand this regulation, we analyzed several components of the *M. crystallinum* CCVs complexes: clathrin light chain (*Mc*CLC) and subunits μ1 and μ2 of the adaptor complex (*Mc*AP1μ and *Mc*AP2μ). Co-localization analyses revealed the association between *Mc*PIP1;4 and *Mc*AP2μ and between *Mc*PIP2;1 and *Mc*AP1μ, observations corroborated by mbSUS assays, suggesting that aquaporin abundance at the PM is under the control of CCVs. The ability of *Mc*PIP1;4 and *Mc*PIP2;1 to form homo- and hetero-oligomers was tested and confirmed, as well as their activity as water channels. Also, we found increased phosphorylation of *Mc*PIP2;1 only at plasma membrane in response to salt stress. Our results prompt how root PIPs from halophytes are regulated through CCVs trafficking and phosphorylation, impacting their localization, transport activity and abundance under salinity conditions.

**One-sentence summary:** Abundance of plasma membrane aquaporins in *M. crystallinum* roots increases in response to salinity via a clathrin-coated vesicle-dependent mechanism.

## INTRODUCTION

Roots are the first plant organs to deal directly with salinity, therefore, the root’s ability to respond efficiently to this condition is essential to maintain water and nutrient absorption (Chen *et al*., 2011, Sanchez-Romera and Aroca, 2020). The absorption of water occurs mainly due to the hydraulic conductivity of the root (L_p_*_r_*), which largely depends on the function of aquaporins (AQPs) (Aroca *et al*., 2012). AQPs are channel-forming membrane proteins through which water moves into the cells according to their water potential (Javot and Maurel, 2002). Root AQPs participates in the short-distance distribution of water across cell membranes, via the cell-to-cell pathway and contribute to up 90% of the radial water flow, which is crucial to maintain water homeostasis at any cellular level (Afzal *et al*., 2016; Gambetta *et al*., 2017). The Plasma membrane Intrinsic Protein (PIP) family is the most studied family of AQPs affecting L_p_*_r_*, and its members are mainly localized at the plasma membrane (PM) (Chaumont *et al*., 2001; Boursiac *et al*., 2008; Danielson and Johanson, 2010). The PIP subfamily is divided into two phylogenetic groups, PIP1 and PIP2, which are further divided into different isoforms and can form homo- and hetero-oligomers (Fetter *et al*., 2004; Hacke and Laur, 2016). Functionally, PIP2s are more water permeable than PIP1s (Chaumont *et al*., 2000; Moshelion *et al*., 2002; Suga and Maeshima, 2004). Additionally, water permeability is regulated at the quaternary structure level, with PIP1-PIP2 hetero-oligomers having higher water permeability than the corresponding homo-oligomers (Yaneff *et al*., 2014; Ozu *et al*., 2018).

In glycophytes (salt-sensitive), there is a direct relationship between the decrease in L_p_*_r_* and a diminished abundance and/or activity of root PIPs; which is a key factor contributing to the rapid loss of water balance and stomatal closure in plants exposed to salinity (Sutka *et al*., 2011; Vaziriyeganeh *et al*., 2018). In *A. thaliana,* 100 mM NaCl treatment causes a drastic decrease in L_p_*_r_*, due to both, a decrease in PIPs transcripts expression, and most importantly, a drop of 40% for PIPs protein abundance at the PM (Boursiac *et al*., 2005). This downregulation is associated with the relocation of PIPs from the PM to intracellular compartments, a process that occur through endocytosis (Dhonukshe *et al*., 2007; Ueda *et al*., 2016). Furthermore, changes in the phosphorylation status can also modulate the activity, abundance, and subcellular localization of *At*PIPs during salt stress (Di Pietro *et al*., 2013; Verdoucq *et al*., 2014; Lee and Zwiazek., 2015). Phosphorylation at Ser283 in *At*PIP2;1 is required for its targeting to the PM (Prak *et al*., 2008), while its dephosphorylation promotes its internalization from the PM to endosomal compartments (Chevalier and Chaumont., 2015). Interestingly, short-term salt treatment causes a 30% decrease in the phosphorylation level in Ser283 and promotes the dephosphorylation of Ser280 (Prak *et al*., 2008). Based on these data, it is thought that PIPs regulation through changes in phosphorylation and internalization, is an early response glycophytes possess to decrease their abundance and activity at the PM, preventing water loss due to the osmotic stress caused by the presence of NaCl in the soil (Baral *et al*., 2015). Despite these early responses, glycophytes are not able to survive for long periods in NaCl concentrations greater than 50 mM (Munns, 2002), displaying the effects of dehydration and ionic damage over the course of a few days.

In contrast to glycophytes, halophytes withstand salinity and, show an increase in abundance of root AQPs in response to salt that results in water transport increases through cell membranes, allowing the stabilization of L_p_*_r_* (Alexandersson *et al*., 2005; Aroca *et al*., 2012; Vaziriyeganeh *et al*., 2018). In *E. salsuginea,* the abundance of root PIPs does not decrease when plants are exposed to 400 mM NaCl for 14 d (Shenghao *et al*., 2019). Also, in the moderately salt-tolerant plant *Brassica oleracea,* the abundance of root PIPs increases with NaCl treatments up to 90 mM for 15 days (d) (Muries *et al*., 2011; Zaghdoud *et al.,* 2013). Therefore, halophytes’ ability to maintain a higher abundance of root AQPs in long term exposure to salt, is an essential mechanism to maintain L_p_*_r_* and to overcome salinity stress (Aroca *et al*., 2001 and 2012; Alexandersson *et al*., 2005; Sutka *et al*., 2011). However, how the activity, abundance, and localization of PIPs at the PM are regulated in the roots of halophytes is little known.

*Mesembryanthemum crystallinum* is a halophyte known to maintain water balance during salt stress by differential regulation of AQPs (Kirch *et al*., 2000; Vera-Estrella *et al*., 2004). *M. crystallinum* has six identified PIP genes, *McPIP1;2*, *McPIP1;4* and *McPIP1;5* from PIP1 group, and *McPIP2;1*, *McPIP2;2* and *McPIP2;8* from PIP2 group (Yamada *et al*., 1995; Kirch *et al*., 2000). Protein abundance of *Mc*PIP1;2, *Mc*PIP2;1 and *Mc*PIP1;4 is up-regulated in roots under salt conditions (200 mM NaCl for 14 d), regardless of their transcript levels (Vera-Estrella *et al*., 2012). Also, *Mc*PIP2;1 is phosphorylated at PM of *Xenopus leavis* oocytes, which regulates its activity (Amezcua-Romero *et al*., 2010). However, its phosphorylation status in planta and its regulation in response to salinity has not yet been explored. Here we demonstrate how the CCVs-driven trafficking of *Mc*PIP1;4 and *Mc*PIP2;1 and phosphorylation of *Mc*PIP2;1 is associated with their increased abundance at the PM under salinity conditions.

## RESULTS

### Free flow zonal electrophoresis allows the identification of positively charge microsomes from roots of **M. crystallinum**

To investigate how root aquaporins from *M. crystallinum* (*Mc*PIPs) respond to salinity, we used Free Flow Zonal Electrophoresis (FFZE), a versatile method that allows the separation of cellular endomembranes based on their net surface charge (Heidrich and Hannig, 1989; Barkla *et al*., 2007 and 2018). We isolated root microsomes from *M. crystallinum* grown in hydroponic medium with (T) or without (NT) 200 mM NaCl for seven d. These samples were separated into 96 FFZE fractions, with a similar distribution profile between NT and T plants, showing a broad peak between fractions 30 to 60, and a smaller one between fractions 80 to 85 (Fig. 1A, black and gray lines, respectively). To identify the cellular compartments corresponding to these peaks, we analyzed these microsomal fractions by immuno-blot against endomembrane markers. We used the P-ATPase (AHA3) as the plasma membrane (PM) marker; the E subunit of the vacuolar translocating H^+^-ATPase (VHA-E) and tonoplast intrinsic protein (*Mc*TIP1;2) as tonoplast (TP) markers; Calreticulin (CTR1) as endoplasmic reticulum (ER) marker, and the reversibly glycosylated polypeptide (RGP1) as Golgi Apparatus (GA) marker. TP membranes were widely distributed between fractions 20 to 37, as previously described (Barkla *et al*., 2007), while PM-enriched fractions were located between fractions 38 to 46 (Fig. 1A and 1B, purple box). The ER and GA markers overlapped in fractions 47 to 55. Fractions 80 to 85 were characterized by being positively charged microsomes (PCM), given their migration towards the cathode of the FFZL system, forming an independent cluster that contained all the organelle markers tested (Fig.1A and 1C, green box). Protein content of the PCM fractions was higher in both T and NT root microsomes than in leaf microsomes, representing 19% of the total protein, while only 0.09% of the protein when leaf samples were used (Fig. 1A, blue line), indicating the PCM correspond to a root-specific endomembrane compartment, with a mixed nature. We observed that the TP marker showed a change in abundance in fractions 30 to 35 when compared T/NT (Fig. 1B *Mc*TIP1;2), however, protein markers in root microsomes from the rest of the fractions did not change significantly between T and NT samples (Fig. 1B). In contrast, TP, ER and GA membrane markers increased in abundance under salt conditions, in all PCM, while AHA3, the PM marker, only increased in fractions 83 and (Fig. 1C), suggesting a core response to salt, in these root-specific PCM fractions.

**Figure 1.**
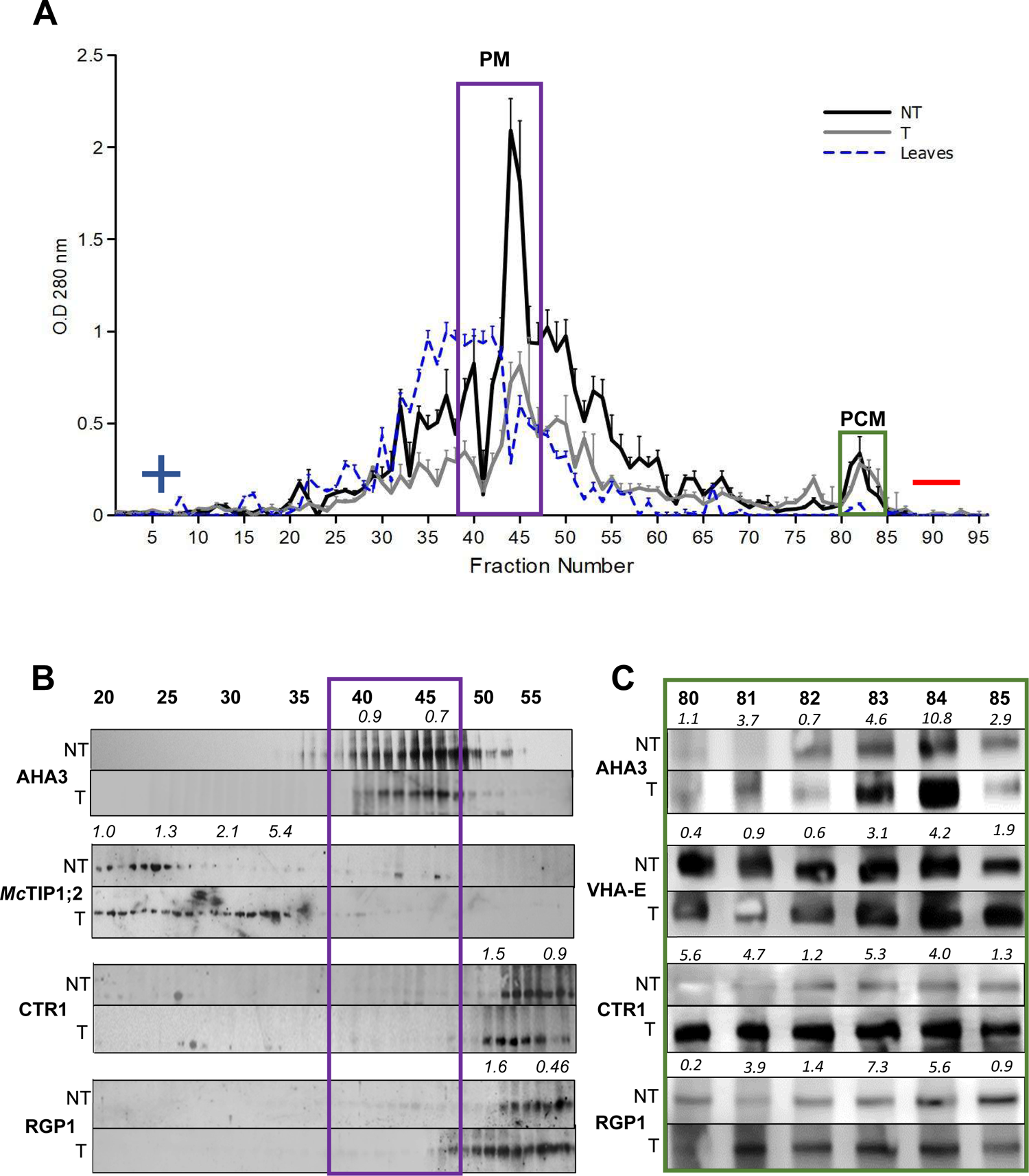
FFZE allows the identification of salt responsive positively charge microsomes (PCM) in roots of *M. crystallinum*. **A)** Protein profile of root microsomes from *M. crystallinum* treated (T, gray) or not (NT, black) for seven d with 200 mM NaCl, and leaf microsomes fractions are shown in blue. Positive and negative symbols represent the electrical field applied in the separation chamber, n=3. **B)** and **C)** Immunological detection of membrane markers AHA3 (PM, plasma membrane), *Mc*TIP1;2 and VHA-E (TP, tonoplast), CTR1 (ER, endoplasmic reticulum) and RGP1 (GA, Golgi apparatus), from fractions 20 to 55 and 80 to 85, respectively. PM enriched fractions are enclosed with a purple box, and PCM (positively charge fractions) are enclosed with a green box. The numbers in bold above each blot lane indicates the fraction number, and the number in italics show the difference in abundance (densitometry) between T/NT. The molecular weights of the immunodetected proteins are: AHA3, 100 kDa; *Mc*TIP1;2, 34 kDa; VHA-E, 31 kDa; CTR1, 57 kDa; RGP1, 41kDa. The blots are representative of three independent experiments.

### McPIPs protein abundance increases in association with vesicular trafficking proteins in roots of **M. crystallinum** during salinity conditions

In view of the apparent enrichment and salt response of the root-specific PCM fractions, we proceeded to analyze their protein composition by LC-MS/MS, from both, NT and T plants which resulted in the identification of 400 proteins (Supplementary Table S1). Analysis of these proteins demonstrated that 40% were shared between all six fractions in both T and NT samples, while only 15% were unique to single fractions in T and 10% in NT samples, suggesting that these fractions share a common endomembrane origin (Supplementary Fig. 1 and Supplementary Table S2). We performed a functional annotation and enrichment analysis of the *cell component* GO terms associated with the proteins identified in both NT and T PCM fractions (Fig. 2). These results showed that the GO terms: vesicle (GO:0005798), coated membrane (GO:0048475), Golgi-associated vesicle and membrane (GO:0005798, GO:0030660), endomembrane system (GO:0012505) and membrane protein complex (GO:0098796) were enriched in the PCM fractions from NT plants (Fig. 2A). The PCM fractions from T roots were enriched in the GO terms: clathrin coat of trans-Golgi network vesicles (GO:0030130), protein-containing complex (GO:0032991), bounding membrane of organelle (GO:0098588) and symplast (GO:0055044) (Fig. 2B). Also, the GO terms for Golgi Apparatus (GO: 0005794) and Plasma Membrane (GO:0005886) were enriched in both conditions (Fig. 2A and B). These results suggest that in both conditions, the proteins from the PCM fractions function in association with the GA and PM, and in salinity conditions, they associate with the TGN. In the PCM we identified proteins associated with CCVs, like the coat protein Clathrin Heavy Chain (CHC) and two μ subunits of the clathrin adapter complexes AP1 (APμ1) and AP2 (APμ2, Fig. 3A). The AP complexes are implicated in the formation of CCVs, with AP1 regulating the cargo transport from the TGN to vacuolar sorting or towards the PM during exocytosis; and AP2 operating in endocytosis (Arora and Van Damme, 2021; Yan *et al*., 2021). We also identified RABA1f, a member of a GTPase family involved in the movement of proteins between TGN and the PM, in all the PCM fractions (RABA1f, Fig. 3A, Asaoka *et al*., 2013). LC-MS/MS analysis also helped to identify the presence of several aquaporins, *Mc*PIP1;2, *Mc*PIP1;4, *Mc*PIP2;1, and *Mc*PIP2;5 in PCM fractions in both conditions (Fig. 3B), that were also among the 111 proteins responsive to salinity (Supplementary Table S3). LC-MS/MS data showed that salt exposure caused an increase in protein abundance for several PIPs (>2-fold; Fig. 3B and Supplementary Table S3), while the abundance of CCV associated proteins was marginally affected (Fig. 3A). According to the emPAI values, PIP2 aquaporins were more abundant than PIP1s; while among those associated with the CCV, CHC and RABA1f were more abundant than the APμs (Fig. 3). Identification of these two groups of proteins from the PCM fractions opened the possibility that aquaporin abundance between PM and intracellular pools could be regulated by CCVs in *M. crystallinum*, as is described for other species (Dhonukshe *et al*., 2007). To confirm this possibility, we analyzed the association between *Mc*PIPs and CCVs under salinity conditions by CCVs enrichment assays verified by immunoblotting using anti-CHC and anti-CLC (Fig. 4, CCVs). We observed the presence of PIP’s in CCVs from both T and NT roots, with *Mc*PIP1;2, *Mc*PIP1;4 and *Mc*PIP2;1 having an increased abundance >3-fold in T plants compared to NT plants (Fig. 4, CCVs). We also investigated possible changes in aquaporin abundance at the PM-enriched (fraction 43) and PCM (fraction 83) by immunoblot analysis. Under salinity conditions *Mc*PIP2;1 abundance increased by 3-fold in both PM and PCM fractions, while *Mc*PIP1;4 showed a 5-fold increase in the PM and 3-fold in PCM; for *Mc*PIP2;1 we observed a 2-fold increase in both, PM and PCM fractions (Fig. 4 PM and PCM, respectively). These data confirmed that the trafficking of *Mc*PIPs could be mediated or regulated by CCVs to increase their abundance at the PM and maintain/increase water conductivity under salinity conditions in *M. crystallinum* roots.

**Figure 2.**
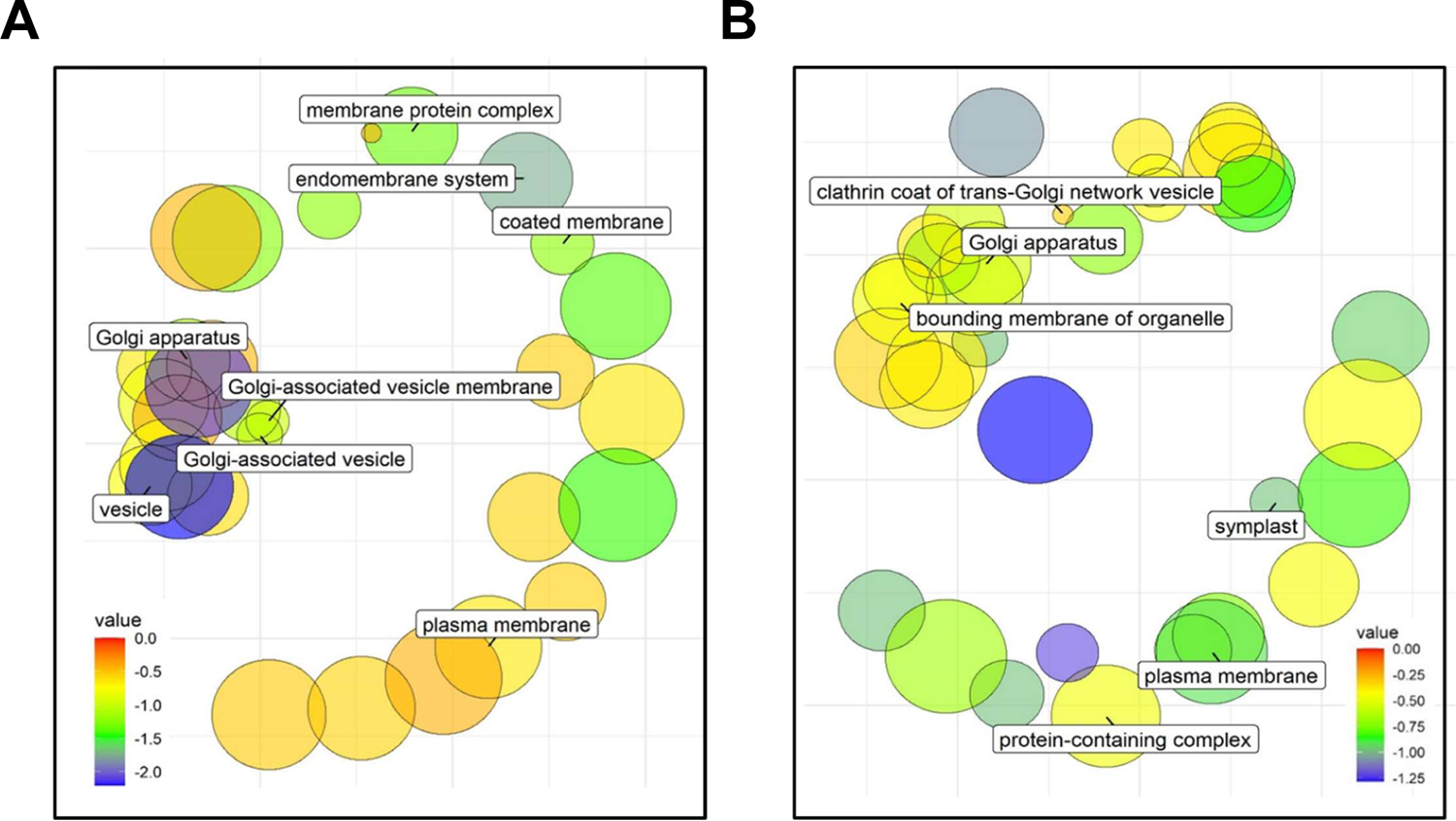
Cellular component-GO enrichment analyses of proteins identified in PCM fractions. Plot of significantly enriched GO terms related to cellular component in **A)** NT and **B)** T-PCM fractions. GO terms are represented by circles and are clustered according to semantic similarities to other GO terms (more general terms are represented by large size circles, and smaller circles show more specific GO terms). Color of the circle indicates the *p-value* of enrichment according to the scale. The most representative terms are described with labels. Endomembrane system (GO:0012505), coated membrane (GO:0048475), membrane protein complex (GO:0098796), plasma membrane (GO:0005886), vesicle (GO:0039982), Golgi apparatus (GO: 0005794) and Golgi-associated membrane (GO:0030660) and vesicle (GO:0005798), clathrin coat of trans-Golgi Network vesicle (GO:0030130), symplast (GO:0055044), protein-containing complex (GO:0032991) and bounding membrane of organelle (GO:0098588).

**Figure 3.**
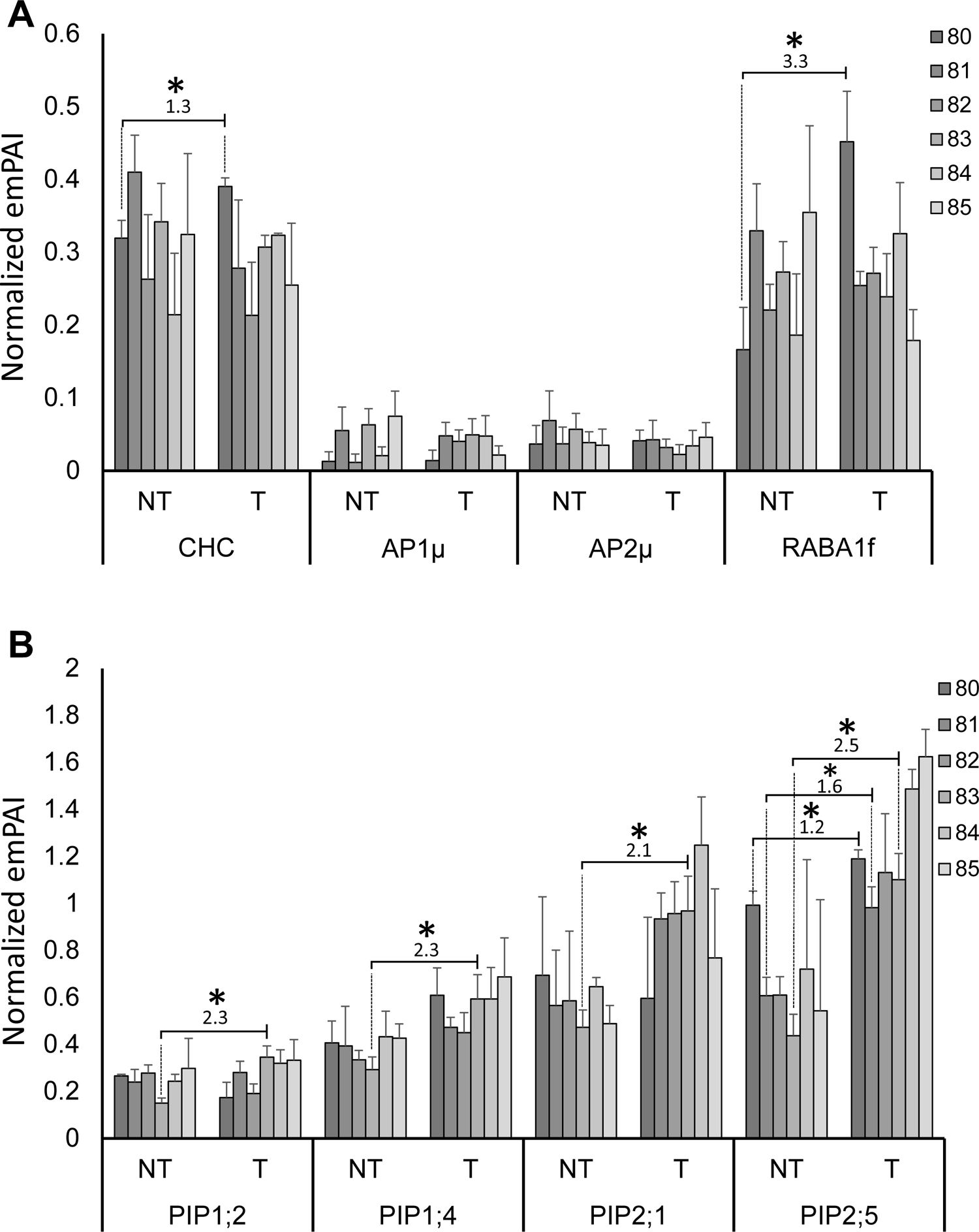
CCVs-related and PIPs identified in PCM fractions. Normalized emPAI (exponentially modified Protein Abundance Index) of **A)** CCV-forming proteins (CHC, AP1µ, AP2µ) and a GTPase (RABA1f) involved in transport between the TGN and the PM and **B)** Salt-responsive PIPs (PIP1;2, PIP1;4, PIP2;1 and PIP2;5) in NT and T conditions from 80 to 85 PCM fractions. Each fraction is represented with a color according to the legend. The asterisks indicate a significance *p-value* (< 0.05) obtained by a student’s t-test according to the emPAI of each protein; the number above the bar correspond to the fold change in T fractions.

**Figure 4.**
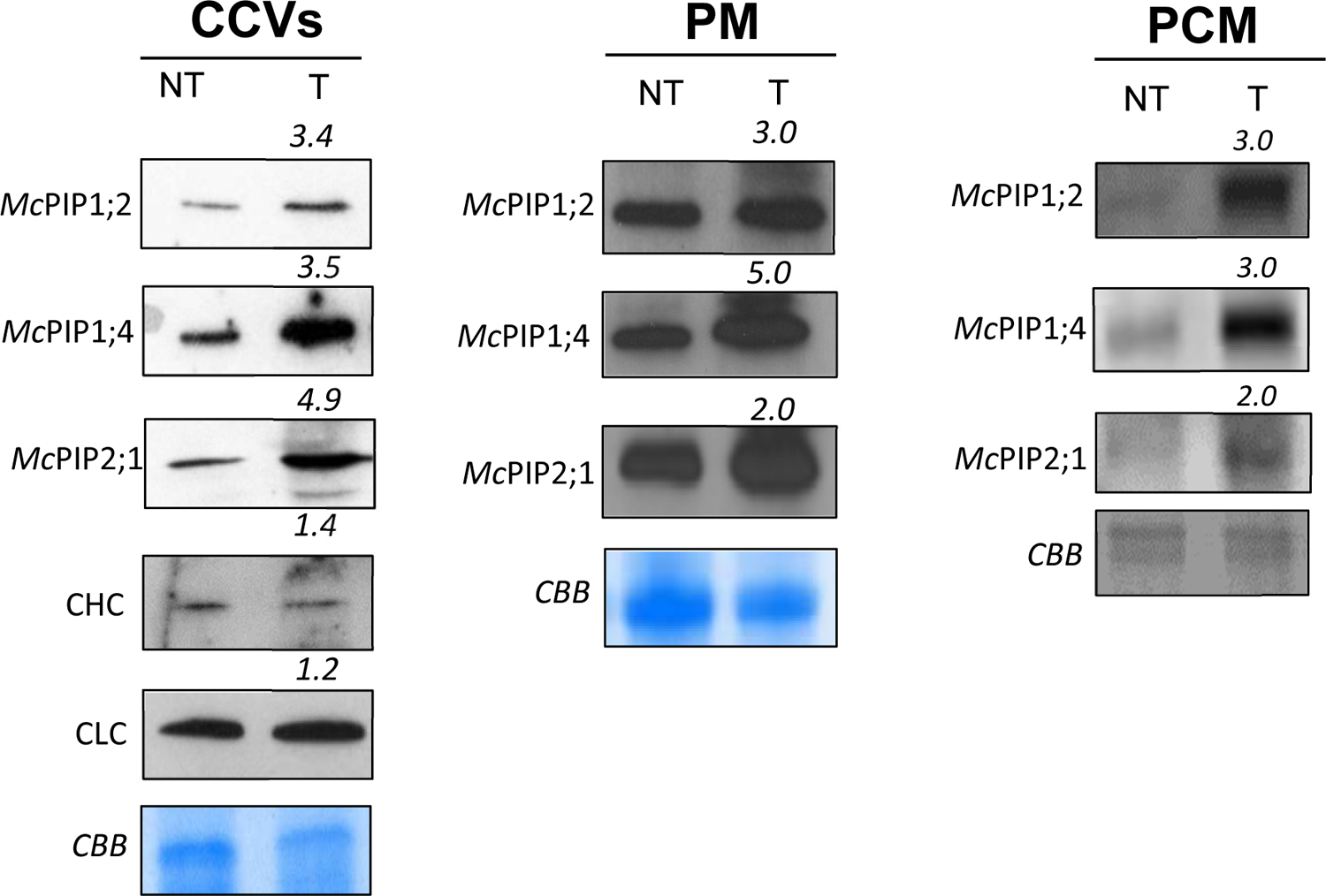
Salinity stimulates the abundance of *Mc*PIPs in different subcellular compartments in *M. crystallinum* roots. CCVs) Immunological detection of *Mc*PIP’s, CHC and CLC, in CCVs isolated from roots of *M. crystallinum* treated (T) or not (NT) for seven d with 200 mM NaCl. Immunological detection of *Mc*PIPs from PM-enriched fractions **PM)** and PCM-enriched fractions **PCM)** isolate by FFZE in T or NT plants. The number in italics above each blot lane shows the difference in abundance (densitometry) between T/NT. The molecular weights of the immunodetected proteins are 170 kDa, CHC; 58 kDa, CLC; 25 kDa *Mc*PIP1;2, 40 kDa, *Mc*PIP1;4, and 31/33 kDa for *Mc*PIP2;1. CBB (Coomassie Brilliant Blue) is show as a loading control. The blots are representative of three independent experiments.

### McCLC, McAP1μ, and McAP2μ are functional orthologs of the plant-identified clathrin vesicle components

Our results indicated that the increase in *Mc*PIPs abundance at the PM could be associated with their vesicular trafficking through CCVs, that function in clathrin-mediated endocytosis (CME) and post-Golgi trafficking. Given that a coordination of exocytic and endocytic trafficking is critical to regulate cell surface area and PM protein abundance and activity (Zhang *et al*., 2019; Yan *et al*., 2021; Arora and Van Damme, 2021), we decided to study *M. crystallinum* CLC (clathrin light chain), AP1μ, and AP2μ proteins, to assess the conservation of their corresponding functional traits in this plant. The mRNA sequences of putative *Mc*CLC, *Mc*AP1μ, and *Mc*AP2μ were obtained from a *M. crystallinum* RNASeq data set as described in *Material and methods* (Oh *et al*., 2015). Using these sequences, we performed phylogenetic analysis using 79 and 136 annotated CLC and AP1/2μ proteins, respectively. The analysis showed that *Mc*CLC shares an evolutionary origin closer to the CLC2 subfamily than to CLC1 and CLC3 homologues (Supplementary Fig. 2). Similarly, phylogenetic analysis showed that *Mc*AP1μ clusters in the AP1μ subfamily while *Mc*AP2μ does in the AP2μ group (Supplementary Fig. 3).

Next, we analyzed by confocal laser scanning microscopy (CLSM) the subcellular distribution of the CCV-associated proteins using the fluorescence-tagged proteins *Mc*CLC-mCherry, EYFP-*Mc*AP1μ, and EYFP-*Mc*AP2μ under the control of the 35S promoter, in abaxial epidermal peels from agro-infiltrated *N. benthamiana* leaves. In these experiments, we observed that *Mc*CLC-mCherry was distributed in the periphery of the cell and with dynamic intracellular vesicles (Fig. 5 and Supplementary Fig. 4A), consistent with previous observations of the *Arabidopsis* orthologs (Di Rubbo *et al*., 2013; Wang *et al*., 2013). EYFP-*Mc*AP1μ and EYFP-*Mc*AP2μ were observed near to the periphery of the cell (Fig. 5) and appeared as mobile puncta throughout the cytosol (Supplementary Fig. 4B and C). Overlapping of the two fluorescent images indicated the colocalization of *Mc*CLC-mCherry with EYFP-*Mc*AP1μ (Fig. 5A, merge) and *Mc*CLC-mCherry with EYFP-*Mc*AP2μ (Fig. 5B, merge). This observation was confirmed by quantifying the colocalization between *Mc*CLC-mCherry and EYFP-*Mc*AP1μ or EYFP-*Mc*AP2μ. According to the calculated linear Pearson correlation coefficient (*Rp = 0.97*) and the nonlinear Spearman correlation coefficient (*Rs* = 0.95) *Mc*CLC-mCherry and EYFP-*Mc*AP1μ did colocalized (Fig. 5A). *Mc*CLC-mCherry and EYFP-*Mc*AP2μ colocalization was also confirmed with a *Rp* = 0.88 and *Rs* =0.80 calculated values (Fig. 5B).

**Figure 5.**
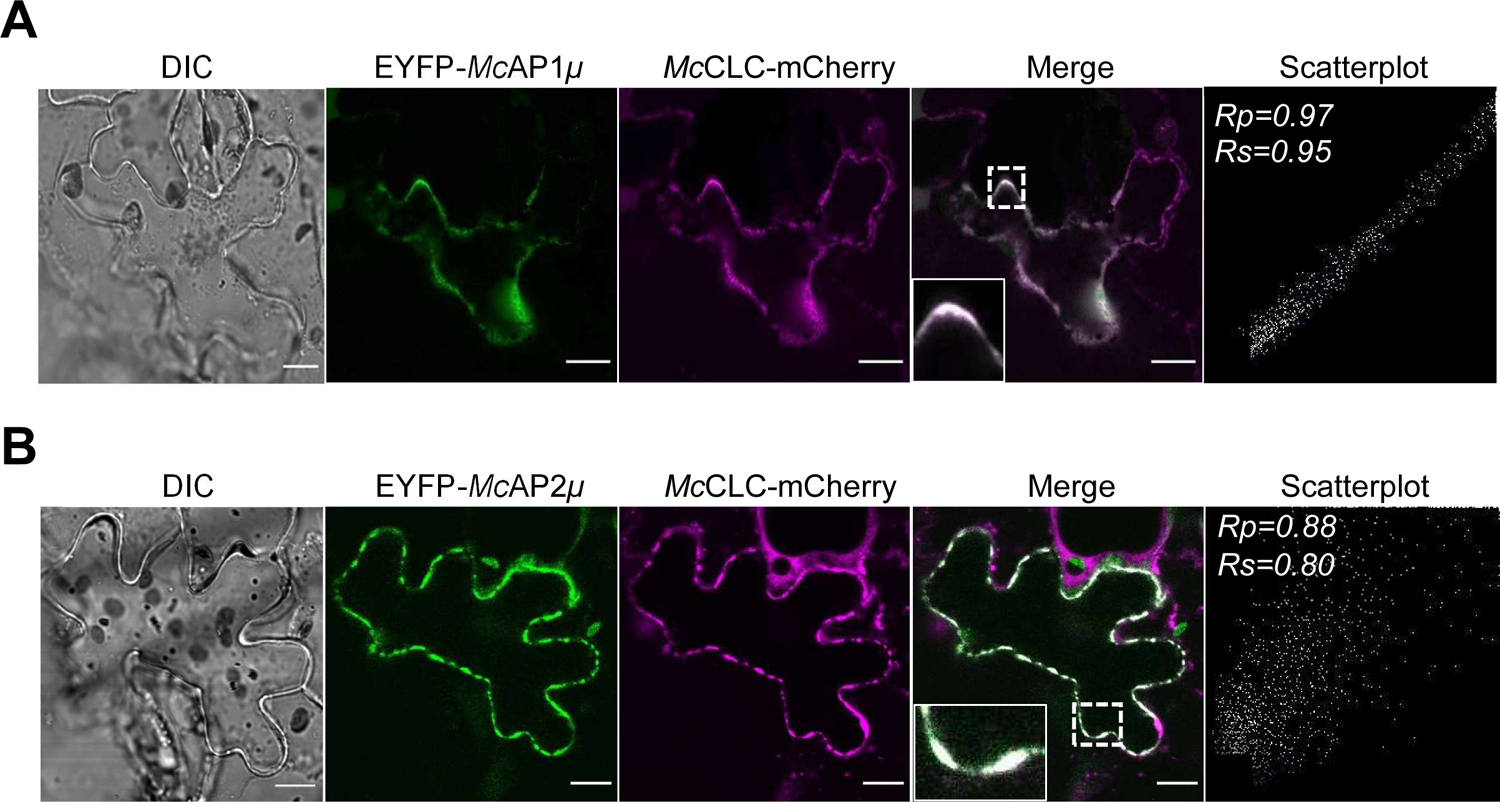
The CCV-associated proteins *Mc*AP1μ and *Mc*AP2 μ co-localize with *Mc*CLC. Confocal images from the abaxial epidermis of *N. benthamiana* leaves coinflitrated with **A)** EYFP-*Mc*AP1*µ* and *Mc*CLC-mCherry or **B)** EYFP-*Mc*AP2*µ* and *Mc*CLC-mCherry. Colocalization analysis was performed using the PSC plug-in (French *et al.,* 2008) in a region of interest (ROI) represented in the insets as areas of higher magnification. The obtained values for the linear Pearson correlation coefficient (*Rp*) and the nonlinear Spearman rank correlation coefficient (*Rs*) are shown in the corresponding scatterplots. Values for *Rp* and *Rs* ≥ 0.5 indicated colocalization between both proteins. Images are representative of at least three independent replicates *n*=3. DIC, bright field image. Scale bar 10 µM.

In other biological models, AP1μ is typically associated with post-Golgi trafficking and exocytosis, while AP2μ participates in endocytosis (Yan *et al*., 2021). To identify these mechanisms in *M. crystallinum* APμ genes, we performed colocalization analysis between both *Mc*APμ and FM4-64, a dye widely used to initially mark PM and subsequently follow the endocytic process (Rigal *et al*., 2015). Like the observations for EYFP-*Mc*AP1μ and EYFP-*Mc*AP2μ, we found the FM4-64 signal around the periphery of the cell (Supplementary Fig. 5). Furthermore, we observed a significant colocalization between EYFP-*Mc*AP2μ and FM4-64 in vesicle-like structures near the periphery of the cells (Supplementary Fig. 5A, *Rp* = 0.70; *Rs* = 0.72). In contrast, we did not detect colocalization between EYFP-*Mc*AP1μ and FM4-64 (Supplementary Fig. 5B, *Rp* = 0.34; *Rs* = 0.25), indicating that *Mc*AP2μ but not *Mc*AP1μ is associated to endocytosis. In addition, treatment with brefeldin A (BFA, 90 μM for 3h), a toxin that produces the agglomeration of GA, TGN and endosomal membranes in BFA bodies (Rigal *et al*., 2015, Robinson *et al*., 2008; Langhans *et al*., 2011), resulted in the relocation of EYFP-*Mc*AP1μ to BFA bodies (Supplementary Fig. 6A). This observation indicated the association of *Mc*AP1μ with TGN and post-Golgi trafficking, consistent with previous data from *Arabidopsis* orthologs (Park *et al*., 2013; Teh *et al*., 2013; Wang *et al*., 2013). Also, treatment with BFA resulted in the arrest of EYFP-*Mc*AP2μ in BFA bodies (Supplementary Fig. 6B). A similar observation for *At*AP2μ in the presence of BFA is observed in epidermal cells of *Arabidopsis* roots (Di Rubbo *et al*., 2013). Together, these results demonstrate that *Mc*CLC, *Mc*AP1μ, and *Mc*AP2μ are functional orthologs of the corresponding vesicle components that associate to form CCVs.

### McPIPs establish differential interactions with McAPμ proteins

To gain further insight into the trafficking of *Mc*PIPs through CCVs for exocytosis and/or endocytosis, we analyzed the distribution of *Mc*PIP1;4-*mC*herry and *Mc*PIP2;1-*mC*herry *in N. benthamiana* epidermis. We observed that both *Mc*PIPs distributed in discrete puncta in the cytosol and the periphery of the cell, suggesting an association with the PM (Supplementary Fig. 7A and B). To determine if both aquaporins located to the PM in this system, we plasmolyzed the cells expressing *Mc*PIP1;4-mCherry and *Mc*PIP2;1-mCherry to induce the formation of Hechtian strands, extensions of the PM connecting with the cell wall (Oparka *et al*., 1994, Cheng *et al*., 2017). Under this treatment, we observed the fluorescence associated to both *Mc*PIPs in thread-like structures linking the protoplasma with the cell wall, characteristic of the Hechtian strands (Supplementary Fig. 7C and D), confirming the presence of both proteins at the PM. We then assessed the co-localization between *Mc*PIP1;4-EYFP, and *Mc*PIP2;1-EYFP with *Mc*CLC-mCherry, by co-expression in *N. benthamiana* epidermis. In both cases, the signals associated with the fluorescent proteins were observed in the periphery of the cell, and in association with vesicular structures close to the PM. In the colocalization test, we found *Rp* and *Rs* values greater to 0.5 between *Mc*CLC-mCherry and *Mc*PIP1;4-EYFP (Fig. 6A*; Rp*=0.80; *Rs*=0.72), and *Mc*CLC-mCherry and *Mc*PIP2;1-EYFP (Fig. 6B*; Rp*=0.65; *Rs*=0.61). Consistent with our findings with PM-fractions and purified CCVs (Fig. 4), these results confirmed the association of *Mc*PIPs with CCVs at the PM.

**Figure 6.**
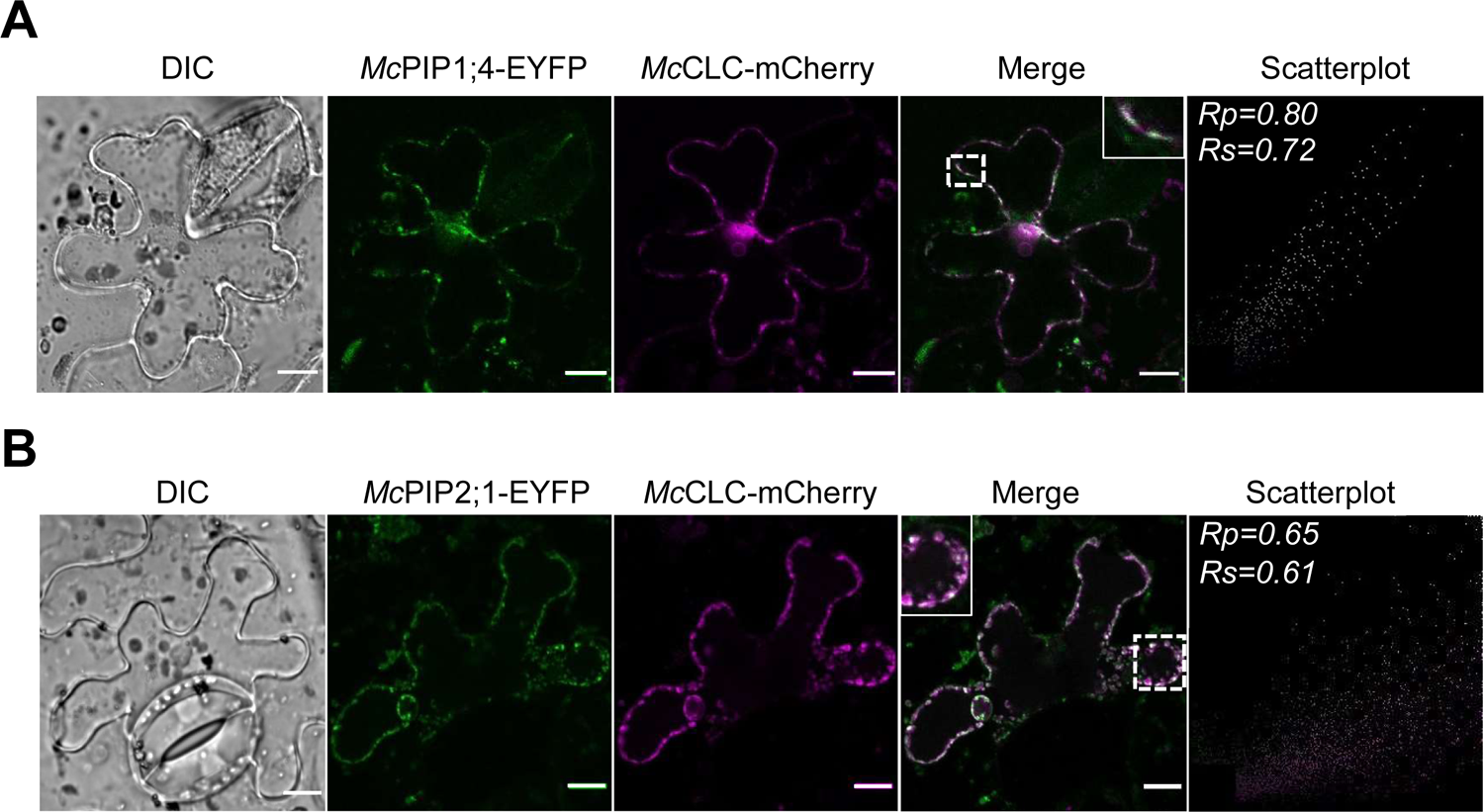
*Mc*CLC-mCherry co-localize with *Mc*PIP1;4-EYFP and *Mc*PIP2;1-EYFP. Confocal images of abaxial epidermal peels of *N. benthamiana* leaves coinflitrated with **A**) *Mc*PIP1;4-EYFP or **B**) *Mc*PIP2;1-EYFP and *Mc*CLC-mcherry. Colocalization analysis was performed using the PSC colocalization plug-in in Fiji (French *et al.,* 2008) in a region of interest (ROI) represented in the inset as areas of higher magnification. The obtained values for the linear Pearson correlation coefficient (*Rp*) and the nonlinear Spearman rank correlation coefficient (*Rs*) are shown in the corresponding scatterplot. Values for *Rp* and *Rs* ≥ 0.5 indicate colocalization between both proteins. All images are representative of at least three independent replicates. DIC, bright field image. Scale bar 10 µM.

Current knowledge indicates that adaptor complexes are involved in cargo loading to CCVs, and different AP complexes may not be fully redundant and participate in different intracellular transport pathways (Yan *et al*., 2021; Arora and Van Damme, 2021). To test the possibility that the APs associated with PIPs as direct cargoes, we carried out colocalization analysis. In these assays, we found that *Mc*PIP1;4-mCherry colocalized with EYFP-*Mc*AP1μ mainly in large intracellular vesicles, as indicated by the values of *Rp*=0.97 and *Rs*=0.96 (Fig. 7A). *Mc*PIP1;4-mCherry also colocalized with EYFP-*Mc*AP2μ in the periphery of the cell and intracellular vesicles (Fig. 7B*; Rp*=0.93; *Rs*=0.87). Similarly, we also found colocalization between *Mc*PIP2;1-mCherry and *Mc*AP1μ in the periphery of the cells and to a lesser extent in intracellular vesicles (Fig. 7C*; Rp*=0.88; *Rs*=0.88); while the colocalization between *Mc*PIP2;1-mCherry and EYFP-*Mc*AP2μ was observed near to the PM (Fig. 7D*; Rp*=0.71; *Rs*=0.73). These results indicated that both *Mc*PIPs and *Mc*AP1μ or *Mc*AP2μ coexist in subcellular compartments (near to the PM or in intracellular vesicles).

**Figure 7.**
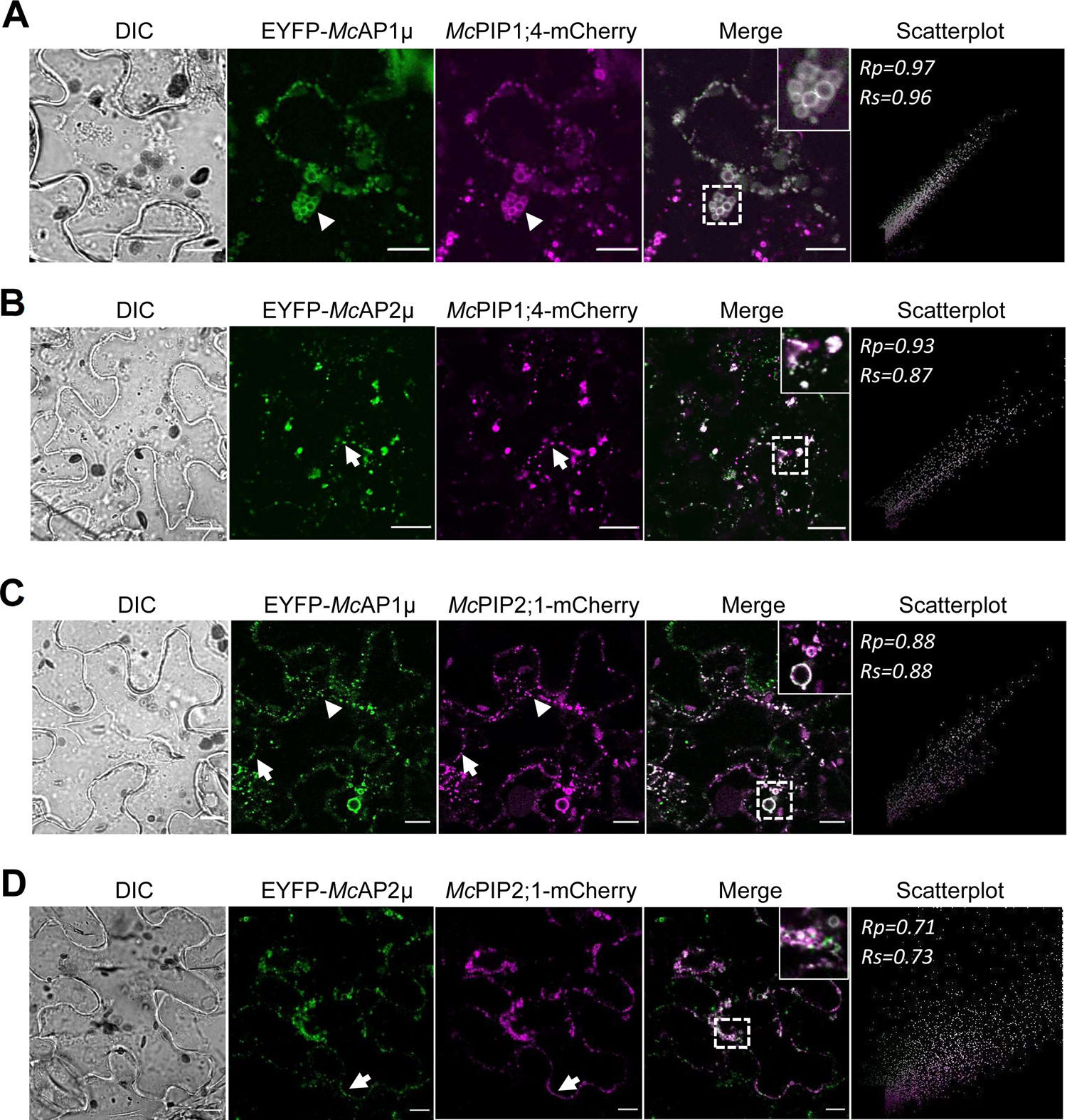
*Mc*PIP1;4-mCherry and *Mc*PIP2;1-mCherry co-localize with *EYFP-McAP1µ* or *EYFP-McAP2µ*. Confocal images of abaxial epidermal peels of *N. benthamiana* leaves co-infiltrated with **A)** EYFP-*Mc*AP1µ and *Mc*PIP1;4-mCherry; **B)** EYFP-*Mc*AP2µ and *Mc*PIP1;4-mCherry; **C)** EYFP-*Mc*AP1µ and *Mc*PIP2;1-mCherry and **D)** EYFP-*Mc*AP2µ and *Mc*PIP2;1-mCherry. Colocalization analysis was performed using the PSC colocalization plug-in in Fiji (French *et al.,* 2008) in a region of interest (ROI) represented in the inset as areas of higher magnification. The obtained values for the linear Pearson correlation coefficient (*Rp*) and the nonlinear Spearman rank correlation coefficient (*Rs*) are shown in the corresponding scatterplot. Values for *Rp* and *Rs* ≥ 0.5 indicate colocalization between both proteins. All images are representative of at least three independent replicates. DIC, bright field image. Arrow and arrowheads indicate -PM or intracellular vesicles signal association, respectively. Scale bar 10 µM.

To confirm the previous results, we identified that *Mc*PIPs contain the conserved motif YXXØ at their C-terminal, which is important for cargo loading to CCVs and direct interaction with the μ subunit of AP complexes (Madrid *et al*., 2001; Smith *et al*., 2017; Supplementary Fig. 8), indicating that *Mc*PIP1;4 and *Mc*PIP2;1 may establish direct protein-protein interactions with *Mc*AP1μ and/or *Mc*AP2μ. To probe this, we performed interaction assays using the mating-based split-ubiquitin system in yeast (mbSUS). The growth of diploid yeasts in the selection medium indicated the interactions between *Mc*PIP1;4 with *Mc*AP2μ, and *Mc*PIP2;1 with *Mc*AP1μ (0 µM, Fig. 8), however, growth was inhibited in the presence of 500 µM methionine (500 µM, Fig. 8), indicating that these interactions were not too strong. Moreover, LacZ reporter activity was observed by the development of a light blue signal in the presence of X-Gal, confirming the protein-protein interactions (LacZ, Fig. 8). Together, these results show that *Mc*PIP1;4 and *Mc*PIP2;1 and *Mc*APμs proteins coexist in the same cell compartments, but specific physical interactions are established only between *Mc*PIP1;4 with *Mc*AP2μ, and *Mc*PIP2;1 with *Mc*AP1μ.

**Figure 8.**
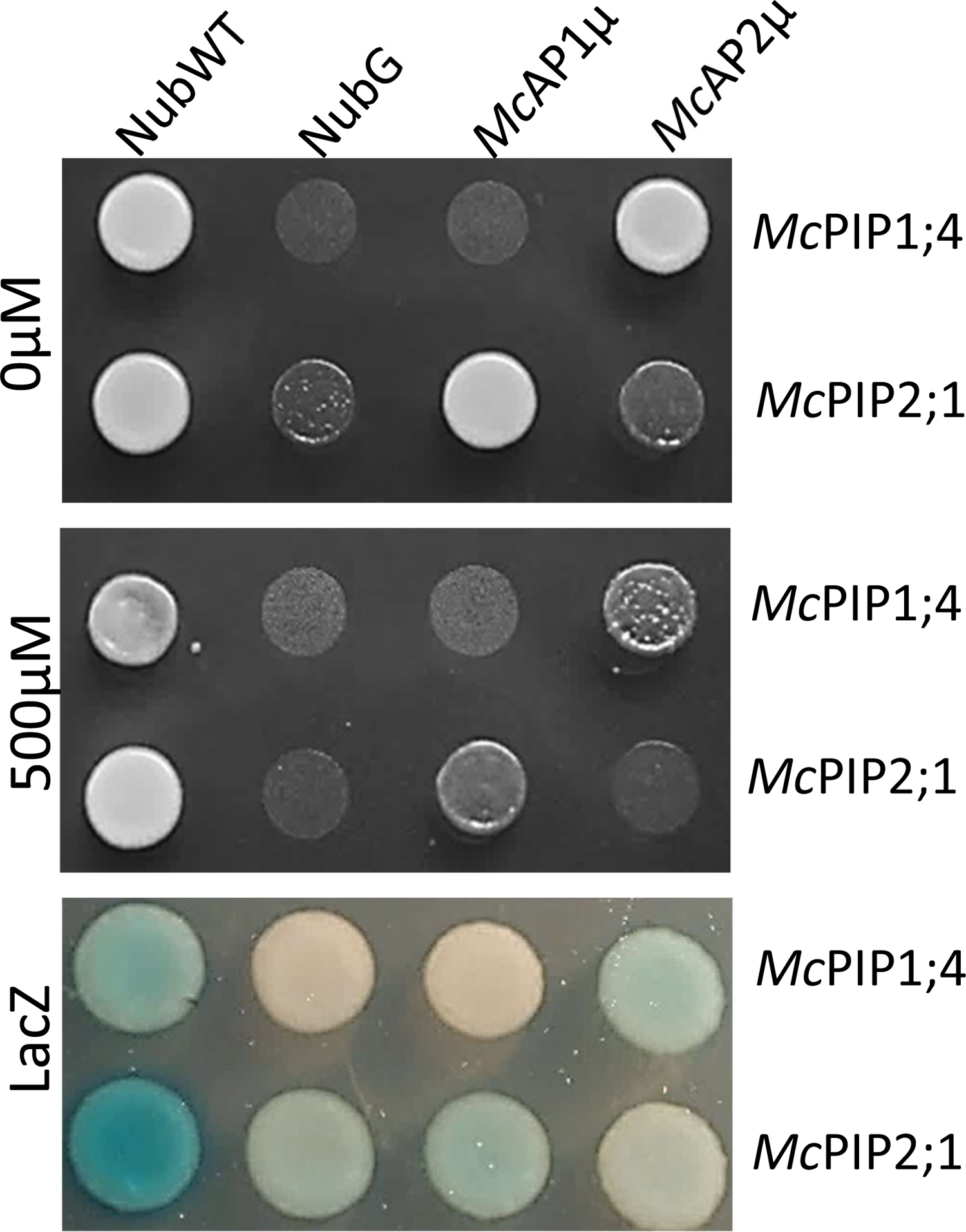
Aquaporins *Mc*PIP1;4 and *Mc*PIP2;1 interact with *Mc*AP2µ or *Mc*AP1µ, respectively. Protein-protein interaction between *Mc*PIP1;4 or *Mc*PIP2;*1* with *Mc*AP1µ and/or *Mc*AP2µ was assayed with mbSUS and confirmed by cell growth in the selective medium in the absence (0 µM) or in the presence of methionine (500 µM). Corroboration of the interactions was confirmed by activation of LacZ and revealed with X-Gal as substrate (LacZ). NubWT and NubG were used as false negative and false positive controls, respectively.

### McPIP1;4 and McPIP2;1 form hetero and homo-oligomers with differential water transport activity

Current evidence indicates that PIPs can form functional water-permeable oligomers in several biological models (Fetter *et al*., 2004; Temmei *et al*., 2005; Bienert *et al*., 2018; Ozu *et al*., 2018). To identify if *Mc*PIP1;4 and *Mc*PIP2;1 form hetero-oligomers, we first analyzed their co-localization in *N. benthamiana* epidermal cells co-infiltrated with *Mc*PIP1;4-YFP and *Mc*PIP2;1-*mC*herry. From these studies, we observed the colocalization of the two PIPs in the periphery of the cells (Fig. 9A; Rp= 0.84; Rs= 0.81), suggesting that the two proteins may interact at the PM. Furthermore, we analyzed the ability of *Mc*PIP1;4 and *Mc*PIP2;1 to form direct interactions using the mbSUS assay. Growth of diploid yeasts in the selection medium (0 μM, Fig. 9B) indicated that *Mc*PIP1;4 and *Mc*PIP2;1 form strong physical interactions that are stable even under stringent conditions (500 µM, Fig. 9B). We confirmed the interaction by testing the activity of the LacZ reporter, diploid yeasts developed a strong blue signal in the presence of X-Gal (LacZ, Fig. 9B). Since oligomerization between AQPs from different families does not occur, we used *Mc*TIP1;2 as a negative control in the mbSUS assay; the absence of yeast growth (*Mc*TIP1;2, Fig. 9B) indicated that the interaction between *Mc*PIP1;4 and *Mc*PIP2;1 is highly specific.

**Figure 9.**
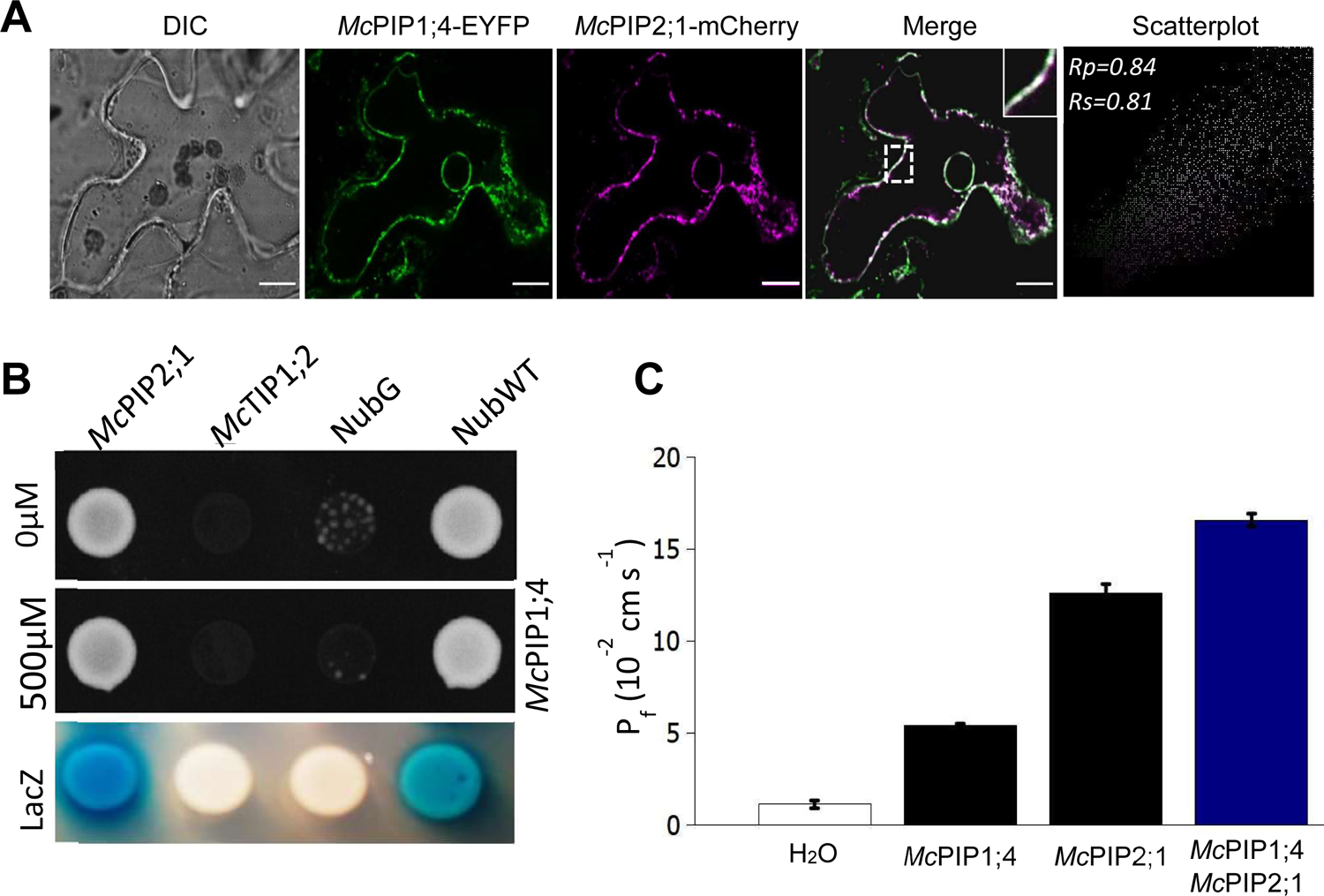
*Mc*PIP1;4 and *Mc*PIP2;1 interact and form heterooligomers with increased water permeability. **A)** Colocalization of *Mc*PIP1;4 and *Mc*PIP2;1 at the PM was confirmed by the calculated Pearson correlation (*Rp*) and the nonlinear Spearman rank correlation (*Rs*) coefficients higher than 0.5 (Scatterplot). Images are representative of a Z-stack and are representative of at least three independent transformations. DIC, bright field image. Scale bar 10 µM. **B)** Protein-protein interaction between *Mc*PIP1;4 and *Mc*PIP2;1 was indicated by the growth of yeast cells in the selective medium in the absence (0 µM) or in the presence (500 µM) of methionine. Corroboration of the interactions was demonstrated by activation of LacZ and revealed with X-Gal as substrate (LacZ). NubWT and NubG were used as false negative and false positive controls, respectively. The absence of growth with *Mc*TIP1;2 demonstrates the specificity of the interaction between *Mc*PIP1;4 and *Mc*PIP2;1. **C)** Water permeability (P_f_) of *Xenopus* oocytes injected with water (H_2_O), or with 25 ng mRNA of *Mc*PIP1;4 or *Mc*PIP2;1(Black bars). For the co-injected oocytes, 6.25 ng of *Mc*PIP2;1 and 18.75 ng of *Mc*PIP1;4 (1:3 ratio) were used (Blue bar). The values represent the mean ± SD (n = 10).

To assess the functionality of the *Mc*PIP1;4 and *Mc*PIP2;1 hetero-oligomers *in vivo*, we evaluated their ability to affect water permeability of membrane (P_f_) by co-injecting the corresponding cRNAs in *Xenopus* oocytes. Quantification of osmotic water permeability (P_f_) of the oocytes PM was assessed as described (Amezcua-Romero *et al*., 2010). Oocytes injected with 25 ng *Mc*PIP1;4 showed P_f_ values of 5.3 x 10^-2^ cm s^-1^, while those injected with *Mc*PIP2;1 had a P_f_ 12.6 x 10^-2^ cm s^-1^ (Fig. 9C, black bars). In contrast, oocytes co-injected with *Mc*PIP1;4 and *Mc*PIP2;1 showed a P_f_ of 16.5 x 10^-2^ cm s^-1^ (Fig. 9C, blue bar), which is 3.1 and 1.3 times larger than that for *Mc*PIP1;4 and *Mc*PIP2;1 alone, respectively. Control oocytes injected with ribonuclease-free water did not show any relevant volume change (Fig. 9C, white bar). Together, these results indicate that *Mc*PIP1;4 and *Mc*PIP2;1 do establish direct physical interactions to form functional water-permeable oligomers *in vivo*, that may localize at the PM.

### Phosphorylation of McPIP2;1 at the PM increases under salt treatment

Post-translational processes implicated in the control of abundance and activity of PIPs in the PM has been clearly demonstrated in glycophytes (Chaumont and Tyerman, 2014; Verdoucq *et al*., 2014). We previously reported that the phosphorylation of residues Ser123 and Ser282, positively affected the water permeability of *Mc*PIP2;1 in *Xenopus* oocytes (Amezcua-Romero *et al*., 2010). However, little is known about the *in-planta* phosphorylation status of *Mc*PIP2;1. In addition to the increase in protein abundance in salt-treated plants (Fig. 1 and Supplementary Fig. 9), a molecular weight shift in *Mc*PIP2;1 under both conditions was observed only in PM fractions (Supplementary Fig. 9; 33 kDa), while bands of lower molecular weight were observed at the TP and ER/GA fractions. (Supplementary Fig. 9; 31 kDa). To investigate if the molecular weight shift observed in PM fractions was due to a phosphorylation process, we performed *in vitro* phosphorylation and immunoprecipitation assays using the FFZE-purified TP, GA/ER, and PM-enriched fractions from NT and T *M. crystallinum* roots. We only detected phosphorylated *Mc*PIP2;1 in the PM-enriched fractions under both conditions (Fig.10A, NT and T), but not in the TP or ER/GA fractions. Furthermore, the phosphorylation level of *Mc*PIP2;1 was 4.7 times higher in salt-treated *M. crystallinum* roots (Fig. 10A, T). Immunodetection of *Mc*PIP2;1 in the same immunoprecipitated protein samples demonstrated that the same amount of *Mc*PIP2;1 was used across the different conditions (Fig. 10B), indicating that the results reflected a *bona fide* phosphorylation increase in T samples, and by the molecular weight shift in the *Mc*PIP2;1 only detected in the PM-enriched fractions (Fig. 10B and Supplementary Fig. 10). Moreover, we immunoprecipitated *Mc*PIP2;1 from PM-enriched fractions using specific antibodies and probed with antibodies that recognize the phosphorylated residues, anti-pSer, anti-pThr or anti-pTyr. We found that despite occurring in all three residues, *Mc*PIP2;1 phosphorylation was mainly directed to serine residues (Fig. 10C), as previously described (Amezcua-Romero *et al*., 2010). This confirms that *Mc*PIP2;1 is phosphorylated at the PM *in-planta* in NT and T conditions, modification that is enhanced by salinity conditions.

**Figure 10.**
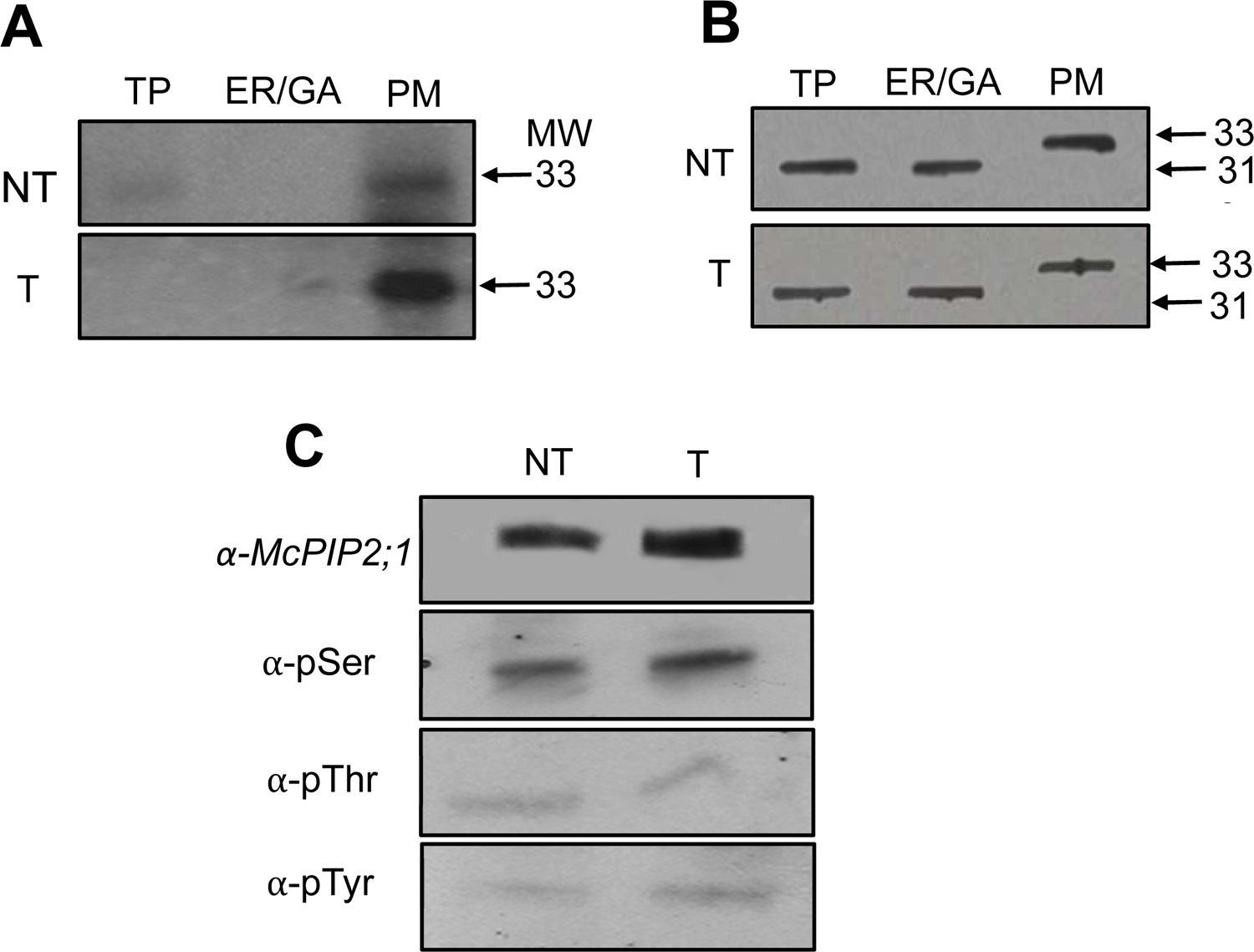
*Mc*PIP2;1 is phosphorylated in the plasma membrane of *M. crystallinum* roots. **A)** Autoradiograms of immunoprecipitated *Mc*PIP2;1 labeled with [γ-P] ATP in enriched TP, ER/Golgi and PM fractions from NT and T plants. A 4.7-fold stronger signal in salt-treated roots was derived from densitometry analysis between T and NT. **B)** Immunodetection of the 33 and 31 kDa isoforms of *Mc*PIP2;1 from immunoprecipitated fractions shown in **A**. The 33 kDa isoform (phosphorylated) was only observed in the PM fraction. **C)** Immunological detection of immunoprecipitated *Mc*PIP2;1 and its phosphorylated forms in enriched PM fractions. Fractions were probed with *α*-*Mc*PIP2;1, anti-phosphorylated serine (α-pSer), anti-phosphorylated threonine (α-pThr) or anti-phosphorylated tyrosine (anti-pTyr) antibodies. Representative results from two independent assays.

## DISCUSSION

From the first description of AQPs as water channels by Agre and coworkers in 1992 (Preston *et al*., 1992), we have gained understanding on the relevance of transmembrane water transport, and how they regulate cellular homeostasis under different physiological conditions. In plants, most water is transported along the cell-to-cell path, an increase AQP abundance or activity leads to an increase in L_p_*_r_*. Evidence indicates that AQP regulation achieves L_p_*_r_* changes faster (Suku *et al*., 2013; Gambetta *et al*., 2017). Here, we show that in the halophyte *M. crystallinum* increasing the abundance of PIPs at the PM is part of an adaptation process triggered by exposure to NaCl (Fig. 4). According to previous observations, this adaptation is only post-transcriptional, as it does not depend on increases in AQP transcript levels (Vera-Estrella *et al*., (2012). Similar responses to salt occur in roots of *Brassica oleracea,* a moderately salt-tolerant plant, where accumulation of PIP1 and PIP2 proteins at the PM was observed, regardless of the expression level of the corresponding transcripts (Muries *et al*., 2011). These data indicate that increasing the abundance of PIPs at the PM is an important feature of halophytes that may help to increase water transport through the cell-to-cell pathway, that is coupled with an increased accumulation of intracellular osmolytes (Adams *et al*., 1998, Vera-Estrella *et al*., 1999 and 2012), during the osmotic stress imposed by long term salinity conditions.

Our results also demonstrated that *Mc*PIP1;4 and *Mc*PIP2;1 form both homo- and hetero-oligomers, with the latter displaying greater permeability to water (Fig. 9 B and C). Additionally, because both *Mc*PIPs were found in PM and PCM enriched fractions (Fig. 4), together with their colocalization at the PM in tobacco epidermal cells (Fig. 9A), we propose that *Mc*PIP1;4/*Mc*PIP2;1 hetero-oligomers are formed at the PM of *M. crystallinum* root cells. Furthermore, as the abundance of both aquaporins increased in salt-exposed roots, it is possible that this condition induces the increase of both proteins as hetero-oligomers at the PM to improve water permeability at the root, to secure water uptake under osmotic stress coupled with intracellular osmolytes accumulation. Stimulated water permeability by the hetero-oligomerization of aquaporins has been reported in maize, where association of *Zm*PIP1;2 with different members of the *Zm*PIP2 subfamily has been observed, as well as with *Mp*PIP1;1 and *Mp*PIP2;1 from *Mimosa pudica* (Fetter *et al*., 2004; Temmei *et al*., 2005).

Previous studies have located PIPs within different vesicular trafficking routes, to finally locate at the PM and regulate their abundance during normal and stress conditions. However, most of these data come from glycophytes, and it is still an open question if these same routes and events occur in halophyte plants. In *M. crystallinum* we show that in purified CCVs the abundance of *Mc*PIP1;2, *Mc*PIP1;4 and *Mc*PIP2;1 increased in roots under salinity conditions (Fig. 4A) suggesting their trafficking through CCVs. This view is supported by the proteomics results of PCM fractions that demonstrated an association between the *Mc*PIPs and CCVs, concomitant with an increase in *Mc*PIPs abundance, possibly by an interplay between exo and endocytic events as indicated by the presence of proteins like CHC, RABA1F, *Mc*AP1μ and *Mc*AP2μ (Fig. 3) and the GO enrichment analysis (Fig. 2). These observations could be related with the increased abundance of aquaporins at the PM caused by salinity (Fig. 4), that contrary to what is observed in glycophytes, would suggest that exocytotic events dominate endocytosis in a halophyte, where sodium accumulation forces the osmotic uptake of water to maintain cell homeostasis. Analogous conclusions have recently been reached for *Arabidopsis* roots, where a coordinated regulation of exocytic and endocytic vesicle trafficking, dependent on CCVs and AP’s, was demonstrated as an important cellular mechanism to impact protein abundance and trafficking between the PM and the TGN (Yan *et al*., 2021). In contrast to *M. crystallinum* PIPs response, long-term salt stress in *A. thaliana* elicited the canonical glycophyte response, increasing endocytosis to limit the abundance of *At*PIP2;1 at the PM (Boursiac *et al*., 2005; Dhonukshe *et al*., 2007; Baral *et al*., 2015). Given the dynamic nature of the exocytic and endocytic trafficking, it is possible that rather than committing to a single pathway, it seems that PIPs are under a constitutive cycling process, although with different outcomes in halophytes and glycophytes to fine-tune their abundance at the PM during salt adaptation. As observed during short salt treatments were induced constitutive cycling of *At*PIP2;1 between intracellular compartments and PM occurred in the root (Luu *et al*., 2012; Martiniere *et al*., 2012).

Stronger evidence supporting the role of CCV in *Mc*PIPs trafficking was derived from the protein-protein interactions observed in the mbSUS and the co-localization of *Mc*PIP2;1 with *Mc*AP1μ, and *Mc*PIP1;4 with *Mc*AP2μ (Fig. 7). Moreover, the interactions *Mc*PIP2;1/*Mc*AP1μ and *Mc*PIP1;4/*Mc*AP2μ suggest that PIPs trafficking to and from the PM is tightly controlled (Fig. 8), that together with the hetero-oligomerization of *Mc*PIP1;4 with *Mc*PIP2;1 (Fig. 9), open the possibility that both aquaporins could interact indirectly with *Mc*AP1μ and *Mc*AP2μ, respectively.

If this occurs, it will allow their controlled trafficking within the CCVs-exocytic or endocytic pathway. PIP1 proteins do not reach efficiently the PM as homo-oligomers, however, formation of PIP1-PIP2 hetero-oligomers facilitate the arrival of PIP1s to the PM (Heymann and Engel, 1999; Fetter *et al*., 2004; Zelazny *et al*., 2007; Bienert *et al*., 2018). Therefore, the formation of hetero-oligomers may not only be involved in the exit of PIP1 from the ER as previously observed (Zelazny *et al*., 2007; Fetter *et al*., 2004), but also in their trafficking through the CCVs to and from the PM. Additional analyzes are necessary to confirm the interaction at the PM of *Mc*PIP2;1 and *Mc*PIP1;4 *in planta*.

In addition to the intracellular trafficking of *Mc*PIPs regulated by CCVs, the present and increase phosphorylation in the *Mc*PIP2;1 at the PM (Fig. 10A) suggests that this post-translational modification would be involved in the adaptation response to salinity, as phosphorylation on serine residue is known to positively regulate hydraulic conductivity at tissue level (Prado et al., 2013). Previously, we have demonstrated that phosphorylation at residues Ser123 and Ser282 of *Mc*PIP2;1 positively regulates the water permeability of the aquaporin (Amezcua-Romero *et al*., 2010). In contrast, phosphorylation of Ser283 in *At*PIP2;1 diminished in the PM when roots were exposed to 100 mM NaCl, a response associated with a salt-induced internalization of dephosphorylated *At*PIP2;1 (Park *et al*., 2008) and opposite to what we observed in *M. crystallinum*. These contrasting results open the possibility that regulation of PIPs abundance at the PM by phosphorylation in halophytes and glycophytes, plays an opposing role. In halophytes, PIPs increased abundance could be associated to the required demand in water uptake or flow as a consequence of Na^+^ absorption to maintain water homeostasis; while in glycophytes, a diminished abundance in PIPs would prevent water loss to reach the same goal.

## CONCLUSION

Under salinity conditions, *M. crystallinum* roots increase *Mc*PIP1;2. *Mc*PIP1;4 and *Mc*PIP2;1 abundance at the PM via CCVs dependent trafficking. Direct interaction between *Mc*PIP2;1/*Mc*AP1μ and *Mc*PIP1;4/*Mc*AP2μ, together with the oligomerization of the two aquaporins and the associated increase in water permeability, which suggest this type of interactions may be required to ensure halophytes maintain water uptake during long-term salt exposure. In addition to increased abundance, increased phosphorylation of *Mc*PIP2;1 at the PM is an important feature for the salt-tolerance ability of *M. crystallinum*. Our results demonstrate that plant adaptation to salinity conditions depends on the selective regulation of similar cellular components that lead to either, an increase or a decrease in the abundance of PM aquaporins to maintain cellular homeostasis according to the ecology of the species.

## MATERIALS AND METHODS

### Plant materials and growth conditions

*Mesembryanthemum crystallinum (*Adams *et al.,*1998) and *Nicotiana benthamiana* plants were grown from seed and cultivated in a soil mixture of peat moss, agrolite, and vermiculite in a proportion of 3:2:1, respectively. The soil mixture was supplemented with a controlled-release fertilizer (Osmocote Smart-release, The ScottsMiracle-Gro Company, US). Plants were maintained in a glasshouse under natural bright light and photoperiod. Glasshouse temperatures were varied between 12° C to 28° C. Three weeks after germination on soil mixture individual seedlings of *M. crystallinum* plants, were transferred to 1 L opaque tubs containing 800 mL of 1/2 strength Hoagland’s medium (Hoagland and Arnon., 1938) under constant aeration. The solution was changed weekly. Salt treatment was started three weeks after transplantation to hydroponics conditions and consisted in adding 200 mM NaCl in the Hoagland’s medium and keeping the plants in these conditions for seven d before harvesting. *N. benthamiana* were cultivated for six weeks and were watered with tap water every two d until use.

### Root microsomal membrane isolation

*M. crystallinum* root (30 g) was sliced into small pieces and placed directly into 300 mL of ice-cold homogenization medium, all subsequent operations were carried out at 4 °C. The homogenization medium consisted of 400 mM mannitol, 10% (w/v) glycerol, 5% (w/v) PVP-10, 1 mM PMSF, 30 mM Tris, 2 mM DTT, 5 mM EDTA, 5 mMMgSO4, 0.5 mM butylated hydroxytoluene, 0.25 mM dibucaine, 1 mM benzamidine, and 26 mM K^+^-metabisulfite, adjusted to pH 8.0 with H_2_SO_4_. Microsomal membranes were isolated as previously described (Vera-Estrella *et al*., 2004). Protein in microsomal preparations was measured by a modification of the Bradford method (Bradford., 1976), in which membrane protein was partially solubilized with 0.5% (v/v) Triton X-100 for 5 min before the addition of the dye reagent concentrate; the final concentration of Triton X-100 in the assay was 0.015%. BSA was employed as the protein standard. Membranes were frozen directly in liquid N_2_ and stored at −80 °C until use.

### Clathrin-coat vesicle isolation

For the CCVs isolation, we used the previously reported protocol (Reynolds *et al*., 2014) with some modifications. *M. crystallinum* roots (30g) was cut into small fragments and homogenized with 5 pulses of 30 seconds with breaks of 30 seconds between each pulse in a commercial blender (Waring, Mexico) with 300 mL of modified Clathrin Isolation Buffer medium (CIB, 400 mM Mannitol; 100 mM MES, 1mM EDTA, 3 mM EGTA, 1 mM PMSF, 0.5 mM BHT, 1 mM Benzamidine, 2 mM DTT and 26 mM K_2_S_2_O_5_). The homogenized tissue was filtered through six layers of gauze and centrifuged at 10,000 g_av_ for 25 min in the JA25.5 rotor (Beckman-Coulter, Mexico). The supernatant was then recovered and centrifuged at 30,000 g_av_ for one hour in the Ti45 rotor and in a Beckman L-80 ultracentrifuge (Beckman-Coulter, Mexico). The resulting pellet was re-suspended in 5 mL of modified CIB medium and centrifuged at 16,000 g_av_ for 10 min in 5415R refrigerated centrifuge (Eppendorf, USA). The resulting supernatant was the Sucrose Gradient Load (SGL), which was placed on a discontinuous sucrose gradient formed by concentrations of 10, 20, 35 and 50% of sucrose. This gradient was ultracentrifuged at 116,000 g_av_ for 50 min in the SW40 Ti rotor. The 10% phase and the 10/35% inter-phase were collected, and then subjected to a washing by ultracentrifugation at 200,000 g_av_ for 50 min with the CIB buffer, once centrifuged; 2.5 mL were collected with CIB buffer to obtain Deuterium-Ficoll Gradient Load (DFG-L); which was placed on top of a linear gradient of D2O-Ficoll, and then subjected to centrifugation for 16 hr at 80,000g_av_ in the SW40 Ti rotor. The resulting phases were collected and washed by ultracentrifugation at 174,000 g_av_ in the Ti 50.2 rotor. The final CCVs pellet was resuspended in 100 µL of CIB medium and store at −80 °C for analyze.

### Free flow zonal electrophoresis

Microsomal membranes were fractionated by Free Flow Zonal Electrophoresis (FFZE) using the BD FFE system (BD Proteomics, Germany), as previously described (Barkla *et al*., 2007 and 2017). Before fractionation, the microsomal sample was diluted 1:1 (v/v) in a separation medium and centrifuged at 13 000 rpm for 20 min at 4 °C. The supernatant was collected, adding 25 µL of 6 mM MgSO_4_, and 25 µL of 3 mM ATP. The sample (3 mg/mL) was injected continuously via a peristaltic pump at a rate of 1.2 mL/h using the anodic sample inlet. The media inlets of the chamber had the following buffer composition: inlets 2-6, separation medium (10 mM TEA, 10 mM acetic acid, 2 mM KCl, and 250 mM sucrose); inlets 1 and 7, stabilization medium (40 mM TEA, 40 mM acetic acid, 8 mM KCl, and 180 mM sucrose). The cathodic and anodic circuit electrolyte solutions consisted of 100 mM TEA, 100 mM acetic acid, and 20 mM KCl adjusted to pH 7.4 with NaOH, with 0.4% formaldehyde added to the anodic solution to prevent loss of chloride by anodic oxidation. The counter flow medium for inlets C1, C2, and C3 was the same as the separation medium. FFZE was performed in horizontal mode at a constant voltage of 750 V (118 mA), with a media and counterflow rate of 250 mL/h. The temperature during the run was maintained at 5 °C by the continual flow of coolant, below the glass separation plate, from a circulating water bath. Following separation in the chamber, membrane fractions were collected continually in 96 deep-well cultivated microtiter plates (4 mL/well; Sunergia Medical, VA). Fractions from sequential runs were pooled, and membranes were concentrated by centrifugation in a Beckman 55.2 Ti rotor in an L8-M ultracentrifuge at 100 000 g_av_ for 50 min at 4°C. Membrane pellets were resuspended in 50-100 mL of suspension buffer containing 250 mM mannitol, 10% glycerol (w/v), 10 mM Tris/MES pH 8, and 2 mM DTT and frozen in liquid N_2_ for storage at −80 °C. Separation by FFZE was monitored by collecting microtiter plates (250 mL/well) at several time points during the run and measuring protein (OD_280_) using a microplate scanning spectrophotometer (Power WaveX, Bio-Tek Instruments, VT).

### SDS-PAGE and immunoblotting

The denaturing electrophoresis from the isolated protein samples of *M. crystallinum* was performed as described (Vera-Estrella *et al*., 2012). Twenty-five µg of protein were precipitated in a solution of 50% ethanol and 50% acetone in a 1:50 ratio and incubated overnight at −30°C. Samples were centrifuged for 20 min at 14,000 g in a 5415R refrigerated tabletop centrifuge (Eppendorf, USA). The pellet was dried and suspended in 1X Laemmli buffer with 2.5% SDS (Laemmli, 1970). The samples were heated to 60°C for 5 min to allow their denaturation. They were loaded onto a 10% (v/v) polyacrylamide gel. SDS-PAGE electrophoresis was carried out in a buffer solution (25 mM Tris/HCl pH 8.3, 250 mM glycine, and 0.1% SDS) for 45 minutes at a voltage of 200 V, in a Protean Tetra Cell electrophoresis system (Bio-Rad). After SDS-PAGE, the transfer of the proteins to the nitrocellulose membrane was carried out employing a transfer buffer solution (Tris 25 mM, Glycine 190 mM, Methanol 25%), at 100 V for 75 min. After transfer, proteins were stained with Ponceau S 0.2% (w/v) and pictures of the blots were taken, to be used as loading controls (Sander *et al*., 2019); also, Coomassie blue (R-250) staining was use as loading control, when indicate. Next, the free protein-binding sites were blocked with a 5% solution of nonfat dry milk (Svelty, Nestlé, Mexico) dissolved in TBS (NaCl 150 mM, 50 Mm Tris / HCl at pH 7.5 and 0.05% sodium azide) for at least 1 h at room temperature. The nitrocellulose membranes were incubated for 12 h at room temperature with the specific primary antibodies; washed successively with TBS, TBS-tween-20, and TBS for 10 min and then are incubated for at least 2 h with the secondary antibody, IgG H + L conjugated to horseradish peroxidases at a 1: 5000 dilutions. After this time the membranes were washed successively as in the previous step.

### Primary and Secondary Antibodies

Peptides representing the second extracellular loop of the deduced amino acid sequences of *M. crystallinum McPIP1;2*; *McPIP1;4*; *McPIP2;1* or the carboxy terminus of *McTIP1;2*, were synthesized and coupled to keyhole limpet haemocyanin to generate antibodies against each aquaporin (Kirch *et al*., 2000). Antibodies against the P-ATPase (AHA3) from Arabidopsis were provided by R. Serrano (Pardo and Serrano, 1989). Calreticulin (CTR1) antibodies were against the protein from Arabidopsis (Nelson *et al*., 1997). RGP1 antibodies were kindly supplied by P.M. Ray (Dhugga *et al*., 1997). Polyclonal antibodies against the E subunit of the vacuolar translocating H^+^-ATPase (VHA-E) from barley were supplied by K.-J. Dietz (Dietz and Arbinger, 1996). Clathrin light chain (CLC2) antibodies were donated by Sebastian Y. Bednarek (Wang *et al*., 2013). Clathrin heavy chain (CHC) antibodies were from Agrisera. Monoclonal antibodies against phosphorylated serine (pSer), threonine (pThr) and tyrosine (pTyr) residues were from Life Technologies. Anti-pSer, anti-pThr and anti-pTyr antibodies were used at a dilution of 1:100; whereas antisera for *Mc*PIP1;2; *Mc*PIP1;4; *Mc*PIP2;1; *Mc*TIP1;2, AHA3, CTR, RGP1, VHA-E, CLC and CHC were used at a dilution of 1:1000. Immunocomplexes were detected using a 1:3000 dilution of peroxidase labeled goat anti-mouse IgG secondary antibodies (pSer, pThr, pTyr,) and developed using the enhanced chemiluminescent (ECL) detection substrate (GE, Healthcare, NJ). For the detection of all other primary antibodies, a 1:5000 dilution of anti-rabbit IgG secondary antibodies was employed and detected by chemiluminescence with the Luminata pack (Millipore, Billerica MA, USA).

### Protein precipitation by trichloroacetic acid and acetone

To PCM fractions obtained from the FFZE (4 separate independent experiments), were added 2% of the TE buffer solution pH 8.0, 0.06% sodium deoxycholate, 15% trichloroacetic acid were added for 2 h on ice. Then were centrifuged at 14000 g for 20 min at 4 °C. The supernatant was aspirated, and the pellet was resuspended in the initial volume with 90% acetone, and then incubated at −30°C overnight. After this, PCM fractions were centrifuged at 14,000 g for 20 min at 4°C. The supernatant was removed, and the resulting pellet was dried in a vacuum centrifuge (Savant, DNA120, Thermo Sci., Mexico) for 20 min. All samples were sent to the Montreal proteomics unit (Institut de Recherches Cliniques de Montréal) for processing and sequencing.

### Label-free proteomics and protein analyses

Data acquisition for the LC-MS/MS was performed using 11 cycles for each MS scan acquired on the Orbitrap. The spectra were extracted in the Mascot Daemon version 2.2.2 program. All MS/MS samples were analyzed using Mascot (Science Matrix, London, England) and X! Tandem (The GMP, https://thegpm.org/; version 2007.01.01.1). The Mascot was programmed to analyze the database nr_20101214. Both Mascot and X! Tandem were programmed to look for spectra that had a tolerance of 0.6 Da for mass and an ion tolerance of 12 ppm. Cysteine-derived methionine and iodine acetamide oxidations were considered as modifications in both programs. The Scaffold program (version 4.0 Proteome Software Inc., Portland, OR, USA) was used to validate the identifications of the peptides and proteins based on the MS/MS spectra and to identify the differentially regulated proteins for the no treatment and treatment conditions. Protein identifications was manually refined to accept only those with a probability greater than 99% and coverage percentage greater than 95%. Also, that they had at least two unique peptides, in at least two of the four experimental repetitions. Dimensional Venn diagrams of shared and unique proteins identified by LC-MS/MS was performed using nVenn code for the visualization of intersecting sets (Perez-Silva *et al.,* 2018). The functional annotation and the ontology enrichment (GO) analysis of the proteins were obtained with the Blast2Go bioinformatics tool (version 3.0.11, Conesa *et al.,* 2005 and 2008). The subcellular location and the functional annotation assigned by the Blast2Go were verified manually through Quick-GO of EMBL-EBI (Binns *et al*., 2009). The GO enrichment analysis under the GO domains cell component was performed using the topGO package (Alexa *et al*., 2006; Alexa and Rahnenführer, 2021) and the results were summarized and schematized using REViGO (Supek *et al*., 2011).

### In Vitro Phosphorylation and Immunoprecipitation

Purified microsomal membranes (30 µg of protein) were incubated in 100 µl phosphorylation medium containing 6 mM Tris/MES, pH 6.5, 5 mM MgSO4, 5 mM KCl, 2 mM dithiothreitol, 250 mM mannitol, 10 % (w/v) glycerol and 1 µl of [γ-32P]-ATP (10 µCi/µl) for 45 min at room temperature (25 °C). Afterward, membrane proteins were solubilized with 400 µl of NET-gel buffer (50 mM Tris/ HCl, pH 7.5, 150 mM NaCl, 1 mM EDTA, 0.1 % Nonidet P-40, 0.25 % gelatin, 0.02 % NaN3) and incubated overnight at 4 °C with anti-McPIP2;1 antibody (1/500 dilution) on a rotating table. Protein A-Sepharose CL-4B (20 µl, GE healthcare, NJ) was added and incubation continued for 2 h at 4 °C. Membrane proteins were pelleted at 12,000 g for 20 s. After two washings with NET-gel buffer, the protein pellet was washed with 10 mM Tris/HCl, pH 7.5, and 0.1 % Nonidet P-40, and centrifuged at 12,000 g for 20 s. The pellet was then resuspended with 20 µl of 2.5 % Laemmli sample buffer (Laemmli, 1970) and heated at 95 °C for 3 min. Protein A-Sepharose was removed by centrifugation at 12 000 g for 20 s. Immunoprecipitated proteins were resolved by 12.5 % SDS-PAGE. After staining, the gel was dried under vacuum for 3 h. Autoradiography was carried out by exposing the gel to Kodak X-OMAT film at −80 °C for 24 h.

### Gene cloning and plasmid constructions

Cloning of McPIPs was carried out using the Gateway technology (Thermo Fisher). Briefly, the full-length cDNAs of McTIP1;2, McPIP1;4 (Vera-Estrella *et al*., 2012) and McPIP2;1 (Amezcua-Romero *et al*., 2010) without stop codon, were amplified by PCR. PCR products were inserted into the pDNOR221 vector with a BP clonase II reaction kit. Entry vectors were verified by restrictions and sequencing. Transfer of the Gateway cassette from the entry vectors to the destination vectors, pX-EYFP-GW and pX-mCherry-GW (Rosas-Santiago *et al*., 2015) for tobacco expression vectors; and pMETYC_GW (Cub clones) and pXN32_GW (Nub clones) for yeast expression vectors in the mbSUS assay, was achieved with an LR clonase II reaction. The mRNA sequences for *McCLC* (Comp12569), *McAP1μ* (Comp25471) and *McAP2μ* (Comp 26254 and Comp27452) were obtained from the RNA seq data from Donh-Han *et al*., 2015. Oligonucleotides used can be found in Supplementary Material and Methods Table. *McCLC*, *McAP1μ* and *McAP2* were cloned using cDNA obtained from mRNA extracted from the root and KAPA3G plant PCR Kit product (KAPA BIOSYSTEM). PCR products were verified by sequencing. Cloning of *McCLC* was carried out using the Gateway technology (Thermo Fisher), to initially obtain the *pMcCLC-DONOR221* entry vector and later *McCLC-mCherry* expression vector, using BP and LR clonase II reactions, respectively. For cloning of *McAP1μ* and *McAP2μ*, pENTR/D-TOPO (Thermo-Fisher) cloning kit was used to obtain entry vectors that were verified by sequencing and recombined in the destination vector pEarleyGate-104 or pXN32_GW using LR clonase II kit.

### Evolutionary analysis by Maximum Likelihood method

The proteins sequences for CLC and AP protein orthologs were retrieved from NCBI (https://www.ncbi.nlm.nih.gov/) with the BLAST tool, using the *A. thaliana* sequences as queries and setting an expect threshold of 0.05. We retrieved a total 78 AP orthologues from seven species and 135 CLC orthologues from 49 species. Additionally, we used homolog proteins from *Synechococcus sp.* and *Streptococcus pyrogenes* as outgroups for the evolutionary analysis of APs and CLCs respectively.

Evolutionary analyses were conducted in MEGA X (Kumar *et al*., 2018). The evolutionary history was inferred by using the Maximum Likelihood method of Le and Gascuel., 2008 model for AP, and Whelan and Goldman., 2001 model for CLC proteins. Highest likelihood values of −14641.66 for AP and −15452.87 for CLC trees were found. Discrete Gamma distributions were used to model evolutionary rate differences among sites; five categories were set with (+G, parameter = 2.5496) for the APs; while for CLCs (+G, parameter = 0.7554) were used. All positions with less than 75% site coverage were eliminated. In the case of the AP proteins, the rate variation model allowed for some sites to be evolutionarily invariable ([+I], 2.15% sites).

### Transient Expression in N. benthamiana

The following constructions, *pMcPIP2;1-EYFP*, *pMcPIP1;4-EYFP*, *pMcCLC-mCherry*, *pEYFP-McAP1μ*, *pEYFP-McAP2μ* were independently transformed into *A. tumefaciens* strain GV3101 and selected on 30 mL of LB plates with the combination of suitable antibiotics; rifampicin (50 μg ml^-1^) and spectinomycin (50 μg ml^-1^) for pMcPIP2;1-EYFP, pMcPIP1;4-EYFP, pMcCLC-mCherry; or rifampicin (50 μg ml^-1^) and kanamycin (50 μg ml^-1^) for pEYFP-McAP1μ, pEYFP-McAP2μ. Overnight cultures of single colonies were grown at 28 °C on liquid LB with the appropriate antibiotics and were twice washed and resuspended with sodium phosphate buffer pH 7, 0.1 mM acetosyringone (Sigma), and 28mM glucose at a final O.D_600_ of 0.3 to 0.5 for each colocalization test or individual construction transformation. The infiltration of *N. benthamiana* leaves was made with an insulin syringe without a needle in the abaxial face. After infiltration, the plants were watered and kept at low illumination for four days until microscopic observation.

### Confocal laser scanning microscopy

Abaxial epidermal peels were made of mature infiltrated leaves and placed onto microscope slides in water, covered with a cover slide. For plasma membrane staining and monitoring of endocytosis, abaxial epidermal peels were incubated in 10 μM of FM4-64 (Invitrogen) for 5 min to ensure saturation of the PM with the dye. Observation of Hechtian strands was achieved through addition of 1M NaCl to the epidermal peels prior imaging. For Brefeldin A (BFA, Invitrogen) treatment, BFA was dissolved in DMSO to obtain a stock solution; 90 μM of BFA was infiltrated with an insulin syringe without the needle 3 h before imaging. Confocal laser scanning microscopy (CLSM) was made using an inverted multiphoton confocal microscope (Olympus FV1000) equipped with a x60 oil immersion objective (NA 1.3). Excitation wavelengths were 488 nm (argon laser) for EYFP and 543 nm (diode laser) for mCherry and FM4-64. Emission was detected at 520 nm (BA505-520) for EYFP and 572 nm for mCherry (BA560-620) and FM4-64 (BA560-660). All imaging was performed using sequential scans to prevent bleed-through fluorescence. Results are representative images from >3 cells from at least three different independent biological transformations.

### Image Analysis

Colocalization quantification of CLSM images was done in a region of interest (ROI) in at least ten optical sections of the Z-stack in each biological replicate. The PSC colocalization plugin of ImageJ (Schindelin *et al*., 2012), was used to calculate Pearson and Spearman correlation coefficients after adjusting the background level according to instructions (French *et al*., 2008). The reported images were adjusted for better observation; however, image analyzes were carried out, on the unmodified images.

### mbSUS assay

Yeast media were prepared as previously described (Lalonde *et al*., 2010). The THY.AP4 (*MATa ura3, leu2, lexA::LacZ::trp1 lexA::HIS3 lexA::ADE2*) and THY.AP5 (*MATα URA3, leu2, trp1, his3 loxP::ade2*) yeast strains were transformed with the pMcPIP1;4-Cub, pMcPIP2;1-Cub, pMcAP1m-Nub and pMcAP2m-Nub or pMcPIP1;4-Nub constructs. Cub constructs and Nub constructs were transformed into THY.AP4 and THY.AP5 yeast strains, respectively (Obrdlik *et al*., 2004), employing the LiAc protocol previously described (Lalonde *et al*., 2010). Those fusions that did not interact with the soluble NubWT, which has a strong affinity for the Cub domain, were considered as false negatives; while those that did interact with the NubG corresponded to false positives.

### Heterologous expression in Xenopus laevis oocytes

The cDNAs encoding McTIP1;2 (Vera-Estrella *et al*., 2004), McPIP2;1 (Amezcua-Romero *et al*., 2010), and McPIP1;4 (Vera-Estrella *et al*., 2012) were cloned into the pGEM-HE vector. cRNAs from McTIP1;2, McPIP2;1 and McPIP1;4 were prepared with the T7 RNA polymerase after linearization with NheI, using the mMESSAGE mMACHINE in vitro transcription kit (Ambion, Austin, TX). Defolliculated oocytes were injected with 46.0 ml of DEPC-H_2_O or the corresponding cRNA (1 ng/nl) using a NANOJECT II automatic injector (Drummond, Broomall, PA). Oocytes were incubated in iso-osmotic (200 mosmol/kg) Barth’s solution (10 mM HEPES-NaOH, pH 7.4, 88 mM NaCl, 1mM KCl, 2.4mM NaHCO_3_, 0.33mM Ca(NO_3_)2, 0.41 mM CaCl_2_, 0.82 mM MgSO_4_) for 4–5 days before swelling assays.

### Oocyte Swelling assays

Swelling assays were performed by transfering of the oocytes from iso-osmotic to hypo-osmotic (100 mOsmol kg^-^) Barth’s solution, according to Amezcua-Romero *et al.,* 2010. After the transfer, oocyte volume changes were recorded with a Hitachi KP-D50 color video camera (Hitachi Denshi, Woodbury, NY) on a Nikon Eclipse TE 300 inverted microscope (Nikon, Mexico). Images were captured at intervals of 10 s for 2 min and digitized by the Image-Pro Plus software (Version 4, Media Cybernetics, Silver Spring, MD). Data were analyzed as the proportional change in volume and normalized to the initial volume at time 0. All the assays were performed at room temperature (25 °C). Volume changes values were calculated for each independent set of experiments.

## SUPPLEMENTAL DATA

The following supplemental materials are available:

**Supplementary Figure 1.** Positively-charge fractions in NT and T conditions shared a significant number of proteins.

**Supplementary Figure 2.** CLC is conserved in *M. crystallinum*.

**Supplementary Figure 3.** AP1µ and AP2µ are conserved in *M. crystallinum*.

**Supplementary Figure 4.** Intracellular localization of *Mc*CLC-mCherry, EYFP-*Mc*AP1μ and EYFP-*Mc*AP2μ.

**Supplementary Figure 5.** EYFP-*Mc*AP1μ and EYFP-*Mc*AP2μ co-localization with FM4-64.

**Supplementary Fig 6.** EYFP-*Mc*AP1μ and EYFP-*Mc*AP2μ respond to the BFA in a heterologous expression system.

**Supplementary Figure 7.** Subcellular localization of *Mc*PIP1;4-mCherry and *Mc*PIP2;1-mCherry.

**Supplementary Figure 8.** The carboxyl terminus of the *M. crystallinum* PIPs have the APµ binding motif.

**Supplementary Figure 9.** Molecular weight shift of McPIP2;1 in PM fractions

**Supplementary Table S1.** Proteins identified from the PCM fractions analyzed by LC-MS/MS from *M. crystallinum* root.

**Supplementary Table S2.** Proteins identified by LC-MS/MS in each PCM fraction isolate by FFZE from *M. crystallinum* root.

**Supplementary Table S3.** Salt up-regulated proteins in each PCM fractions from *M. crystallinum* root.

Supplementary Material and Methods Table. Oligonucleotides use for gene cloning.

## ACKNOWLEDGMENTS

We dedicate this research to the long scientific career of Rosario Vera Estrella and her interest in aquaporins; it would not have been possible without your research and support. We thank Dr. Denis Faubert at the Institut de Recherches Cliniques de Montréal-Proteomics Discovery Platform for LC-MS/MS analysis. We want to thank to Laboratorio Nacional de Microscopia Avanzada (LNMA, IBT-UNAM) by technical support with confocal microscopy. Also, technical help from Maria Guadalupe Muñoz Garcia is recognized. G-M was the recipient of a scholarship from CONACyT-Mexico.

## REFERENCES

1. Adams P, Nelson DE, Yamada S, Chmara W, Jensen RG, Bohnert HJ and Griffiths H (1998). Growth and development of Mesembryanthemum crystallinum (Aizoaceae). The New Phytologist 138: 171–190 DOI: 10.1046/j.1469-8137.1998.00111.x.

2. Afzal Z, Howton TC, Sun Y and Makhtar MS (2016). The roles of Aquaporins in plant stress responses. Journal of Developmental Biology 4: 9 DOI: 10.3390/jdb4010009 4:9.

3. Alexa A, Rahnenfuhrer J (2021). topGO: Enrichment Analysis for Gene Ontology. R package version 2.46.0.

4. Alexa A, Rahnenführer J and Lengauer T (2006). Improved scoring of functional groups from gene expression data by decorrelating GO graph structure. Bioinformatics 22: 1600–1607 DOI: 10.1093/bioinformatics/btl140.

5. Alexandersson E, Fraysse L, Sjövall-Larsen S, Gustavsson S, Fellert M, Karlsson M, Johanson U and Kjellbom P (2005). Whole gene family expression and drought stress regulation of aquaporins. Plant Molecular Biology 59: 469–484 DOI: 10.1007/s11103-005-0352-1.

6. Amezcua-Romero JC, Pantoja O and Vera-Estrella R (2010). Ser123 is essential for the water channel activity of McPIP2;1 from *Mesembryanthemum crystallinum*. Journal of Biological Chemistry 285: 16739–16747 DOI: 10.1074/jbc.M109.053850.

7. Aroca R, Porcel R and Ruiz-Lozano JM (2012). Regulation of root water uptake under abiotic stress conditions. Journal of Experimental Botany 63: 43–57 DOI: 10.1093/jxb/err266.

8. Aroca R, Tognoni F, Irigoyen JJ, Sanchez DM and Pardossi A (2001). Different root low temperature response of two maize genotypes differing in chilling sensitivity. Plant Physiology and Biochemistry 39: 1067–1073 DOI: 10.1016/S0981-9428(01)01335-3.

9. Arora D and Van Damme D (2021). Motif-based endomembrane trafficking. Plant Physiology 186: 221–38 DOI: 10.1093/plphys/kiab077.

10. Asaoka R, Uemura T, Nishida S, Fujiwara T, Ueda T and Nakano A (2013). New insights into the role of Arabidopsis RABA1 GTPases in salinity stress tolerance. Plant signaling & behavior 8:9 e25377 DOI: 10.4161/psb.25377.

11. Baral A, Shruthi KS and Mathew MK (2015). Vesicular trafficking and salinity responses in plants. IUBMB Life 67: 677–686 DOI: 10.1002/iub.1425.

12. Barkla B (2018). Free flow zonal electrophoresis for fractionation of plant membrane compartments prior to proteomic analysis. Edited by Mock HP Matros A, Witzel K in Plant Membrane Proteomics. Methods in Molecular Biology vol, 1696. Humana Press, New York, NY DOI: 10.1007/978-1-4939-7411-5_1.

13. Barkla BJ, Vera-Estrella R and Pantoja O (2007). Enhanced separation of membranes during Free Flow Zonal Electrophoresis in plants. Analytical Biochemistry 15: 5181–5187 DOI: 10.1021/ac070159v.

14. Bienert MD, Diehn TA, Richet N, Chaumont F and Bienert GP (2018). Heterotetramerization of Plant PIP1 and PIP2 aquaporins is an evolutionary ancient feature to guide PIP1 plasma Membrane localization and function. Frontiers in Plant Science 26: 382 DOI: 10.3389/fpls.2018.00382.

15. Binns D, Dimmer E, Huntley R, Barrell D, O’Donovan C and Apweiler R (2009). QuickGO: a web-based tool for Gene Ontology searching. Bioinformatics 25: 3045–3046 DOI: 10.1093/bioinformatics/btp536.

16. Boursiac Y, Boudet J, Postaire O, Luu D.T, Tournaire-Roux C and Maurel C (2008). Stimulus-induced downregulation of root water transport involves reactive oxygen species-activated cell signalling and plasma membrane intrinsic protein internalization. The Plant Journal 56: 207–218 DOI: 10.1111/j.1365-313X.2008.03594.x.

17. Boursiac Y, Chen S, Luu DT, Sorieul M, van den Dries N and Maurel C (2005). Early effects of salinity on water transport in *Arabidopsis* roots. Molecular and cellular features of aquaporin expression. Plant Physiology 139: 790–805 DOI: 10.1104/pp.105.065029.

18. Bradford MM (1976). A rapid and sensitive method for the quantitation of microgram quantities of protein utilizing the principle of protein-dye binding. Analytical Biochemistry 7: 248–54 DOI: 10.1006/abio.1976.9999.

19. Chaumont F and Tyerman SD (2014). Aquaporins: highly regulated channels controlling plant water relations. Plant Physiology 164:1600–1618 DOI 10.1104/pp.113.233791.

20. Chaumont F, Barrieu F, Jung R and Chrispeels MJ (2000). Plasma membrane intrinsic proteins from maize cluster in two sequence subgroups with differential aquaporin activity. Plant Physiology 122: 1025–1034 DOI: 10.1104/pp.122.4.1025.

21. Chaumont F, Barrieu F, Wojcik E, Chrispeels MJ and Jung R (2001). Aquaporins constitute a large and highly divergent protein family in maize. Plant Physiology 125: 1206–1215 DOI: 10.1104/pp.125.3.1206.

22. Chen T, Cai X, Wu X, Karahara I, Schreiber L and Ling J (2011). Casparian strip development and its potential function in salt tolerance. Plant Signaling & Behaviour 6:1499–1502 DOI: 10.4161/psb.6.10.17054.

23. Cheng X, Lang I, Adeniji OS and Griffing L (2017). Plasmolysis-deplasmolysis causes changes in endoplasmic reticulum form, movement, flow, and cytoskeletal association. Journal of Experimental Botany 168: 4075–4087 DOI: 10.1093/jxb/erx243.

24. Chevalier AS and Chaumont F (2015). Trafficking of plant plasma membrane aquaporins: multiple regulation levels and complex sorting signals. Plant and Cell Physiology 56: 819–829 DOI: 10.1093/pcp/pcu203.

25. Conesa A and Götz S (2008). Blast2GO: A comprehensive suite for functional analysis in plant genomics. International Journal of Plant Genomics 2008: 619832 DOI: 10.1155/2008/619832.

26. Conesa A, Götz S, García-Gómez JM, Terol J, Talón M and Robles M (2005). Blast2GO: a universal tool for annotation, visualization and analysis in functional genomics research. Bioinformatics 21: 3674–3676 DOI: 10.1093/bioinformatics/bti610.

27. Danielson JÅ and Johanson U (2010). Phylogeny of major intrinsic proteins. Edited by JahnTP and Bienert GP In *MIPs and Their Role in the Exchange of Metalloids*, Advances in Experimental Medicine and Biology, vol 679, Springer DOI: 10.1007/978-1-4419-6315-4_2.

28. Dhonukshe P, Aniento F, Hwang I, Robinson DG, Mravec J, Stierhof YD and Friml J (2007). Clathrin-mediated constitutive endocytosis of PIN auxin efflux carriers in *Arabidopsis*. Current Biology 17: 520–527 DOI: 10.1016/j.cub.2007.01.052.

29. Dhugga KS, Tiwari SC and Ray PM (1997). A reversibly glycosylated polypeptide (RGP1) possibly involved in plant cell wall synthesis: purification, gene cloning, and trans-Golgi localization. Proceedings of the National Academy of Sciences of the United States of America 94: 7679–7684 DOI: 10.1073/pnas.94.14.7679.

30. Di Pietro M, Vialaret J, Li GW, Hem S, Prado K, Rossignol M, Maurel C and Santoni V (2013). Coordinated post-translational responses of aquaporins to abiotic and nutritional stimuli in *Arabidopsis* roots. Molecular & Cellular Proteomics 12: 3886–3897 DOI: 10.1074/mcp.M113.028241.

31. Dietz KJ and Arbinger B (1996). cDNA sequence and expression of subunit E of the vacuolar H(+)-ATPase in the inducible Crassulacean acid metabolism plant *Mesembryanthemum crystallinum*. Biochemica et Biophysica Acta 1281: 134–138. DOI: 10.1016/0005-2736(96)00044-2.

32. Fetter K, Van Wilder V, Moshelion M and Chaumont F (2004). Interactions between plasma membrane aquaporins modulate their water channel activity. The Plant cell 16: 215–228 DOI: 10.1105/tpc.017194.

33. French AP, Mills S, Swarup R, Bennett MJ and Pridmore TP (2008). Colocalization of fluorescent markers in confocal microscope images of plant cells. Nature Protocols 3: 619–628 DOI: 10.1038/nprot.2.

34. Gambetta GA, Knipfer T, Fricke W and McElrone AJ (2017). Aquaporins and root water uptake. Edited by Chaumont F and Tyerman SD In *Plant Aquaporins, Signaling and Communications in Plants*, Spriger, DOI: 10.1007/978-3-319-49395-4_6.

35. Hacke UG and Laur J (2016). Aquaporins: Channels for the molecule of life. Edited by Chichester In eLS. John Wiley & Sons, DOI: 10.1002/9780470015902.a0001289.pub2.

36. Heidrich HG and Hannig K (1989). Separation of cell populations by free-flow electrophoresis. Methods in Enzymology 171:513–531 DOI: 10.1016/s0076-6879(89)71028-4.

37. Heymann JB and Engel A (1999). Aquaporins: Phylogeny, Structure, and Physiology of Water Channels. News in Physiological Sciences 14: 187–193 DOI: 10.1152/physiologyonline.1999.14.5.187.

38. Hoagland DR and Arnon DI (1938). The water culture method for growing plants without soil. Californian Experimental Station Circular 347: 1–39.

39. Javot H and Maurel C (2002). The role of aquaporins in root water uptake. Annuals of Botany 90: 301–313 DOI: 10.1093/aob/mcf199.

40. Kirch HH, Vera-Estrella R, Golldack D, Quigley F, Michalowski CB, Barkla BJ and Bohnert HJ (2000). Expression of water channel proteins in *Mesembryanthemum crystallinum*. Plant Physiology 123: 111–124 DOI: 10.1104/pp.123.1.111.

41. Kumar S, Stecher G, Li M, Knyaz C and Tamura K (2018). MEGA X: Molecular Evolutionary Genetics Analysis across Computing Platforms. Molecular Biology and Evolution 35: 1547–1549 DOI: 10.1093/molbev/msy096.

42. Laemmli UK (1970). Cleavage of structural proteins during the assembly of the head of bacteriophage T4. Nature 227: 680–685 DOI: 10.1038/227680a0.

43. Lalonde S, Sero A, Pratelli R, Pilot G, Chen J, Sardi MI, Parsa SA, Kim DY, Acharya BR, Stein EV, Hu HC, Villiers F, Takeda K, Yang Y, Han YS, Schwacke R, Chiang W, Kato N, Loqué D, Assmann SM, Kwak JM, Schroeder JI, Rhee SY and Frommer WB (2010). A membrane protein/signaling protein interaction network for Arabidopsis version AMPv2. Frontiers in Physiology 1:24 DOI: 10.3389/fphys.2010.00024.

44. Langhans M, Förster S, Helmchen G and Robinson DG (2011). Differential effects of the brefeldin A analogue (6R)-hydroxy-BFA in tobacco and *Arabidopsis*. Journal of Experimental Botany 62: 2949–2957 DOI: 10.1093/jxb/err007.

45. Le SQ and Gascuel O (2008). An improved general amino acid replacement matrix. Molecular Biology and Evolution. 25: 1307–1320 DOI: 10.1093/molbev/msn067.

46. Lee SH and Zwiazek JJ (2015). Regulation of aquaporin-mediated water transport in Arabidopsis roots exposed to NaCl. Plant and Cell Physiology 56: 750–8 DOI: 10.1093/pcp/pcv003.

47. Luu DT, Martinière A, Sorieul M, Runions J and Maurel C (2012). Fluorescence recovery after photobleaching reveals high cycling dynamics of plasma membrane aquaporins in *Arabidopsis* roots under salt stress. The Plant Journal 69: 894–905 DOI: 10.1111/j.1365-313X.2011.04841.x.

48. Madrid R, Le Maout S, Barrault MB, Janvier K, Benichou S and Mérot J (2001). Polarized trafficking and surface expression of the AQP4 water channel are coordinated by serial and regulated interactions with different clathrin-adaptor complexes. The EMBO Journal 20: 7008–7021 DOI: 10.1093/emboj/20.24.7008.

49. Martinière A, Li X, Runions J, Lin J, Maurel C and Luu DT (2012). Salt stress triggers enhanced cycling of *Arabidopsis* root plasma-membrane aquaporins. Plant Signaling & Behavior 7: 529–32 DOI: 10.4161/psb.19350.

50. Moshelion M, Becker D, Biela A, Uehlein N, Hedrich R, Otto B, Levi H, Moran N and Kaldenhoff B (2002). Plasma membrane aquaporins in the motor cells of *Samanea saman*: diurnal and circadian regulation. Plant cell 14: 727–739 DOI: 10.1105/tpc.010351.

51. Munns R (2002). Comparative physiology of salt and water stress. *Plant*, Cell & Environment 25: 239–250 DOI: 10.1046/j.0016-8025.2001.00808.x.

52. Muries B, Faize M, Carvajal M and Martínez-Ballesta Mdel C (2011). Identification and differential induction of the expression of aquaporins by salinity in broccoli plants. Molecular Biosystem 4:1322–35 DOI:10.1039/c0mb00285b.

53. Nelson DE, Glaunsinger B and Bohnert HJ (1997). Abundant accumulation of the calcium-binding molecular chaperone calreticulin in specific floral tissues of *Arabidopsis thaliana*. Plant Physiology 114: 29–37 DOI: 10.1104/pp.114.1.29.

54. Obrdlik P, El-Bakkoury M, Hamacher T, Cappellaro C, Vilarino C, Fleischer C, Ellerbrok H, Kamuzinzi R, Ledent V, Blaudez D, Sanders D, Revuelta JL, Boles E, André B and Frommer WB (2004). K+ channel interactions detected by a genetic system optimized for systematic studies of membrane protein interactions. Proceedings of the National Academy of Sciences of the United States of America 101: 12242–12247 DOI: 10.1073/pnas.0404467101.

55. Oh DH, Barkla BJ, Vera-Estrella R, Pantoja O, Lee SY, Bohnert HJ and Dassanayake M (2015). Cell type-specific responses to salinity - the epidermal bladder cell transcriptome of *Mesembryanthemum crystallinum*. New Phytologist 207:627–644 DOI: 10.1111/nph.13414.

56. Oparka KJ (1994). Plasmolysis: new insights into an old process. New Phytologist 126: 571– 591 DOI: 10.1111/j.1469-8137.1994.tb02952.x.

57. Ozu M, Galizia L, Acuña C and Amodeo G (2018). Aquaporins: More than functional monomers in a tetrameric arrangement. Cells 11: 209 DOI: 10.3390/cells7110209.

58. Pardo JM and Serrano R (1989). Structure of a plasma membrane H+-ATPase gene from the plant *Arabidopsis thaliana*. Journal of Biological Chemistry 25: 8557–8562.

59. Park M, Song K, Reichardt I, Kim H, Mayer U, Stierhof YD, Hwang I and Jürgens G (2013). Arabidopsis μ-adaptin subunit AP1M of adaptor protein complex 1 mediates late secretory and vacuolar traffic and is required for growth. Proceedings of the National Academy of Sciences of the United States of America 18:10318–10323 DOI: 10.1073/pnas.1300460110.

60. Perez-Silva JG, Araujo-Voces M and Quesada V (2018). nVenn: generalized, quasi-proportional Venn and Euler diagrams. Bioinformatics 34: 2322–2324, DOI: 10.1093/bioinformatics/bty109.

61. Prak S, Hem S, Boudet J, Viennois G, Sommerer N, Rossignol M, Maurel C and Santoni V (2008). Multiple phosphorylations in the C-terminal tail of plant plasma membrane aquaporins: role in subcellular trafficking of AtPIP2;1 in response to salt stress. Molecular & Cellular Proteomics 7: 1019–1030 DOI: 10.1074/mcp.M700566-MCP200.

62. Prado K, Boursiac Y, Tournaire-Roux C, Monneuse JM, Postaire O, Da Ines O, Schäffner AR, Hem S, Santoni V, Maurel C (2013). Regulation of Arabidopsis leaf hydraulics involves light-dependent phosphorylation of Aquaporins in veins. The Plant Cell 25: 1029–1039 DOI:10.1105/tpc.112.108456.

63. Preston GM, Carroll TP, Guggino WB and Agre P (1992). Appearance of water channels in *Xenopus* oocytes expressing red cell CHIP28 protein. Science 256: 385–387 DOI: 10.1126/science.256.5055.385. PMID: 1373524.

64. Reynolds GD, August B and Bednarek SY (2014). Preparation of enriched plant clathrin-coated vesicles by differential and density gradient centrifugation. Editor Otegui M in *Plant Endosomes*. Methods in Molecular Biology, vol 1209. Humana Press, New York, NY DOI: 10.1007/978-1-4939-1420-3_13.

65. Rigal A, Doyle SM and Robert S (2015). Live cell imaging of FM4-64, a tool for tracing the endocytic pathways in *Arabidopsis* root cells. Edited by Estevez J. In Plant Cell Expansion. Humana Press, New York, NY. Methods in Molecular Biology 1242: 93–103 DOI: 10.1007/978-1-4939-1902-4_9.

66. Robinson DG, Langhans M, Saint-Jore-Dupas C and Hawes C (2008). BFA effects are tissue and not just plant specific. Trends in Plant Science 13: 405–408 DOI: 10.1016/j.tplants.2008.05.010.

67. Rosas-Santiago P, Lagunas-Gómez D, Barkla BJ, Vera-Estrella R, Lalonde S, Jones A, Frommer WB, Zimmermannova O, Sychrová H and Pantoja O (2015). Identification of rice cornichon as a possible cargo receptor for the Golgi-localized sodium transporter OsHKT1;3. Journal of Experimental Botany 66:2733–2748 DOI: 10.1093/jxb/erv069.

68. Sanchez-Romera B and Aroca R (2020). Plant roots-The hidden half for investigating salt and drought stress responses and tolerance. Edited by Hasanuzzaman M and Tanveer M *In Salt and drought stress tolerance in plants*, Springer DOI: 10.1007/978-3-030-40277-8.

69. Sander H, Wallace S, Plouse R, Tiwari S and Gomes AV (2019). Ponceau S staining for total protein normalization. Analytical Biochemistry 15: 44–53 DOI: 10.1016/j.ab.2019.03.010.

70. Schindelin J, Arganda-Carreras I, Frise E, Kaynig V, Longair M, Pietzsch T, Preibisch S, Rueden C, Saalfeld S, Schmid B, Tinevez JY, White DJ, Hartenstein V, Eliceiri K, Tomancak P, Cardona A (2012). Fiji: an open-source platform for biological-image analysis. Nature Methods Jun 9: 676–82 DOI: 10.1038/nmeth.2019.

71. Shenghao Q, Yiyang L, Yan H, Shubo W, Geng C and Guowei L (2019). Aquaporins and their function in root water transport under salt stress conditions in *Eutrema salsugineum*. Plant Science 287: 110199 DOI: 10.1016/j.plantsci.2019.110199.

72. Smith MS, Baker M, Halebian M and Smith JC (2017). Weak molecular interactions in Clathrin-mediated endocytosis. Frontiers in Molecular Biosciences 4: 72- DOI: 10.3389/fmolb.2017.00072.

73. Suga S and Maeshima M (2004). Water channel activity of radish plasma membrane aquaporins heterologously expressed in yeast and their modification by site-directed mutagenesis. Plant and Cell Physiology 45: 823–830 DOI: 10.1093/pcp/pch120.

74. Suku S, Knipfer T and Fricke W (2013). Do root hydraulic properties change during the early vegetative stage of plant development in barley (*Hordeum vulgare*)?. Annals of Botany 113: 385–402 DOI: 10.1093/aob/mct270.

75. Supek F, Bošnjak M, Škunca N and Šmuc T (2011). REVIGO summarizes and visualizes long lists of gene ontology terms. PLoS One 6: e21800 DOI: 10.1371/journal.pone.0021800.

76. Sutka M, Li G, Boursiac Y, Doumas P and Maurel C (2011). Natural variation of root hydraulics in *Arabidopsis* growth in normal and salt-stressed conditions. Plant Physiology 155: 1264–1276 DOI: 10.1104/pp.110.163113.

77. Teh OK, Shimono Y, Shirakawa M, Fukao Y, Tamura K, Shimada T and Hara-Nishimura I (2013). The AP-1 μ adaptin is required for KNOLLE localization at the cell plate to mediate cytokinesis in *Arabidopsis*. Plant Cell and Physiology 54:838–847 DOI: 10.1093/pcp/pct048.

78. Temmei Y, Uchida S, Hoshino D, Kanzawa N, Kuwahara M, Sasaki S and Tsuchiya T (2005). Water channel activities of *Mimosa pudica* plasma membrane intrinsic proteins are regulated by direct interaction and phosphorylation. FEBS Lett. 15:4417–4422 DOI: 10.1016/j.febslet.2005.06.082.

79. Ueda M, Tsutsumi N, Fujimoto M (2016). Salt stress induces internalization of plasma membrane aquaporin into the vacuole in *Arabidopsis thaliana*. Biochemical and Biophysical Research Communications 10:742–746 DOI: 10.1016/j.bbrc.2016.05.028.

80. Vaziriyeganeh M, Lee SH, Zwiazek JJ (2018). Water transport properties of root cells contribute to salt tolerance in halophytic grasses *Poa juncifolia* and *Puccinellia nuttalliana*. Plant Science 276: 54–62 DOI: 10.1016/j.plantsci.2018.08.00.

81. Vera-Estrella R, Barkla BJ, Amezcua-Romero JC and Pantoja O (2012). Day/night regulation of aquaporins during the CAM cycle in *Mesembryanthemum crystallinum*. Plant, Cell & Environment 35: 485–501 DOI: 10.1111/j.1365-3040.2011.02419.x.

82. Vera-Estrella R, Barkla B, Bohnert H and Pantoja O (1999). Salt stress in *Mesembryanthemum crystallinum* L. cell suspensions activates adaptive mechanisms similar to those observed in the whole plant. Planta 207:426–435. https://doi.org/10.1007/s004250050501.

83. Vera-Estrella R, Barkla BJ, Bohnert HJ and Pantoja O (2004). Novel regulation of aquaporins during osmotic stress. Plant Physiology 135: 2318–2329 DOI: 10.1104/pp.104.044891.

84. Verdoucq L, Rodrigues O, Martinière A, Luu DT and Maurel C (2014). Plant aquaporins on the move: reversible phosphorylation, lateral motion and cycling. Current Opinion in Plant Biology 22:101–107 DOI: 10.1016/j.pbi.2014.09.011.

85. Wang C, Yan X, Chen Q, Jiang N, Fu W, Ma B, Liu J, Li C, Bednarek SY and Pan J (2013). Clathrin light chains regulate clathrin-mediated trafficking, auxin signaling, and development in Arabidopsis. Plant Cell 25: 499–516 DOI: 10.1105/tpc.112.108373.

86. Wang JG, Li S, Zhao XY, Zhou LZ, Huang GQ, Feng C and Zhang Y (2013). HAPLESS13, the Arabidopsis μ1 adaptin, is essential for protein sorting at the trans-Golgi network/early endosome. Plant Physiology 162:1897–1910 DOI:10.1104/pp.113.221051.

87. Whelan S and Goldman N (2001). A general empirical model of protein evolution derived from multiple protein families using a maximum-likelihood approach. Molecular Biology and Evolution 18: 691–699 DOI: 10.1093/oxfordjournals.molbev.a003851.

88. Yamada S, Katsuhara M, Kelly WB, Michalowski CB and Bohnert HJ (1995). A family of transcripts encoding water channel proteins: tissue-specific expression in the common ice plant. The Plant Cell 7: 1129–1142 DOI: 10.1105/tpc.7.8.1129.

89. Yan X, Wang Y, Xu M, Dahhan DA, Liu C, Zhang Y, Lin J, Bednarek SY and Pan J (2021). Cross-Talk between Clathrin-dependent post-Golgi trafficking and clathrin-mediated endocytosis in *Arabidopsis* root cells. The Plant Cell 33: 3057–3075 DOI:10.1093/plcell/koab180.

90. Yaneff A, Sigaut L, Marquez M, Alleva K, Pietrasanta L and Amodeo G (2014). Heteromerization of PIP aquaporins affects their intrinsic permeability. Proceedings of the National Academy of Sciences of the United States of America 111: 231–236. DOI: 10.1073/pnas.1316537111.

91. Zaghdoud C, Mota-Cardenas C, Carvajal M, Muries B, Ferchichi A, Martinez-Ballesta MC (2013). Elevated CO2 alleviates negative effects of salinity on broccoli (*Brassica oleracea* L. var Italica) plants by modulating water balance through aquaporins abundance. Environmental and Experimental Botany 95: 15–24 DOI: 10.1016/j.envexpbot.2013.07.003.

92. Zelazny E, Borst JW, Muylaert M, Batoko H, Hemminga MA and Chamont F (2007). FRET imaging in living maize cells reveals that plasma membrane aquaporins interact to regulate their subcellular localization. Proceedings of the National Academy of Sciences of the United States of America 104:12359–12364 DOI: 10.1073/pnas.0701180104.

93. Zhang L, Xing J, and Lin J (2019). At the intersection of exocytosis and endocytosis in plants. New Phytologist 224: 1479–1489 DOI: 10.1111/nph.16018.

