## Supplemental Figures and Tables for "*Mesembryanthemum crystallinum* plasma membrane root aquaporins are regulated via clathrin-coated vesicles in response to salt stress"

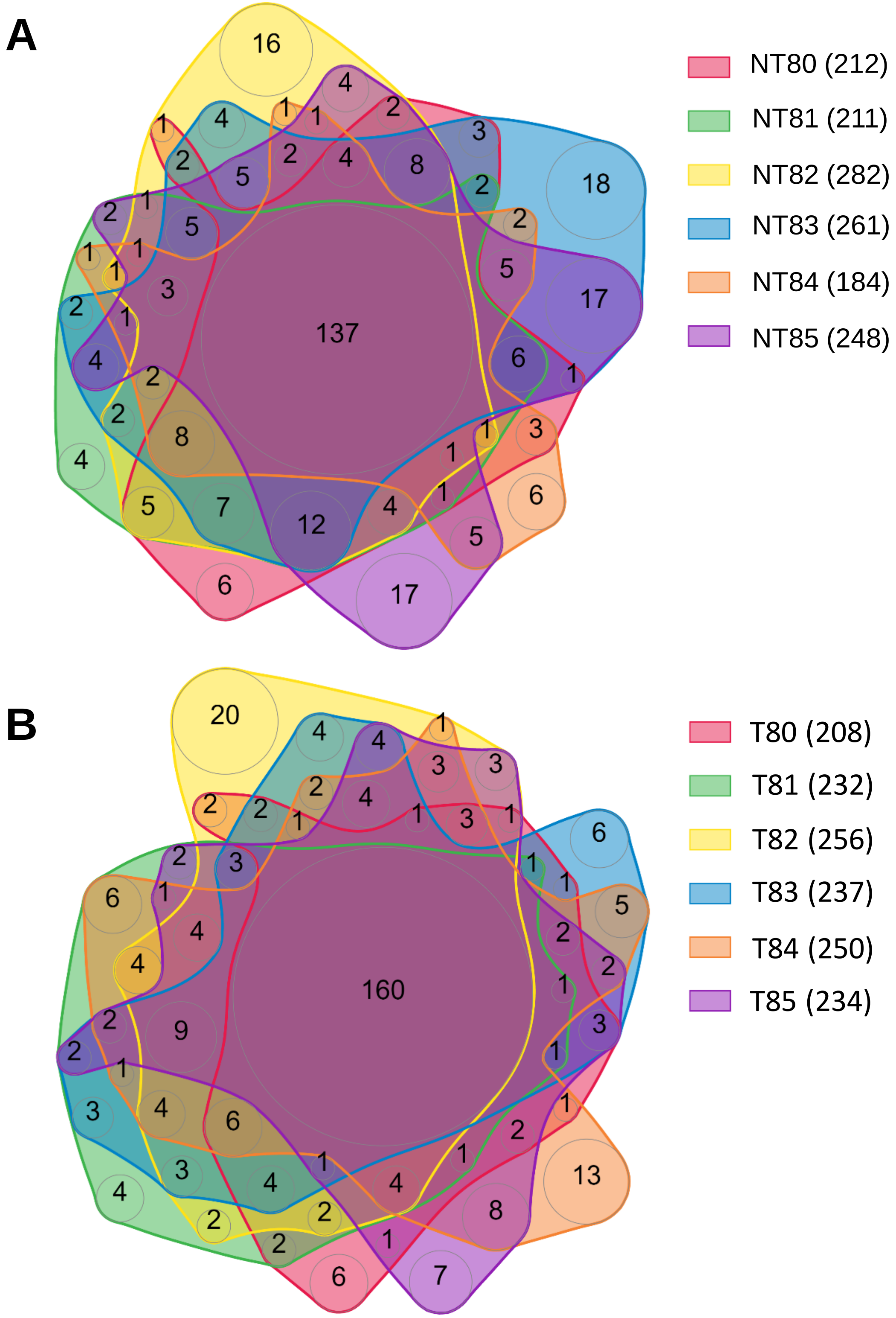


**Supplementary Fig 1. Positively-charge fractions in NT and T conditions shared a significant number of proteins.** nVenn diagram of shared and unique proteins identified by LC-MS/MS present in fractions 80 to 85 isolated by FFZE of ***M. crystallinum***. **A)** non-treated (NT) and **B)** salt-treated (T, 200 mM NaCl-7 d) root microsomes. Each fraction is represented with a color according to the legend. In brackets to the right of each legend is the total number of proteins identified for each fraction, delimited by the corresponding color-line. The intersections in the diagram show the number of shared proteins between fractions. Note that the central circle in the diagram is the number of shared proteins by all fractions.

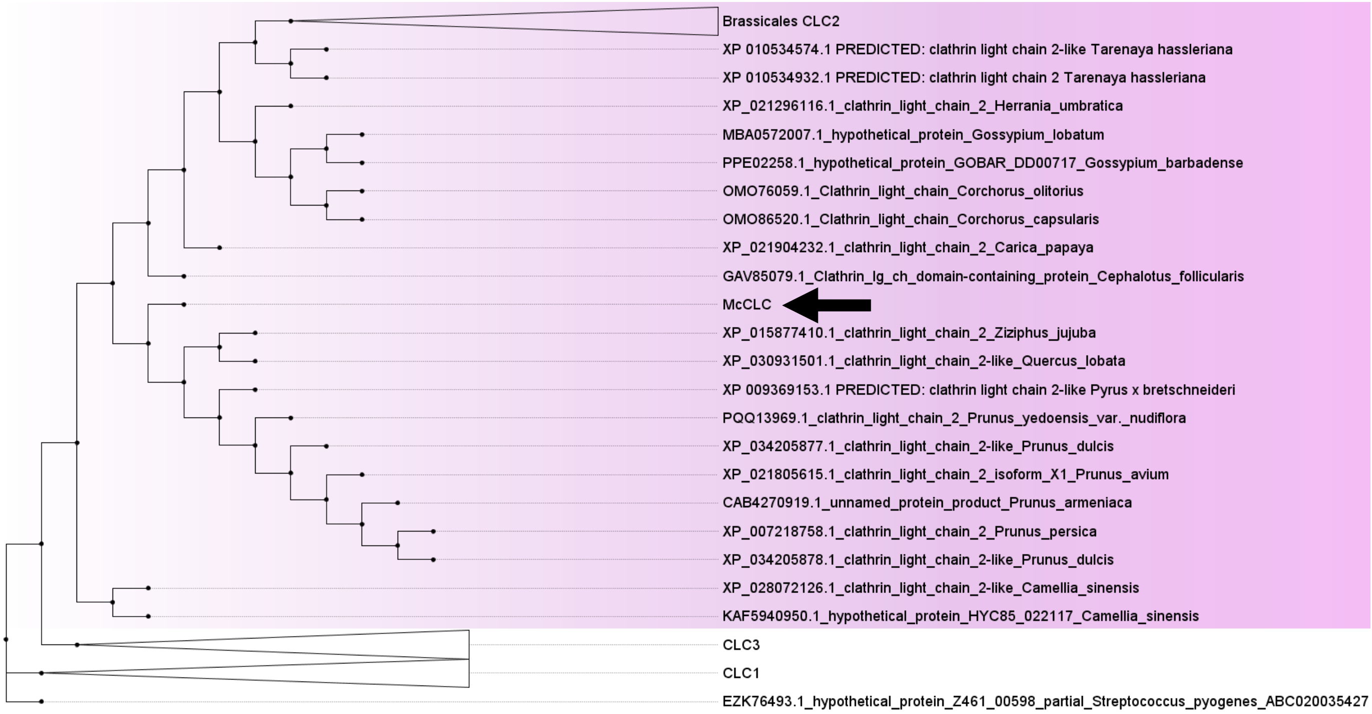


**Supplementary Fig 2. CLC is conserved in *M. crystallinum***. Phylogenetic tree of plant CLCs***.*** The evolutionary history was inferred by using the Maximum Likelihood method by Le & Gascuel., 2008. The tree with the highest log likelihood (-14641.66) is shown. Initial tree(s) for the heuristic search were obtained automatically by applying Neighbor-Join and BioNJ algorithms to a matrix of pairwise distances estimated using a JTT model, and then selecting the topology with superior log likelihood value. A discrete Gamma distribution was used to model evolutionary rate differences among sites (5 categories (+G, parameter = 2.5496)). The rate variation model allowed for some sites to be evolutionarily invariable ([+I], 2.15% sites). This analysis involved 79 amino acid sequences. All positions with less than 75% site coverage were eliminated, i.e., fewer than 25% alignment gaps, missing data, and ambiguous bases were allowed at any position (partial deletion option). There was a total of 418 positions in the final dataset. Evolutionary analyses were conducted in MEGA X (Kumar *et al*., 2018). Arrow indicates evolutionary position of *Mc*CLC in CLC2 gene subfamily marked with a light purple box.

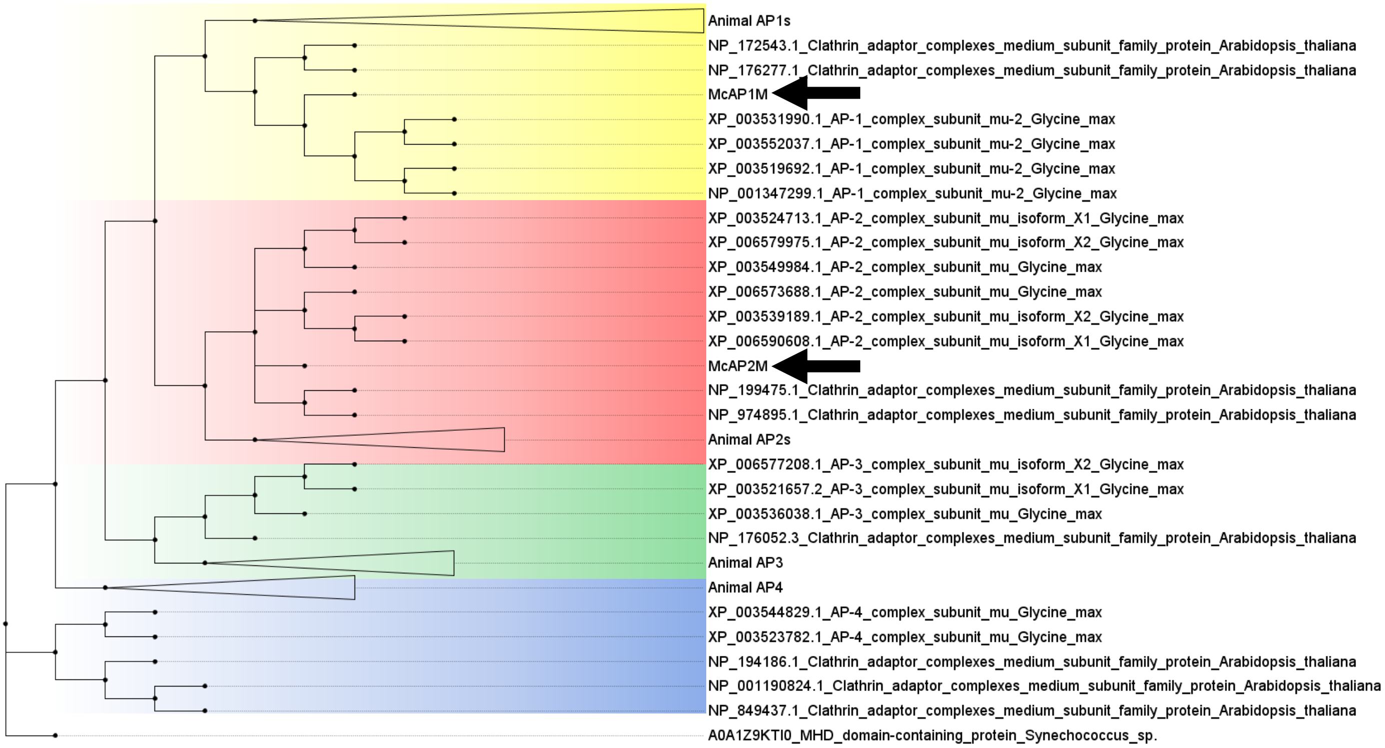


**Supplementary Fig 3. AP1μ and AP2μ are conserved in *M. crystallinum.*** Phylogenetic tree of APμs*.* The evolutionary history was inferred by using the Maximum Likelihood method by Whelan & Goldman., 2001. The tree with the highest log likelihood (-15452.87) is shown. Initial tree(s) for the heuristic search were obtained automatically by applying Neighbor-Join and BioNJ algorithms to a matrix of pairwise distances estimated using a JTT model, and then selecting the topology with superior log likelihood value. A discrete Gamma distribution was used to model evolutionary rate differences among sites (5 categories (+G, parameter = 0.7554)). This analysis involved 136 amino acid sequences. All positions with less than 75% site coverage were eliminated, i.e., fewer than 25% alignment gaps, missing data, and ambiguous bases were allowed at any position (partial deletion option). There was a total of 235 positions in the final dataset. Evolutionary analyses were conducted in MEGA X (Kumar et al., 2018). Arrows indicated evolutionary position of *Mc*AP1μ and *Mc*AP2μ in AP1μ or AP2μ gene subfamily marked with a yellow or red box, respectively.

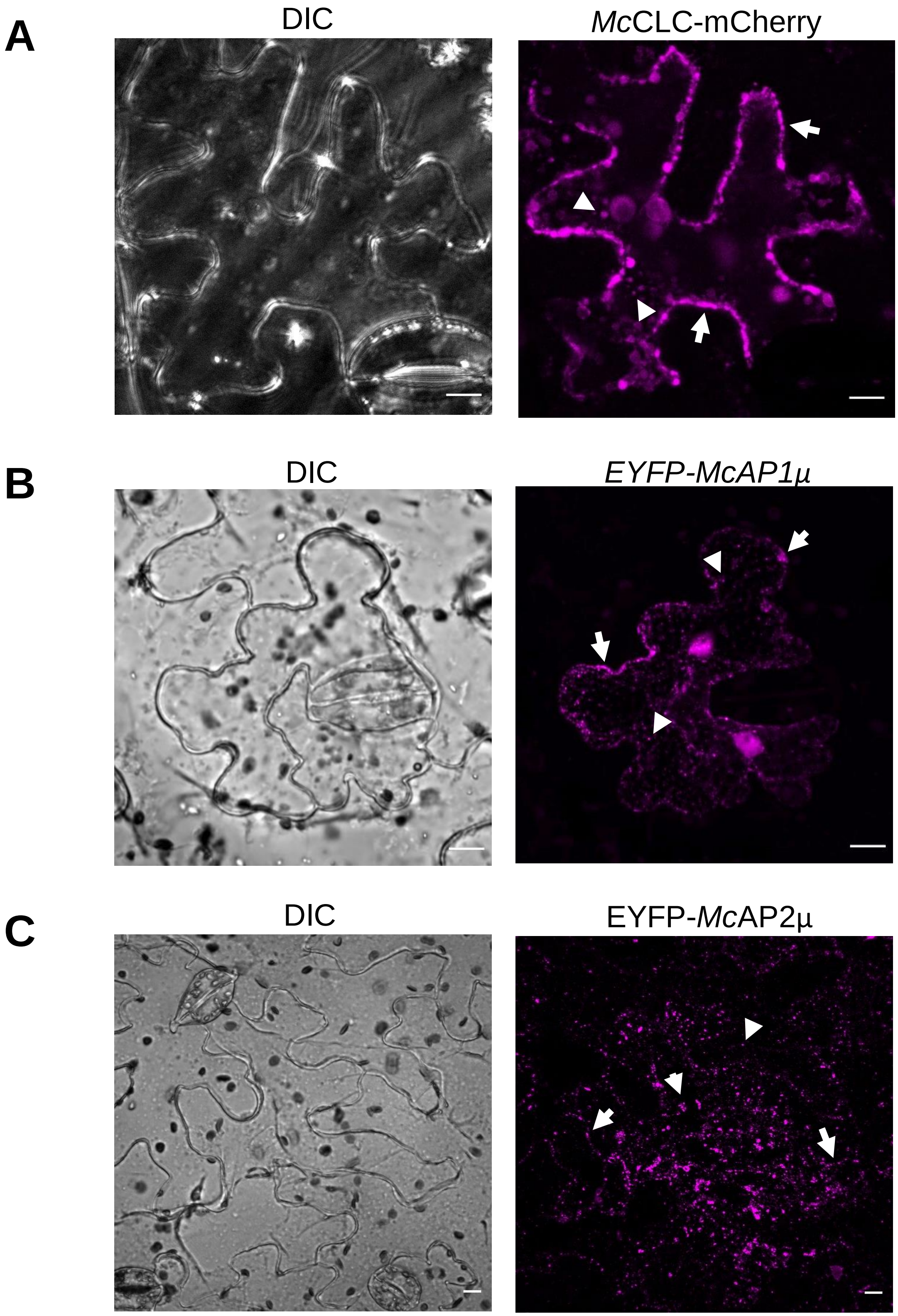


**Supplementary Fig 4. Intracellular localization of *Mc*CLC-mCherry*,* EYFP-*Mc*AP1µ and EYFP-*Mc*AP2µ**. Confocal images of abaxial epidermal peels of ***N. benthamiana*** leaves infiltrated with **A)** *McCLC*-mCherry; **B)** EYFP-*McAP1µ*; or **C)** EYFP-*McAP2µ*. Arrow and arrowheads indicates periphery of the cell or intracellular vesicles association, respectively. All images are representative of a Z-projection of at least three independent replicates *n*=3. DIC, bright field image. Scale bar 10 µM.

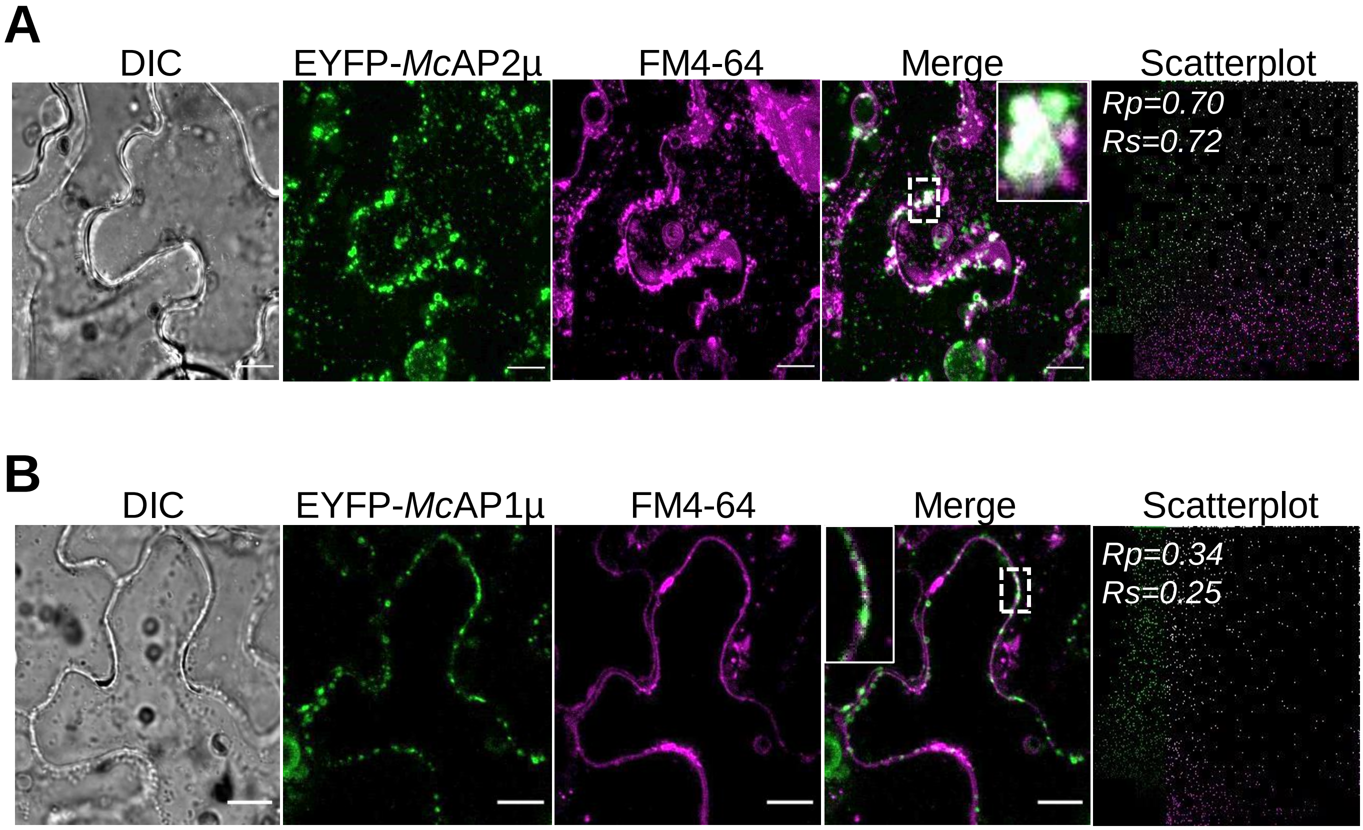


**Supplementary Fig 5**. **EYFP-*Mc*AP1µ and EYFP-*Mc*AP2µ co-localize with FM4-64**. Confocal images of abaxial epidermal peels from ***N. benthamiana*** leaves infiltrated with **A)** EYFP-*Mc*AP2µ or **B)** EYFP-*Mc*AP1µ. The colocalization analysis was performed using the PSC colocalization plug-in (French et al., 2008) in a region of interest (ROI) represented in the inset as areas of higher magnification. The obtained linear Pearson correlation coefficient (*Rp*) and the nonlinear Spearman rank correlation coefficient (*Rs*) are present in the resulting scatterplot. Values for *Rp* and *Rs* ≥ 0.5 indicate colocalization between both proteins. All images are representative of a Z-projection of ten images and least three independent replicates n=3. DIC, bright field image. Scale bar 10 µM.

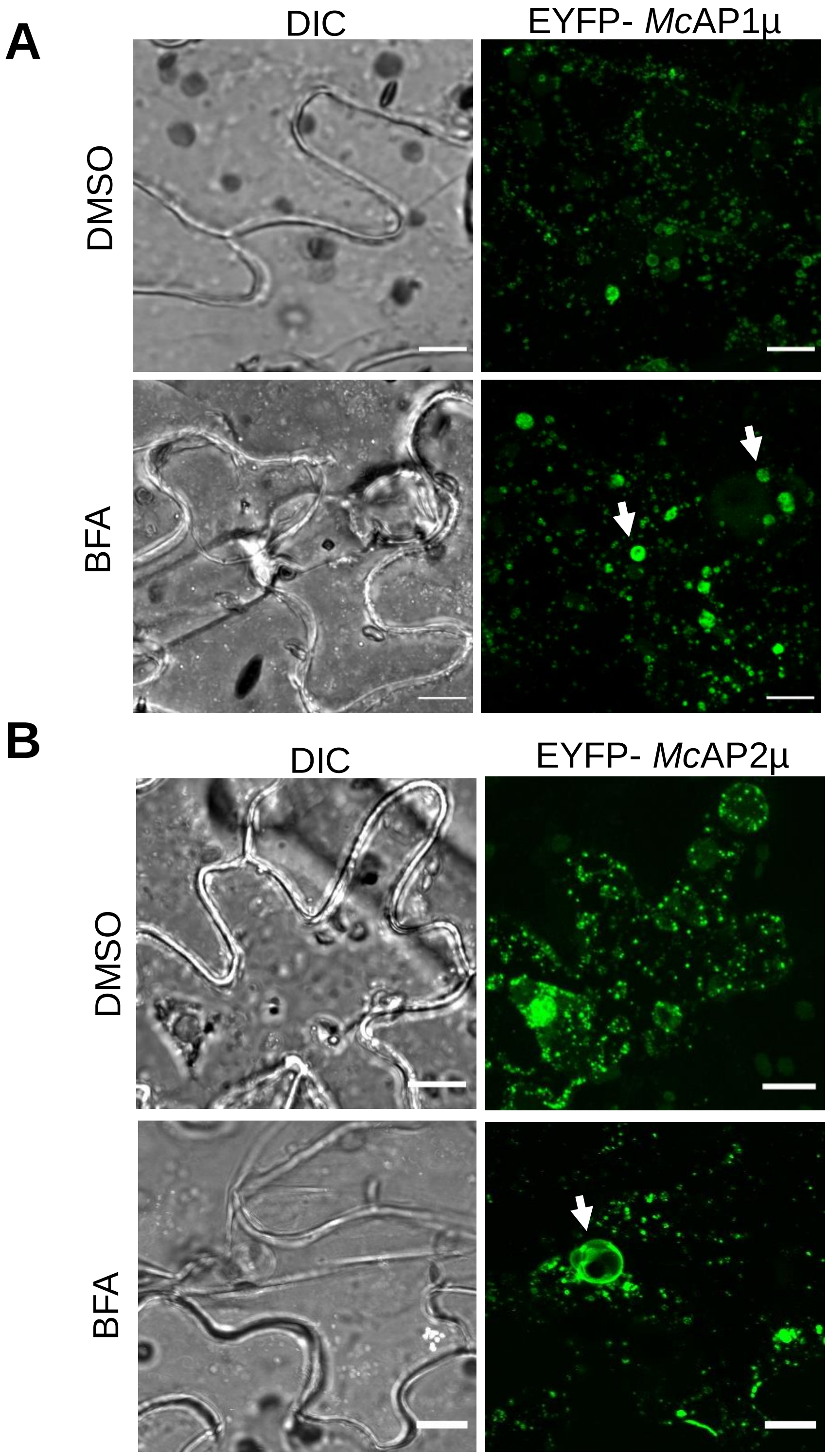


**Supplementary Fig 6**. **EYFP-*Mc*AP1µ and EYFP-*Mc*AP2µ respond to the BFA in a heterologous expression system.** Confocal images of abaxial epidermal peels from ***N. benthamiana*** leaves infiltrated with **A)** *EYFP-McAP1µ* or **B)** *EYFP-McAP2µ*. Agro-infiltrated leaves were treated with the fungal toxin BFA (90 µM for three hours) or DMSO (as a control) before imaging. Arrow indicates the presence of a BFA body. All images are representative of a Z-projection of ten images and are representative of at least three independent transformations. DIC, bright field image. Scale bar 10 µM.

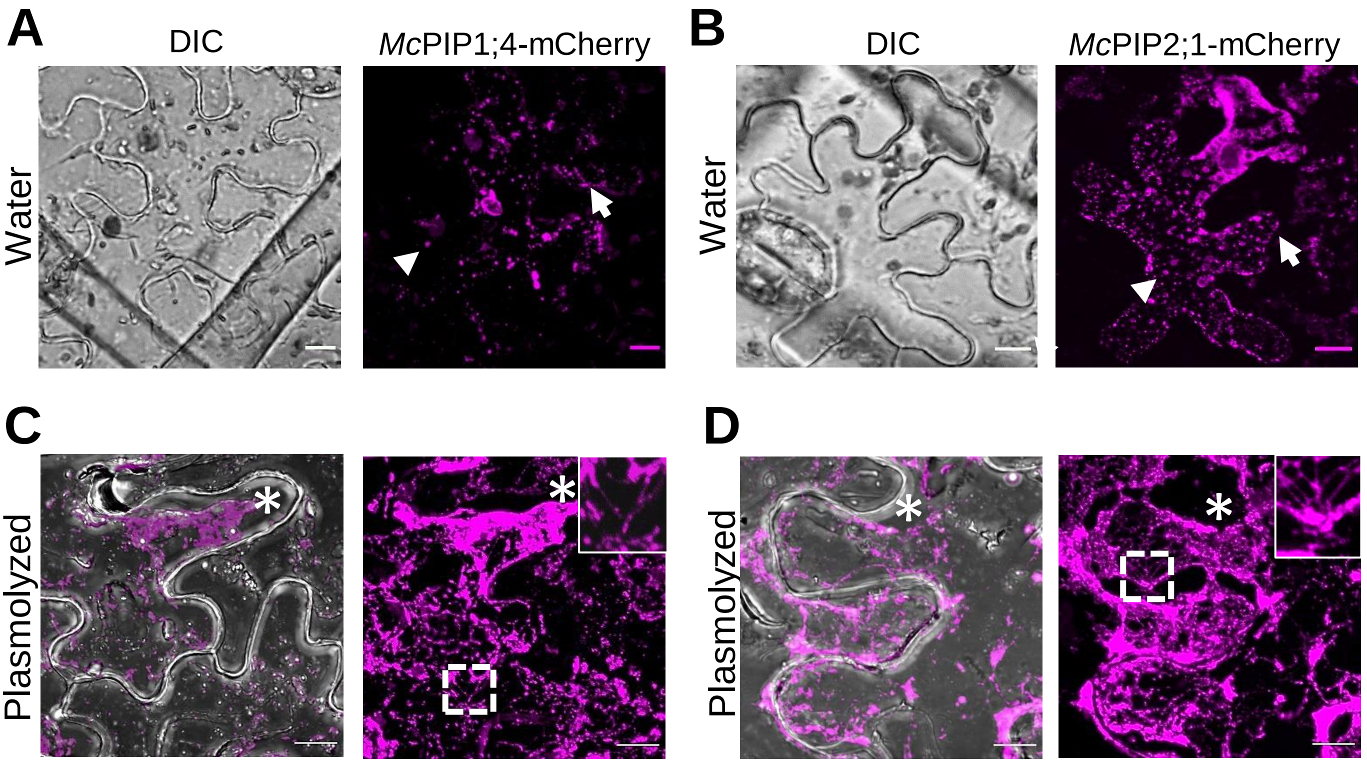


**Supplementary Fig 7. Subcellular localization of *McPIP1;4-mCherry* and *McPIP2;1-mCherry.*** Confocal images of abaxial epidermal peels from *N. benthamiana* leaves infiltrated with **A)**, **C)** *McPIP1;4-mCherry* or **B)**, **D)** *McPIP2;1-mCherry*. Arrow and arrowheads indicate -PM or intracellular vesicles, respectively in **A)** and **B)**. Transformed cells plasmolyzed with 4M NaCl show the presence of **C)** *McPIP1;4-mCherry* and **D)** *Mc*PIP2;1-mCherry signal in Hechtian strands. Inset boxes show areas of higher magnification in **C)** and **D)** with the presence of Hechtian strands. Asterisks in images show retraction of the protoplast away from the cell wall. All images are Z- projections and are representative of at least three independent replicates. DIC, bright field image. Scale bar 10 µM.

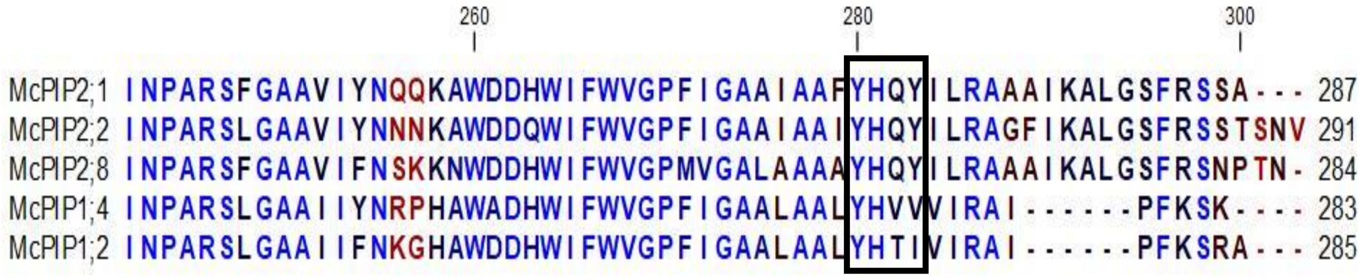


**Supplementary Fig 8**. **The carboxyl terminus of the *M. crystallinum* PIPs have the APµ binding motif.** Sequence alignment of the carboxyl terminal end of the *McPIP*s showing the presence of the YXXØ motif, enclosed in a box. The conserved residues are shown in blue. Alignment was performed in CLC-main Workbench 21.0.4

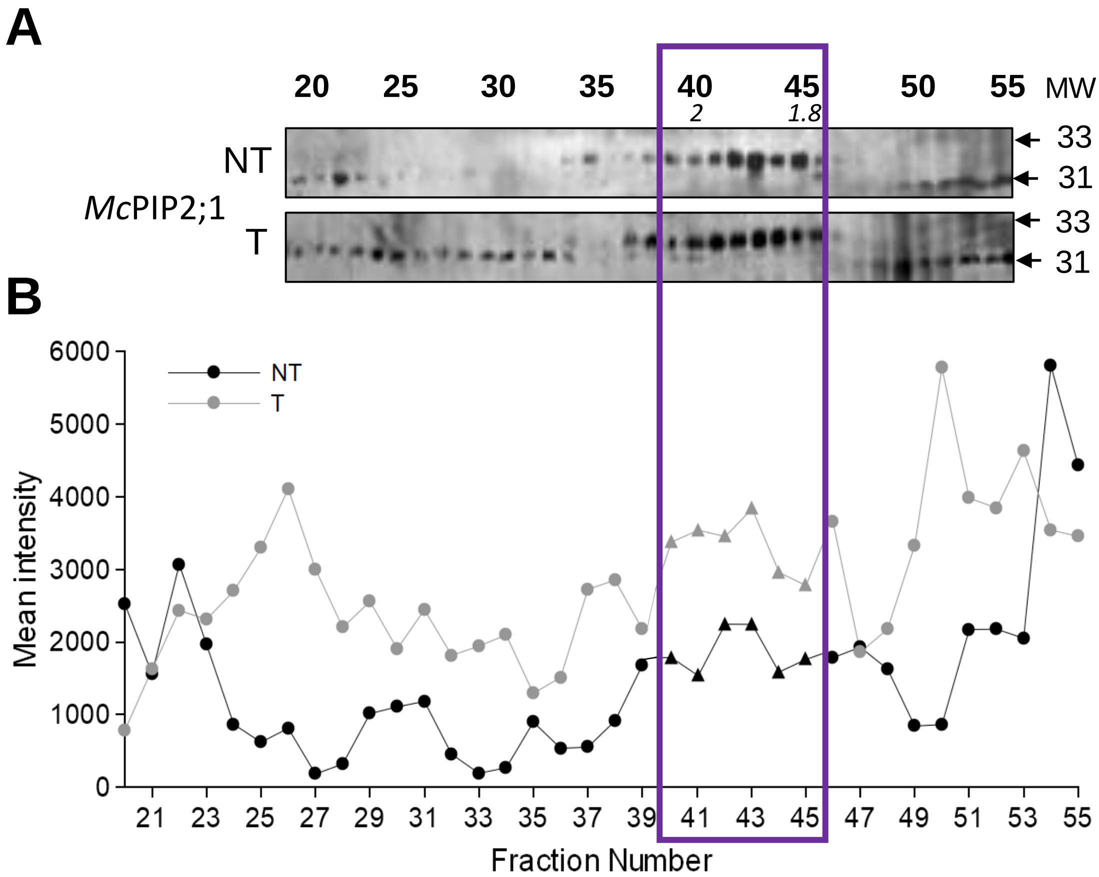


**Supplementary Fig 9. *Mc*PIP2;1 shows a molecular weight shift in PM fractions. A)** Immunological detection of *Mc*PIP2;1 in FFZE fractions enriched in: TP, PM and GA/ER (see Figure 1, for more detail) from non (NT) or salt-treated (T) *M. crystallinum* roots. The numbers in bold above the blots lane indicate the fraction number, and the number in italics show the difference in abundance (densitometry) between T/NT. The molecular weight (MW) of *McPIP2;1* is shown with arrow in the blots n=3. Notice how the 33 kDa isoform is only present in the PM fractions. **B)** Mean intensity of bands shown in **A)**, corresponding to the 31 kDa (circles) and 33 kDa (triangles) isoforms of *Mc*PIP2;1 from NT (black line) and T (gray line). PM enriched fractions are enclosed with a purple box in both panels.

***Table S1. Proteins identified from the PCM fractions analyzed by LC-MS/MS from M. crystallinum root.***

*Protein name and  accesion numer are display. Taxonomy colum show the species throught protein identification homology was carry out.*

*Molecular weight (MW) is show in kDa.*

*Cellular localization and biological process was obtein from  Blast2Go bioinformatics tool (version 3.0.11, Conesa et al., 2005 and 2008) and Quick-GO of EMBL-EBI (Binns et al., 2009).*

| Protein name | Uniprot Accession | Taxonomy | MW (kDa) | Cellular localization | Biological process |
|---|---|---|---|---|---|
| 14-3-3 protein | C1KG74 | *Vitis vinifera* | 29 | Cytosol/Vacuole/Nucleus/PM | Signaling |
| 14-3-3 protein 9 | P93214 | *Solanum lycopersicum* | 29 | Cytoplasm | Signaling |
| 14-3-3 protein SGF14h | E9KNA4 | *Glycine max* | 29 | PM/Chloroplast/Cyt/Vacuole | Signaling |
| 14-3-3-like protein | O65352 | *Helianthus annus* | 29 | Cytoplasm | Signaling |
| 14-3-3-like protein | P93259 | *Mesembryanthemum crystallinum* | 30 | Cytoplasm | Signaling |
| 14-3-3-like protein D | O49996 | *Nicotiana tabacum* | 28 | Cytoplasm | Signaling |
| 2,3-bisphosphoglycerate-independent phosphoglyc | Q42908 | *Mesembryanthemum crystallinum* | 61 | PM/Mitochondria/Cyt | Response to stress |
| 20S proteasome alpha subunit E2 | Q42134 | *Arabidopsis thaliana* | 26 | Proteasome | Proteolysis |
| 26S protease regulatory subunit 6A homolog A-like | A0A0A0LJT3 | *Cucumis sativus* | 47 | PM/Cyt | ATP catabolic process/Response to stress/Proteosome core assembly |
| 26S protease regulatory subunit S10B homolog B-like | A0A0A0LNP0 | *Cucumis sativus* | 45 | PM/Cyt/Cell wall/Proteasome | Proteolysis |
| 26S proteasome non-ATPase regulatory subunit 2 1A-like | UPI0002C338E5 | *Fragaria vesca* | 98 | Proteasome | Proteolysis |
| 26S proteasome non-ATPase regulatory subunit 4 homolog isoform X1 | I1J8K2 | *Glycine max* | 43 | Membrane/Cyt/Proteasome | Water transport/Response to stress/Golgi organization/Response to salt stress |
| 26S proteasome non-ATPase regulatory subunit 6 homolog | A0A6J5UYR1 | *Prunus mume* | 44 | Cyt/Proteasome | Root hair elongation/ER to Golgi vesicle mediated transort |
| 26S proteasome regulatory complex, Rpn2/Psmd1 subunit | A0A061ELR7 | *Theobroma cacao* | 110 | Proteasome | Proteasome assembly |
| 26S proteasome regulatory subunit 4 homolog A-like | A0A3Q7HYJ1 | *Solanum lycopersicum* | 49 | Cyt/Proteosome | Protein catabolic process |
| 40S ribosomal protein S11 | P17093 | *Glycine max* | 18 | Ribosome/Chloroplast | Translation |
| 40S ribosomal protein S13 | P62302 | *Glycine max* | 17 | Ribosome | Translation |
| 40S ribosomal protein S14-3 | A0A6P5TX25 | *Prunus avium* | 16 | PM/Ribosome | Lipid metabolic process/Translation |
| 40S ribosomal protein S23 | M7YGU5 | *Triticum urartu* | 19 | Ribosome | Translation |
| 40S ribosomal protein S27-1 | O64650 | *Arabidopsis thaliana* | 9 | Ribosome /PM/ Plasmodesmata | Translation |
| 40S ribosomal protein S3-3-like | A0A0A0LHQ6 | *Cucumis sativus* | 27 | Ribosome | Translation |
| 40S ribosomal protein S3a-like | A0A0A0L4P6 | *Cucumis sativus* | 30 | Cyt | Translation |
| 40S ribosomal protein S4-2-like | A0A1S2XMM1 | *Cicer arietinum* | 30 | Chloroplast/PM/Cyt | Translation |
| 40S ribosomal protein S7 | I1L2H4 | *Glycine max* | 23 | Plasmodesmata/PM/TP/Cell wall/Ch | Translation |
| 40S ribosomal protein S6-like | A0A0A0L5R5 | *Cucumis sativus* | 28 | Clorolasto/Cyt | Translation |
| 40S ribosomal protein S8 | Q08069 | *Zea mays* | 46 | Ribosome | Translation |
| 40S ribosomal protein S8 | UPI00046E0406 | *Prunus mume* | 25 | Chloroplast/Cyt | Translation |
| 40S ribosomal protein S9 | K7WJW2 | *Solanum tuberosum* | 23 | Cyt/nucleus/PM/Chloroplast/Cell wa | Translation |
| 40S ribosomal protein SA | B4G194 | *Zea mays* | 33 | Ribosome/ Cyt | Translation |
| 51 kDa subunit of complex I | A0A061E919 | *Theobroma cacao* | 53 | Mitochondria | Golgi vesicle transport |
| 5-methyltetrahydropteroyltriglutamate--homocysteine methyltransferase | P93263 | *Mesembryanthemum crystallinum* | 85 | Cyt/Plasmodesmata/TP/Golgi/PM | Response to salt stress/Response to stress |
| 60S acidic ribosomal protein P0 | P50346 | *Glycine max* | 34 | Ribosome | Translation |
| 60S ribosomal protein L1 | K7XKT4 | *Solanum tuberosum* | 45 | Ribosome | Translation |
| 60S ribosomal protein L11 | Q0DK10 | *Oryza sativa* | 21 | Ribosome | Translation |
| 60S ribosomal protein L12-like | UPI0002C3142A | *Fragaria vesca* | 18 | Ribosome | Translation |
| 60S ribosomal protein L15-like | UPI0002767146 | *Solanum lycopersicum* | 24 | PM/Ribosome | Translation |

| | | | | | |
|---|---|---|---|---|---|
| 60S ribosomal protein L17 | B6SJV1 | *Zea mays* | 18 | Ribosome | Translation |
| 60S ribosomal protein L18-like | A0A3Q7I5W4 | *Solanum lycopersicum* | 21 | Vacuole/PM/Chloroplast/Ribosome | Translation |
| 60S ribosomal protein L18a-like | UPI0002B4A10B | *Cucumis sativus* | 21 | PM/Ribosome | Translation |
| 60S ribosomal protein L23a | I1N527 | *Glycine max* | 18 | Ribosome | Translation |
| 60S ribosomal protein L24-like | I1GLQ1 | *Brachypodium distachyon* | 18 | Chloroplast/PM/Cyt | Signaling |
| 60S ribosomal protein L26-1 | A0A498JM53 | *Malus domestica* | 17 | TP/Golgi/Chloroplast/PM | Translation |
| 60S ribosomal protein L30-like | I1L7D4 | *Glycine max* | 10 | Ribosome | Translation |
| 60S ribosomal protein L37-3-like | A0A0A0L819 | *Cucumis sativus* | 11 | Ribosome | Translation |
| 60S ribosomal protein L37a | K7VP92 | *Solanum tuberosum* | 10 | Ribosome | Translation |
| 60S ribosomal protein L5-1 | Q0JGY1 | *Oryza sativa* | 35 | PM/Vacuole/Chloroplast/nucleus | Translation |
| 60S ribosomal protein L6 | P34091 | *Mesembryanthemum crystallinum* | 26 | PM/ER/Chloroplast/Nucelo | Translation |
| 60S ribosomal protein L7-3-like | M1BV98 | *Solanum tuberosum* | 28 | Ribosome | Translation |
| 60S ribosomal protein L8 | P29766 | *Solanum lycopersicum* | 28 | Ribosome | Translation |
| 60S ribosomal protein L9, partial | F4YFF3 | *Camellia sinensis* | 19 | Ribosome | Translation |
| 6-phosphofructokinase 6-like | UPI00046DFF87 | *Prunus mume* | 59 | Cytosolic complex | Glycolysis/Phosphorylation |
| AAA-type ATPase family protein | F4JCC9 | *Arabidopsis thaliana* | 111 | PM | ATP catabolic process |
| ABC transporter B family member 15 | UPI00046DFF87 | *Prunus mume* | 136 | Integral to membrane | Transport/ATP catabolic process |
| ABC transporter B family member 15-like | UPI0002767F3D | *Solanum lycopersicum* | 137 | Integral to membrane/PM | Transport/ATP catabolic process |
| ABC transporter C family member 2 | Q42093 | *Arabidopsis thaliana* | 182 | Membrane/TP/ | Transport/ATP catabolic process |
| ABC transporter C family member 4 | A0A3Q7I6V8 | *Solanum lycopersicum* | 170 | TP/Integral to ER/PM/nucleus | Response to water deprivation/Transport |
| ABC transporter F family member 4 | A0A2I4FMJ1 | *Juglans regia* | 81 | | ATP catabolic process |
| Acetyl-CoA carboxylase beta subunit | L0E7F9 | *Camellia sinensis* | 58 | Chloroplast/Plastid | Lipid metabolic process |
| Actin-1 | P30164 | *Pisum sativum* | 42 | Cyt/Plasmodesmata/PM | Response to stress/Organización del Cytoskeleton/Regulation of protein localization |
| Actin-1 | Q10DV7 | *Oryza sativa* | 42 | Chloroplast/Cyt/Plasmodesmata/PM | Cytoskeleton/protein catabolic process |
| Actin-1 | B4FPG2 | *Zea mays* | 52 | Cyt/Cytoskeleton | |
| Acyl-CoA dehydrogenase family member 10-like | K3ZQQ2 | *Setaria italica* | 93 | | Root hair elongation |
| Beta adaptin family protein isoform 1 | A0A061GCM9 | *Theobroma cacao* | 100 | PM/Clathrin adaptor complex | Vesicle mediated transport |
| Adenosine kinase 2-like isoform X2 | A0A1D6E5Z6 | *Zea mays* | 29 | Cyt/Chloroplast/PM | Phosphorylation |
| Adenylate kinase 4 | Q08480 | *Oryza sativa* | 27 | PM/Mitochondria | Phosphorylation |
| ADP,ATP carrier protein 1, mitochondrial-like isofo | A0A1S2YWC7 | *Cicer arietinum* | 40 | Mitochondria/Integral to membrane | Transport |
| ADP,ATP carrier protein 3, mitochondrial-like | A0A1S2YJ61 | *Cicer arietinum* | 40 | Mitochondria/Integral to membrane | Transport |
| ADP-ribosylation factor 1 | P36397 | *Arabidopsis thaliana* | 21 | Endosome/Golgi | Protein transport/Vesicle mediated transport |
| ADP-ribosylation factor 3 | P40940 | *Arabidopsis thaliana* | 20 | Golgi/PM | Protein transport/Vesicle mediated transport |
| ADP-ribosylation factor-like 8 | H1AD89 | *Nicotiana tabacum* | 19 | Intracellular | Signaling GTPase |
| Aldolase-type TIM barrel family protein | A0A178UD24 | *Arabidopsis thaliana* | 48 | Chloroplast/Plastid/Mitochondria/Cyt | Glycolysis |
| Alpha,alpha-trehalose-phosphate synthase [UDP-forming] | UPI00046DC9CC | *Prunus mume* | 97 | | Metabolic process |
| Alpha-tubulin 1 | M1TCB6 | *Salix arbutifolia* | 50 | Microtubules/Cytoskeleton | Protein polymerization/ GTP catabolic process |
| Alpha-tubulin 3 | S8CDY1 | *Genlisea aurea* | 45 | Microtubules/Cytoskeleton | Protein polymerization/ GTP catabolic process |
| AP-1 complex subunit mu-1-I-like | UPI00046DBC20 | *Prunus mume* | 49 | Clathrin adaptor complex | Protein transport/Vesicle mediated transport |
| AP-2 complex subunit alpha-2-like | UPI0002C30E13 | *Fragaria vesca* | 113 | Clathrin adaptor complex | Protein transport/Vesicle mediated transport |
| AP-2 complex subunit mu-like | UPI000276C248 | *Solanum lycopersicum* | 49 | Clathrin adaptor complex | Protein transport/Vesicle mediated transport |
| Aquaporin 7 | G3E8E1 | *Triticum aestivum* | 30 | Integral component of membrane | Transport |
| Aquaporin PIP1;1 | P61837 | *Arabidopsis thaliana* | 31 | Vacuole/PM/Integral to membrane/M | Water transport/ Response to salt and osmotic stress |
| Aquaporin PIP1;2 | A0A6A3BQM0 | *Hibiscus syriacus* | 33 | Integral to membrane | Transport |
| Aquaporin PIP1;4 | UPI00046DDEA9 | *Prunus mume* | 31 | Integral to membrane | Transport |

| | | | | | |
|---|---|---|---|---|---|
| Aquaporin PIP1;4 | A0A222C4Z4 | *Fragaria vesca* | 31 | Integral to membrane | Transport |
| Aquaporin PIP1;5 | M8BEA3 | *Aegilops tauschii* | 24 | Membrane/Integral component of m | Transport |
| Aquaporin PIP2;1 | M1F3S6 | *Quercus petraea* | 30 | Membrane/Integral component of m | Transport |
| Aquaporin PIP2;5 | M7YU72 | *Triticum urartu* | 30 | Membrane/Integral component of m | Transport |
| Ascorbate peroxidase2 | B4G031 | *Zea mays* | 27 | PM/Chloroplast/Cell wall | Response to oxidative stress |
| Aspartic proteinase-like | UPI0002BC88E7 | *Solanum lycopersicum* | 56 | Vacuole | Proteolysis/Lipid metabolic process |
| ATP binding cassette subfamily B4 isoform 1 | A0A061DZI3 | *Theobroma cacao* | 139 | Integral to membrane | Transport/ATP catabolic process |
| ATP synthase subunit 9 | P13547 | *Triticum aestivum* | 8 | Mitochondria | ATP synthesis coupled to electron transport |
| ATP synthase subunit alpha, mitochondrial | P05492 | *Oenothera biennis* | 56 | Mitochondria | ATP synthesis coupled to electron transport |
| ATP synthase subunit beta, mitochondrial | P17614 | *Nicotiana plumbaginifolia* | 60 | Mitochondria | ATP synthesis coupled to electron transport |
| ATP synthase subunit beta, mitochondrial | M7Y940 | *Triticum urartu* | 58 | Mitochondria | ATP synthesis |
| ATP synthase subunit beta, mitochondrial-like | UPI0002C2F91D | *Fragaria vesca* | 59 | Mitochondria | ATP synthesis coupled proton transport |
| ATP synthase subunit d, mitochondrial-like | A0A0A0KRL4 | *Cucumis sativus* | 20 | TP/Ribosome/Chloroplast/Mitochon | ATP synthesis coupled proton transport |
| ATP synthase subunit delta | UPI00046DD172 | *Prunus mume* | 21 | Mitochondria | |
| ATP synthase subunit gamma, mitochondrial | P26360 | *Ipomoea batatas* | 36 | Mitochondria/Cell wall/Chloroplast | Protein catabolic process |
| ATPase family AAA domain-containing protein 3-B-like | UPI0002767FD0 | *Solanum lycopersicum* | 70 | Cell wall | |
| ATPase, AAA-type, CDC48 protein isoform 1 | A0A061ETA0 | *Theobroma cacao* | 90 | | |
| ATP-dependent Clp protease ATP-binding subunit clpA homolog CD4B | A0A6N2C782 | *Solanum lycopersicum* | 102 | Chloroplast | Metabolic process |
| ATP-dependent zinc metalloprotease FTSH 10 | A0A0A0KH94 | *Cucumis sativus* | 91 | Integral to membrane | Protein catabolic process |
| Beta-tubulin, partial | J9RVH4 | *Populus trichocarpa* | 16 | Microtubules/Cytoskeleton | Protein polymerization/ GTP catabolic process |
| Bifunctional polymyxin resistance protein ArnA-like | A0A6N2BQ61 | *Solanum chilense* | 43 | Cyt | |
| Brefeldin A-inhibited guanine nucleotide-exchange protein 2-like | A0A1S2XPV0 | *Cicer arietinum* | 199 | Nucleus/Cyt | Regulation of ARF protein signal transduction |
| BR-signaling kinase 1 | A0A061FL71 | *Theobroma cacao* | 58 | Plasmodesmata/PM | Phosphorylation/Myristoylation |
| C2 domain-containing family protein | UPI0001D464F1 | *Populus trichocarpa* | 220 | Plasmodesmata/Cyt/PM | |
| Ca+2-binding EF hand protein | O23959 | *Glycine max* | 27 | | |
| Calcium-binding EF-hand family protein isoform 1 | A0A061F1V7 | *Theobroma cacao* | 62 | Nucleus/Cyt | |
| Calcium-dependent protein kinase 26 | F4JTL3 | *Arabidopsis thaliana* | 53 | Nucleus | Phosphorylation |
| Calcium-dependent protein kinase 9-like | A0A1S2XM13 | *Cicer arietinum* | 63 | Nucleus/Cyt | Phosphorylation |
| Calcium-transporting ATPase 2 | 460375325 | *Solanum lycopersicum* | 111 | Integral to membrane/ER/PM | Transport/Membrane fusion/ER to Golgi vesicle mediated transport |
| Calcium-transporting ATPase 9 | M8AMH4 | *Triticum urartu* | 169 | Membrane/Integral component of m | Transport |
| Calmodulin | P04464 | *Triticum aestivum* | 17 | Cyt/Vacuole/PM/Nucleus | Response to stimulus/Signaling /Metabolic process |
| Calnexin homolog | K3Y6F3 | *Setaria italica* | 60 | PM/Cytoplasmic membrane bounde | Protein folding |
| Calnexin-like protein precursor | Q4W5U7 | *Solanum lycopersicum* | 61 | ER | Protein folding |
| Calnexin-like protein, partial | S8CMU9 | *Genlisea aurea* | 53 | ER | Protein folding |
| Calreticulin-1 precursor | A0A762 | *Glycine max* | 48 | ER | Protein folding |
| Calreticulin-like | UPI0002C37537 | *Fragaria vesca* | 48 | ER lumen | Protein folding |
| Carbohydrate-binding-like fold-containing protein | Q9LZQ4 | *Arabidopsis thaliana* | 133 | ER/Vacuole/Golgi/Cell wall | Myristoylation |
| Catalase isozyme 1 | Q01297 | *Ricinus communis* | 56 | Cell wall/Mitochondria/Chloroplast/P | Response to stress |
| Cation/H(+) antiporter 18-like | A0A0A0KUY7 | *Cucumis sativus* | 87 | | |
| Cellulose synthase A catalytic subunit 2 [UDP-forming] | O48947 | *Arabidopsis thaliana* | 122 | PM/Integral component of membrane | |
| Chaperonin CPN60-1, mitochondrial precursor | P29185 | *Zea mays* | 61 | Cyt/Mitochondria | Protein folding/Response to stress |
| Chemocyanin | M8B4A2 | *Aegilops tauschii* | 17 | Cyt | |

| | | | | | |
|---|---|---|---|---|---|
| Cinnamate 4-hydroxylase | G3D5B5 | *Leucaena leucocephala* | 58 | PM/Cell wall/ER | Oxide-reduction |
| Coatomer subunit delta | A0A061DXT8 | *Theobroma cacao* | 58 | Clathrin adaptor complex/Cyt/Plasmodesmata | Protein transport/Vesicle mediated transport |
| **Clathrin heavy chain 1** | Q2RBN7 | *Oryza sativa* | 193 | Clathrin coat of coated pit/Clathrin c | Vesicle mediated transort/Protein transport |
| **Clathrin heavy chain 1** | Q0WNJ6 | *Arabidopsis thaliana* | 193 | Clathrin coat of coated pit/Clathrin c | Protein transport/Endocitosis |
| **Clathrin heavy chain 1-like** | A0A3Q7HNN1 | *Solanum lycopersicum* | 193 | Cyt/Clathrin coat vesicle | Protein transport/ Vesicle mediated transport |
| **Clathrin heavy chain 1-like** | UPI0002C3639C | *Fragaria vesca* | 193 | Cyt/Clathrin coat vesicle | Protein transport/ Vesicle mediated transport |
| **Clathrin, heavy chain isoform 1** | A0A061FT37 | *Theobroma cacao* | 193 | Clathrin coat of trans-Golgi network vesicle | Protein transport/Vesicle mediated transport |
| Coatomer subunit alpha-1-like | UPI00046DF541 | *Prunus mume* | 137 | PM/COP1 | Protein transport/Vesicle mediated transport |
| Coatomer subunit alpha-1-like | A0A0A0K130 | *Cucumis sativus* | 136 | PM/COP1 vesicle | Protein transport/Vesicle mediated transport |
| Coatomer subunit alpha-1-like | UPI0002C3407C | *Fragaria vesca* | 137 | PM/COP1 vesicle | Protein transport/Vesicle mediated transport |
| Coatomer subunit beta-1-like | A0A0A0LCU8 | *Cucumis sativus* | 106 | PM/COP1 vesicle | Protein transport/Vesicle mediated transport |
| Coatomer subunit beta-2-like | UPI000276943D | *Solanum lycopersicum* | 105 | Membrane coat | Protein transport/Vesicle mediated transport |
| Coatomer subunit gamma-2-like | UPI00027689D6 | *Solanum lycopersicum* | 98 | COPI vesicle/Chloroplast | Protein transport/ Vesicle mediated transport |
| Coatomer, beta subunit isoform 1 | A0A061GJB6 | *Theobroma cacao* | 106 | COP1 | Protein transport/Vesicle mediated transport |
| Cullin-associated and neddylation dissociated | A0A061G8M4 | *Theobroma cacao* | 135 | Cyt/Cell wall | Lipid storage/Cell wall organization/Sugar signalig |
| D-3-phosphoglycerate dehydrogenase | B6SL40 | *Zea mays* | 64 | Chloroplast/Mitochondria | Oxide-reduction |
| D-3-phosphoglycerate dehydrogenase, chloroplastic-like | UPI0002C32A7C | *Fragaria vesca* | 63 | Chloroplast/Mitochondria/Membrane | Oxide-reduction |
| DEAD/DEAH box RNA helicase family protein isoform 1 | A0A061GK01 | *Theobroma cacao* | 55 | | |
| DEAD-box ATP-dependent RNA helicase 52C-like | A0A1S2YC83 | *Cicer arietinum* | 67 | | |
| DEAD-box ATP-dependent RNA helicase 8-like | UPI0002B4BA64 | *Cucumis sativus* | 56 | | |
| Delta(24)-sterol reductase-like | UPI0002C306D2 | *Fragaria vesca* | 67 | Integral to membrane/Vacuole/PM | Oxide-reduction |
| Delta-1-pyrroline-5-carboxylate synthase | O65361 | *Mesembryanthemum crystallinum* | 78 | Membrane/Chloroplast | Response to stress/Hiperosmotic salinity response |
| Developmentally regulated G-protein | Q9SVA6 | *Arabidopsis thaliana* | 41 | Cytosol | GTP catabolic process |
| Diacylglycerol kinase iota-like | A0A3Q7ICL0 | *Solanum lycopersicum* | 55 | | Phosphorylation/Signaling |
| DNA damage-inducible protein 1-like | UPI0002C33D25 | *Fragaria vesca* | 45 | Nucleus/Cyt | Proteolysis/Response to stress |
| DnaJ homolog subfamily C GRV2-like | UPI0002BC90E9 | *Solanum lycopersicum* | 283 | | Response to stress |
| DnaJ protein homolog ANJ1 | P43644 | *Atriplex nummularia* | 47 | Cell wall/PM | Response to salt stress/Regulation of ATP activity |
| Dolichyl-diphosphooligosaccharide-protein glycosyltransferase | A0A061DGI3 | *Theobroma cacao* | 50 | Nucleus/Plasmodesmata/TP/Cell wa | Golgi vesicle transport/Cell wall |
| Dolichyl-diphosphooligosaccharide--protein glycosyltransferase | A0A3Q7J3H2 | *Solanum lycopersicum* | 86 | Chloroplast/ER/PM | Response to stress/Response to salt stress |
| DUF21 domain-containing protein At4g14240-like | UPI0002209ABB | *Cucumis sativus* | 56 | Membrane/Mitochondria | |
| DUF3411 domain protein | A0A072TIU5 | *Medicago truncatula* | 42 | | |
| Dynamin-2B | M5VHG2 | *Prunus mume* | 102 | Cytosol | GTP catabolic process |
| E3 ubiquitin-protein ligase UPL2-like | A0A314YPJ6 | *Prunus yedoensis* | 403 | | Protein ubiquitination |
| Elongation factor 1 gamma-like protein, partial | S8CML0 | *Genlisea aurea* | 48 | Vacuole/PM/Cell wall | Translation |
| Elongation factor 1-alpha | UPI00046DD67D | *Prunus mume* | 49 | Cyt | GTP catabolic process/Translation |
| Elongation factor 1-delta 1 | B4FNT1 | *Zea mays* | 25 | PM | Translation |
| Elongation factor 2 | O23755 | *Beta vulgaris* | 94 | Cyt | GTP catabolic process/Translation |
| Elongation factor Tu | Q9ZT91 | *Arabidopsis thaliana* | 49 | Cell wall/Mitochondria/Cyt | GTP catabolic process |
| Endoplasmic reticulum [ER]-type calcium ATPase isoform 1 | A0A061GXS4 | *Theobroma cacao* | 117 | TP/Integral to ER/PM/Nucleus | Transport of Ca and Mn |

| | | | | | |
|---|---|---|---|---|---|
| Endoplasmin homolog | K3XVC0 | *Setaria italica* | 93 | TP/ER lumen/PM/Cytoplasmic membrane bounded vesicle | Transport/Water transport/Response to stress/Response to salt stress |
| Endoplasmin homologo | UPI00046E14C3 | *Prunus mume* | 93 | TP/ER lumen/PM/Plasmodesmata | Water transport/Myristoylation/Glycolysis/Stres in ER/Response to salt stress |
| Enolase | Q43130 | *Mesembryanthemum crystallinum* | 48 | Cyt | Metabolic process/Glycolysis |
| Eukaryotic aspartyl protease family protein isoform | A0A061E0D5 | *Theobroma cacao* | 61 | Cell wall/Membrane | Proteolysis |
| Eukaryotic initiation factor 4A-2-like | A0A3Q7HMP2 | *Solanum lycopersicum* | 47 | Cyt/Cell wall/PM | Translation |
| Eukaryotic peptide chain release factor GTP-binding subunit ERF3A-like | K3Y6E4 | *Setaria italica* | 59 | Vacuole | GTP catabolic process |
| Eukaryotic peptide chain release factor subunit 1-1 | B6U1I9 | *Zea mays* | 49 | Cyt | Translation |
| Eukaryotic translation initiation factor 2 family protein isoform 1 | A0A061DJB3 | *Theobroma cacao* | 158 | Cyt | Translation |
| Eukaryotic translation initiation factor 2 subunit alpha-like | M5X0K8 | *Prunus mume* | 39 | Nucleus | Translation |
| Eukaryotic translation initiation factor 3 subunit A-like | UPI0002C2DDD4 | *Fragaria vesca* | 111 | Cyt/PM | Translation |
| Eukaryotic translation initiation factor 3 subunit B-like | UPI00046E04E2 | *Prunus mume* | 83 | Eukaryotic translation initiation factor 3 | Translation |
| Eukaryotic translation initiation factor 3 subunit C | O49160 | *Arabidopsis thaliana* | 103 | Nucleus/Cyt/PM/Cell wall | Translation |
| Eukaryotic translation initiation factor 3 subunit D-like | A0A1S2XCE8 | *Cicer arietinum* | 64 | Eukaryotic translation initiation factor 3 | Translation regulation |
| Eukaryotic translation initiation factor 3 subunit E-like | A0A0A0L0F9 | *Cucumis sativus* | 51 | Cyt | Response to stress/Response to salt stress |
| Eukaryotic translation initiation factor 3 subunit H-like | M1BYD6 | *Solanum lycopersicum* | 39 | Cyt | Translation |
| Eukaryotic translation initiation factor 4A1 | A0A061GKK2 | *Theobroma cacao* | 47 | Nucleus/Cyt/PM | Translation |
| Eukaryotic translation initiation factor 5A-2 | P24922 | *Nicotiana plumbaginifolia* | 17 | Nucleus/Cyt | Translation |
| Exportin 1A | Q9SMV6 | *Arabidopsis thaliana* | 123 | Nucleus/Cyt | Protein export from nucleus/Protein transport |
| F1F0- ATPase inhibitor protein | B6TGD4 | *Brachypodium distachyon* | 12 | Mitochondria/Nucleus | Protein catabolic process/Proteosome assembly |
| Fatty acyl-CoA synthetase A | M8BUY6 | *Aegilops tauschii* | 74 | PM | Metabolic process |
| Flavoprotein WrbA-like | A0A1S2XDH1 | *Cicer arietinum* | 22 | Plasmodesmata/PM | Oxide-reduction |
| Fructokinase-1 | Q9SID0 | *Arabidopsis thaliana* | 35 | Golgi/Plasmodesmata/PM/Cyt | Response to stimulus/Phosphorylation |
| Fructose-bisphosphate aldolase cytoplasmic isozyme | M7ZGS6 | *Triticum urartu* | 39 | Cyt/PM/Vacuole/Plasmodesmata/Chloroplast | Metabolic process/Glycolysis |
| Fructose-bisphosphate aldolase cytoplasmic isozyme | UPI00046DD6EA | *Prunus mume* | 38 | Cyt/PM/Vacuole/Plasmodesmata/Chloroplast | Glycolysis |
| Fructose-bisphosphate aldolase, cytoplasmic isozyme | P29356 | *Spinacia oleracea* | 38 | Cyt | Glycolysis |
| Gamma carbonic anhydrase 1, mitochondrial-like | A0A0A0KWN9 | *Cucumis sativus* | 30 | Mitochondria | Metabolic process/ Starch biosynthesis |
| Gamma carbonic anhydrase-like 2, mitochondrial-like | A0A0A0LR76 | *Cucumis sativus* | 27 | Mitochondria | |
| Gamma carbonic anhydrase-like 2, mitochondrial-like | UPI0002768717 | *Solanum lycopersicum* | 28 | Mitochondria | Response to stress/Water transport/Golgi organization/Protein catalitic process |
| Glucose-6-phosphate 1-dehydrogenase, cytoplasmic isoform 1 | UPI00027688C2 | *Solanum lycopersicum* | 59 | Cyt | Metabolic process |
| Glutamate decarboxylase 1 | UPI00046E1A6C | *Prunus mume* | 56 | | Metabolic process |
| Glutamine synthetase | UPI0002769AB4 | *Solanum lycopersicum* | 39 | Cyt | Metabolic process |

| | | | | | |
|---|---|---|---|---|---|
| Glyceraldehyde-3-phosphate dehydrogenase | M4SIP8 | *Atropa belladonna* | 34 | Cyt | Metabolic process/ Oxide-reduction/Glycolysis |
| Glyceraldehyde-3-phosphate dehydrogenase, cytosolic | P17878 | *Mesembryanthemum crystallinum* | 37 | Chloroplast/Cell wall/PM/Plasmodesmata | Response to salt stress/Response to stress/Protein catabolic process |
| Glyceraldehyde-3-phosphate dehydrogenase, cytosolic | P26518 | *Magnolia liliflora* | 37 | Chloroplast/Cell wall/PM/Plasmodesmata | Response to salt stress/Response to stress/Protein catabolic process |
| Glyceraldehyde-3-phosphate dehydrogenase-like isoform 1 | A0A0A0K8C1 | *Cucumis sativus* | 37 | | Oxide-reduction/Glycolysis |
| Glycerophosphoryl diester phosphodiesterase 1 | M7ZR94 | *Triticum urartu* | 81 | Cyt membrane bounded vesicle/And | Lipid metabolic process/Metabolic process |
| GTP-binding nuclear protein Ran-3 | K3XLR2 | *Setaria italica* | 25 | Golgi/PM/Plasmodesmata/Cell wall | Protein transport/Response to salt stress/GTP catalitic process |
| GTP-binding protein SAR1A-like | A0A3Q7HK84 | *Solanum lycopersicum* | 22 | ER/Golgi | Protein transport/Vesicle mediated transport |
| Guanine nucleotide-binding protein subunit beta-2 | P93398 | *Nicotiana tabacum* | 41 | ER | Response to stress/Protein targeting to membrane/Golgi vesicle transport |
| Guanine nucleotide-binding protein subunit beta-like | O24076 | *Medicago sativa* | 36 | | |
| Heat shock 70 kDa protein 15-like | A0A3Q7J9R5 | *Solanum lycopersicum* | 43 | PM/Plasmodesmata/Cell wall/Cyt | Oxide-reduction/ Response to stress |
| Heat shock cognate protein 70-1 | A0A061H032 | *Theobroma cacao* | 71 | | Response to stress |
| Heat shock cognate protein 80-like | A0A1S2Y850 | *Cicer arietinum* | 80 | Cyt/Cell wall/Mitochondria | Response to salt stress |
| Heat shock cognate protein 90-2 | UPI0002C303FE | *Fragaria vesca* | 80 | Cyt/Cell wall/Mitochondria | Response to salt stress |
| Heat shock protein 81-3 | F4Y5A9 | *Triticum urartu* | 80 | Cyt/Cell wall/Mitochondria | Response to salt stress and water depravation |
| Heat shock protein 89-1 | F4JFN3 | *Arabidopsis thaliana* | 91 | Mitochondria/Cell wall | Response to stress |
| Heat shock protein 90-2 | K9JHZ2 | *Nicotiana attenuata* | 80 | Cyt/TP | Response to water deprivation/Response to salt stress |
| Heat shock protein 90-4 | O03986 | *Arabidopsis thaliana* | 80 | Golgi/Nucleus/Cell wall/Chloroplast | Response to stress |
| Histone H2A.6 | B6SHX9 | *Zea mays* | 16 | Nucleus | Nucleosome assembly |
| Histone H2B.6 | A2WKT1 | *Oryza sativa* | 16 | Nucleus | Nucleosome assembly |
| Histone H4 | P59259 | *Arabidopsis thaliana* | 11 | Nucleus/TP/Golgi/Plasmodesmata | Response to water deprivation |
| Hypersensitive-induced response protein 1 | Q9FM19 | *Arabidopsis thaliana* | 31 | Vacuole/Golgi/PM/Cyt/Plasmodes mata | Signaling |
| Hypersensitive-induced response protein 2-like | M0ZPM8 | *Solanum tuberosum* | 31 | Vacuole/Plasmodesmata/Clorolasto/PM | |
| Importin subunit alpha | O22478 | *Solanum lycopersicum* | 59 | Nucleus/Cyt | Protein transport |
| Importin subunit alpha-1a-like | A0A3Q7EFC3 | *Solanum lycopersicum* | 59 | Cyt | Protein transport |
| Importin subunit beta-1 | M7YB37 | *Triticum urartu* | 96 | Nucleus | Protein transport |
| Importin subunit alpha | A0A3Q7EFC3 | *Solanum lycopersicum* | 100 | Chloroplast/Nucleus | Protein transport |
| Importin-7 homolog | UPI000276C169 | *Solanum lycopersicum* | 119 | Nucleus | Protein transport |
| Inactive leucine-rich repeat receptor-like protein kir | A0A0A0KM77 | *Cucumis sativus* | 91 | Membrane | Phosphorylation |
| Inactive serine/threonine-protein kinase scy1-like isoform X1 | A0A1S2XWA4 | *Cicer arietinum* | 86 | | Phosphorylation |
| Guanylate binding protein 2 | A0A1S3B0B9 | *Cucumis melo* | 120 | Nucleus/Chloroplast | GTP catabolic process |
| Intron-binding protein aquarius-like | K4A4S4 | *Setaria italica* | 179 | PM/Chloroplast | mRNA splicing |
| Isocitrate dehydrogenase [NAD] regulatory subunit 3 | Q945K7 | *Arabidopsis thaliana* | 41 | Mitochondria/Chloroplast | Metabolic Process/ Oxide-reduction |
| Isocitrate dehydrogenase [NADP], chloroplastic | Q40345 | *Medicago sativa* | 48 | Cyt/Plasmodesmata/Chloroplast | Water transport/Regulation of transport/Response to stress/Golgi organization |
| Ketol-acid reductoisomerase | A0A061DS88 | *Theobroma cacao* | 64 | Cyt/Mitochondria/Chloroplast | Response to salt stress/Glycolysis |
| Kinase protein with tetratricopeptide repeat domain isoform 1 | A0A061G6D0 | *Theobroma cacao* | 55 | PM | Phosphorylation |
| Leucine-rich repeat protein kinase family protein | A0A061GN08 | *Theobroma cacao* | 71 | Integral to membrane/PM | Sterol biosynthesis/Signaling |
| Leucine-rich repeat receptor-like serine/threonine-protein kinase BAM1-like | A0A3Q7XWW5 | *Cicer arietinum* | 110 | Integral to membrane | Phosphorylation/Signaling |

| | | | | | |
|---|---|---|---|---|---|
| Leu-rich receptor Serine/threonine protein kinase BAK1 | Q94F62 | *Arabidopsis thaliana* | 68 | PM/Endosome/Integral to membrane | Phosphorylation/Response to stress/Signaling |
| L-gulonolactone oxidase-like | K3YGV3 | *Setaria italica* | 45 | Membrane/Cell wall | Oxide-reduction |
| LMBR1-like membrane protein | Q9M028 | *Arabidopsis thaliana* | 57 | Integral to membrane/PM/Plasmode | Vacuole organization |
| Long chain acyl-CoA synthetase 9 isoform 1 | S8CZD8 | *Theobroma cacao* | 76 | Chloroplast/Membrane | Metabolic process/Faty acid biosynthesis |
| LRR receptor-like serine/threonine-protein kinase | N1QWM1 | *Aegilops tauschii* | 99 | Cyt membrane bounded vesicle | Phosphorylation |
| LRR receptor-like serine/threonine-protein kinase At1g56130-like | UPI000276B496 | *Solanum lycopersicum* | 113 | PM | Phosphorylation |
| L-type lectin-domain containing receptor kinase S.! | K4B7G2 | *Solanum lycopersicum* | 70 | | Phosphorylation |
| Luminal-binding protein | Q42434 | *Spinacia oleracea* | 74 | ER | Response to stress |
| Luminal-binding protein 4 | Q03684 | *Nicotiana tabacum* | 74 | Cell wall/ER lumen/PM/Vacuole | Oxide-reduction/ER associated protein catabolic process |
| Malate dehydrogenase, cytoplasmic | O24047 | *Mesembryanthemum crystallinum* | 35 | Vacuole/PM/Nucleo/Chloroplast | Metabolic process/Response to salt stress |
| Mediator of RNA polymerase II transcription subun | Q94AH9 | *Arabidopsis thaliana* | 34 | Cyt/Nucleus | Metabolic process/Regulation of transcription |
| Membrane steroid-binding protein 2-like | UPI0002768EA1 | *Solanum lycopersicum* | 25 | Chloroplast/ER/PM | Oxide-reductio/Metabolic process |
| Metallopeptidase M24 family protein isoform 1 | A0A061ER94 | *Theobroma cacao* | 43 | PM/Nucleus | Protein transport/Proteolysis |
| Methyltransferase PMT2-like isoform X1 | A0A1S2XIY9 | *Cicer arietinum* | 69 | | Methylation |
| Mitochondrial 2-oxoglutarate/malate carrier protein-like | A0A0A0KZ03 | *Cucumis sativus* | 32 | Integral to membrane/PM/Mitochond | Transport |
| Mitochondrial malate dehydrogenase | I6YI03 | *Linum usitatissimum* | 31 | Cell wall/Membrane | Metabolic process/Oxide-reduction/Response to salt stress |
| Mitochondrial outer membrane protein porin of 36 kDa-like | A0A3Q7EA80 | *Solanum lycopersicum* | 29 | Cell wall/Chloroplast/PM/Vacuole | Transport |
| Mitochondrial pyruvate carrier 2-like | A0A0D3DFE2 | *Brassica rapa* | 12 | Mitochondria | Glycolysis |
| Mitochondrial Rho GTPase | A0A0A0LIM9 | *Cucumis sativus* | 72 | Mitochondria | GTP catabolic process/Signaling via GTPases |
| Mitochondrial-processing peptidase subunit alpha | R7WA10 | *Aegilops tauschii* | 75 | Mitochondria/Chloroplast/PM | Metabolic process/Response to salt stress |
| Mitochondrial-processing peptidase subunit alpha-like | A0A3Q7J3G4 | *Solanum lycopersicum* | 55 | Mitochondria | Proteolysis/Metabolic process |
| Mitochondrial-processing peptidase subunit beta-like | A0A0A0L0I1 | *Cucumis sativus* | 59 | TP/Mitochondria/Chloroplast/Cell wa | Response to stress/Response to salt stress/Protein catabolic process |
| Monocopper oxidase-like protein SKU5-like | A0A1S2Z641 | *Cicer arietinum* | 66 | Plasmodesmata/PM/TP | Root hair elongation/Oxidación-reduccion |
| Monodehydroascorbate reductase | Q42711 | *Cucumis sativus* | 47 | PM/Chloroplast | Oxide-reduction/Response to stress |
| Myosin-related isoform 1 | A0A061GTA5 | *Theobroma cacao* | 110 | | |
| NAD(P)H dehydrogenase B1, mitochondrial-like | UPI000276C5F7 | *Solanum lycopersicum* | 65 | Mitochondria inner membrane | Oxide-reductio |
| NADH dehydrogenase (ubiquinone) Fe-S protein 7 | Q42577 | *Arabidopsis thaliana* | 24 | Mitochondria | Glycosilation |
| NADH dehydrogenase [ubiquinone] 1 alpha subcomplex subunit 5 | UPI0002B487B6 | *Cucumis sativus* | 19 | Mitochondria/Chloroplast | Protein catabolic process/Response to salt stress |
| NADH dehydrogenase [ubiquinone] 1 alpha subcomplex subunit 9 | A0A1S2XV81 | *Cicer arietinum* | 44 | Mitochondria | Protein catabolic process/Response to salt stress |
| NADH dehydrogenase [ubiquinone] iron-sulfur protein 1 | UPI0002C2E6B3 | *Fragaria vesca* | 80 | Mitochondria/Chloroplast | Protein catalitic process/Response to stress |
| NADH dehydrogenase [ubiquinone] iron-sulfur protein 1 | Q43644 | *Solanum tuberosum* | 80 | Chloroplast/Mitochondria | Response to stress |
| NADH dehydrogenase subunit 5 | G9JLP7 | *Millettia pinnata* | 75 | Mitochondria/Integral component of | Oxide-reduction |
| NADH dehydrogenase subunit 7 | G9JLR7 | *Millettia pinnata* | 45 | Mitochondria | Oxide-reduction |
| NADH dehydrogenase subunit 9 | R4I225 | *Raphanus sativus* | 23 | Mitochondria | Oxide-reduction |
| NADH-ubiquinone oxidoreductase chain 1 | Q01148 | *Triticum aestivum* | 36 | Cyt/Integral to membrane | Metabolic process |

| | | | | | |
|---|---|---|---|---|---|
| Nascent polypeptide-associated complex subunit alpha-like protein 1 | A0A5E4FKR1 | *Prunus mume* | 22 | Ribosome | Response to salt stress |
| N-ethylmaleimide sensitive fusion protein | T1RU75 | *Silene vulgaris* | 94 | Vacuola/Golgi/Plasmodesmata/PM | |
| Neutral alpha-glucosidase AB-like | A0A1S2YCV8 | *Cicer arietinum* | 104 | ER/Chloroplast | Golgi vesicle transport/Cell wall |
| Nicalin-like | UPI0002B48A57 | *Cucumis sativus* | 62 | Golgi/Plasmodesmata/PM/TP/ER/ Mitochondria | Myroistoylation |
| Niemann-Pick C1 protein-like | A0A1S2XML4 | *Cicer arietinum* | 140 | | |
| Non-specific phospholipase C4-like | J3N9D2 | *Oryza brachyantha* | 57 | Vacuole/Cyt/PM | Response to stress/Phospholipid catabolism |
| Nuclease domain-containing protein 1 | M7YQT8 | *Triticum urartu* | 114 | Nuclear Envelope/ER/PM/Cyt/Cell w | Metabolic process/Response to stress |
| Nucleolar protein 5-2 | UPI00046E2350 | *Prunus mume* | 60 | | |
| Nucleoside diphosphate kinase 1 | O81372 | *Mesembryanthemum crystallinum* | 16 | | Biosynthesis of GTP |
| Nucleosome assembly protein 1-like 1-like isoform 1 | A0A3Q7J2T2 | *Solanum lycopersicum* | 43 | Nucleus | Nucleosome assembly |
| Obg-like ATPase 1-like | A0A0R4J410 | *Glycine max* | 44 | Cyt/Plasmodesmata | Response to salt stress/Glycolysis/Protein transport |
| PAM domain (PCI/PINT associated module) protei | A0A061FEG6 | *Theobroma cacao* | 56 | Proteasome/Nucleus | Protein catabolic process |
| Patellin-3-like | UPI0002C2E42C | *Fragaria vesca* | 67 | Integral to membrane | Transport |
| Peptide chain release factor eRF subunit 1 | A0A178VIS2 | *Arabidopsis thaliana* | 49 | Cyt/PM | Translation |
| Peroxidase 1 | M7ZBJ0 | *Triticum urartu* | 37 | Membrane/ Cyt protein bounded | Response to estress |
| Peroxidase 72-like | A0A1S2YYU4 | *Cicer arietinum* | 37 | | Oxide-reduction/Response to stress |
| Phosphate carrier protein | R7VZF2 | *Aegilops tauschii* | 32 | Integral to membrane/Mitochondria/PM | Transport |
| Phospho-2-dehydro-3-deoxyheptonate aldolase 1 | UPI00046D98F7 | *Prunus mume* | 68 | Membrane/Chloroplast | Metabolic process/Response to stress |
| Phosphoenolpyruvate carboxykinase 1 | R0F3C0 | *Theobroma cacao* | 73 | Cyt/Membrane | Phosphorylation/Response to stress |
| Phosphoenolpyruvate carboxylase 2 | P16097 | *Mesembryanthemum crystallinum* | 109 | Cyt | Metabolic process |
| Phosphoglucomutase, cytoplasmic | P93262 | *Mesembryanthemum crystallinum* | 63 | Cyt | Metabolic process/Glycolysis |
| Phosphoglycerate kinase isoform 1 | A0A061FI02 | *Theobroma cacao* | 42 | Cyt/Plasmodesmata/TP/PM/Nucleus | Phosphorylation/Response to stress/Glycolysis |
| Phospholipase D delta isoform 1 | A0A061DGE7 | *Theobroma cacao* | 100 | Plasmodesmata/PM/Vacuola | Metabolic process/Lipid metabolic process |
| Phosphorus transporter | K4BDP2 | *Solanum lycopersicum* | 38 | Integral to membrane/TP/Mitochondria | Transport/Response to salt stress |
| Plasma intrinsic protein PIP1.2 | M1EY53 | *Helianthemum almeriense* | 30 | Integral to membrane/TP/Mitochondria/Cell wall | Response to salt stress/Response to water deprivation |
| Plasma membrane ATPase | M8AIK4 | *Triticum aestivum* | 105 | Integral to membrane/Vacuole/PM | Transport/Response to water deprivation |
| Plasma membrane ATPase 4-like | P83970 | *Triticum urartu* | 105 | Integral to membrane | Transport/ATP catabolic process |
| Plasma membrane ATPase 4-like | A0A1S2YPD3 | *Cicer arietinum* | 105 | Integral to membrane | Transport/ATP catabolic process |
| Plasma membrane ATPase 4-like | A0A3Q7H652 | *Solanum lycopersicum* | 105 | Integral to membrane/Vacuole/PM | Transport/Response to water deprivation |
| Plasma membrane H+-ATPase | K7TH91 | *Sesuvium portulacastrum* | 105 | Integral to membrane/Vacuole/PM | Transport/Response to water deprivation |
| Plasma membrane H+-transporting ATPase-like protein | UPI0002215CE2 | *Zea mays* | 106 | Integral to membrane/Vacuole/PM | Transport/Response to water deprivation |
| Pleiotropic drug resistance 12 | A0A061GNG0 | *Theobroma cacao* | 164 | Membrane | ATP catabolic process |
| Pleiotropic drug resistance protein 1 | Q76CU2 | *Nicotiana tabacum* | 162 | Cyt | Transport/ATP catabolic process |
| Polyadenylate-binding protein | A0A1S2YIR6 | *Cicer arietinum* | 64 | | Response to stress/Response to salt stress |
| Potassium transporter 4-like | A0A3Q7JU30 | *Solanum lycopersicum* | 89 | Membrane | Transport/Response to stress |
| Pre-mRNA-processing-splicing factor 8-like | A0A3Q7I1D8 | *Solanum lycopersicum* | 278 | Spliceosomal complex | mRNA splicing |
| Probable calcium-binding protein CML49-like | A0A1S2Z7E5 | *Cicer arietinum* | 31 | | Proteolysis/Golgi localization |
| Probable methyltransferase PMT8 | UPI00046DB4A1 | *Prunus mume* | 70 | Golgi/PM/Nucleus | Methylation |
| Probable phospholipid hydroperoxide glutathione peroxidase | Q9LEF0 | *Mesembryanthemum crystallinum* | 19 | PM/Cyt/Chloroplast | Response to salt stress/Response to stress |

| | | | | | |
|---|---|---|---|---|---|
| Proteasome activating protein 200 isoform 1 | A0A061F4N1 | *Theobroma cacao* | 190 | | |
| Proteasome subunit beta type-6 | A0A0R4J321 | *Glycine max* | 25 | TP/Cyt/Nucleus | Response to stress/Water transport/Golgi organization |
| Protein phosphatase 2C 9 | UPI00046DA708 | *Prunus mume* | 31 | PM | Metabolic process |
| Root hair defective 3-like | UPI00046DD765 | *Fragaria vesca* | 89 | Integral to membrane/ER | GTP catabolic process |
| Protein transport protein SEC23-like | A0A1S3CCL1 | *Cucumis melo* | 86 | COPII vesicle coat | Protein transport/ER to Golgi vesicle mediated transport |
| Protein transport protein Sec24-like At3g07100-like | UPI0002C2DF41 | *Fragaria vesca* | 113 | Cyt/COPII vesicle coat | Protein transport/ER to Golgi vesicle mediated transport |
| Protein transport protein Sec61 subunit alpha-like | A0A3Q7IAQ0 | *Solanum lycopersicum* | 52 | Membrane | Protein transport |
| Pto kinase interactor 1 | S8C641 | *Genlisea aurea* | 38 | | Fosforlación/Response to stress |
| Putrescine aminopropyltransferase | O48660 | *Nicotiana sylvestris* | 35 | PM | Metabolic process |
| Pyrophosphate-energized vacuolar membrane proton pump-like | UPI0002C35E23 | *Fragaria vesca* | 80 | TP/Integral to membrane | Transport |
| Pyruvate kinase, cytosolic isozyme | P22200 | *Solanum tuberosum* | 55 | Cyt | Phosphorylation/Glycolysis |
| Pyruvate kinase, cytosolic isozyme-like | UPI0002B45BAC | *Cucumis sativus* | 57 | Cyt/PM | Metabolic process/Glycolysis/Response to stress |
| Rab GDP dissociation inhibitor alpha-like | O65744 | *Cicer arietinum* | 50 | | Protein transport |
| RAB GTPase homolog B1C | P92963 | *Arabidopsis thaliana* | 23 | Vacuole/Golgi | GTP catabolic process/ER to Golgi mediated transport |
| Rac-like GTP-binding protein ARAC1 | Q38902 | *Arabidopsis thaliana* | 22 | PM/Fragmoplast | Signaling by GTPases |
| Ras-related protein RAB1BV | Q39433 | *Beta vulgaris* | 24 | Vacuole/PM | Protein transport/Signalig by GTPases |
| Ras-related protein RABA1f-like | UPI0002C2EE30 | *Fragaria vesca* | 24 | Cyt/Membrane | GTP catabolic process/Vesicle mediated transort/Protein transport |
| Ras-related protein RABA2a-like | A0A3Q7EQ24 | *Solanum lycopersicum* | 24 | Endosome/Cyt | Vesicle mediated transort/Protein transport |
| Ras-related protein RABG3f-like isoform 1 | A0A0V0HPF5 | *Solanum lycopersicum* | 23 | Golgi/PM | Response to salt stress/Response to ER stress/Vesicle mediated transport |
| Receptor-like protein kinase HERK 1-like | K3Z3U9 | *Setaria italica* | 37 | Mitochondria/Cyt membrane bounde | Phosphorylation |
| Regulatory particle triple-A 1A | A0A061GC06 | *Theobroma cacao* | 48 | Cyt/PM | ATP catabolic process |
| Regulatory particle triple-A ATPase 3 | Q9SEI4 | *Arabidopsis thaliana* | 46 | PM/Plamosdemos/Cyt/Cell wall | Root hair elongation/ER to Golgi vesicle-mediated transport |
| Rhamnose biosynthetic enzyme 1-like | A0A0A0L0D1 | *Cucumis sativus* | 75 | Cyt/Plasmodesmata | Metabolic process |
| Rhicadhesin receptor-like | A0A1S2XTZ4 | *Cicer arietinum* | 23 | Cell wall | Oxide-reduction |
| Ribosomal protein L16p/L10e family protein | A0A0R4J2Q7 | *Theobroma cacao* | 25 | Ribosome | Translation |
| Ribosomal protein L3 | Q6SKP4 | *Solanum lycopersicum* | 44 | Ribosome | Translation |
| Ribosomal protein S5 | A0A061F8V4 | *Theobroma cacao* | 94 | Chloroplast/PM/Cyt | GTP catabolic process |
| S-adenosyl-L-homocysteine hydrolase | P93253 | *Mesembryanthemum crystallinum* | 53 | TP/Golgi/Cyt/Plasmodesmata/PM | Water transport/Response to salt stress/Response to stress/Golgi organization |
| S-adenosyl-L-methionine:delta 24-sterol methyltransferase | P93852 | *Zea mays* | 39 | ER /Vacuole/ Plasmodesmata | Sterol biosynthesis |
| S-adenosylmethionine synthase | P16032 | *Mesembryanthemum crystallinum* | 20 | Cell wall/Chloroplast/Cyt | Response to stress |
| SAR-like protein | B7ZZP2 | *Zea mays* | 22 | Cyt membrane bounded vesicle/Gol | Response to salt stress/ER to Golgi vesicle mediated transport |
| Sec23/Sec24 protein transport family protein | Q9LUG1 | *Arabidopsis thaliana* | 86 | Cyt/COPII vesicle coat | ER to Golgi vesicle mediated transport/Response to salt stress |
| Serine hydroxymethyltransferase 2 | C6ZJY7 | *Glycine max* | 55 | | Metabolic process |
| Serine protease inhibitor (SERPIN) family protein | A0A061FCG1 | *Theobroma cacao* | 43 | | |
| Serine/arginine-rich splicing factor RSZ22-like | A0A0A0LEI3 | *Cucumis sativus* | 21 | | |
| Serine/threonine protein phosphatase 2A | Q39247 | *Arabidopsis thaliana* | 57 | Cytosol | Signaling |
| Serine/threonine-protein kinase HT1-like | UPI0002C35CEB | *Fragaria vesca* | 43 | Plasmodesmata/PM | Phosphorylation/Sterol Biosynthesis |
| Serine/threonine-protein kinase SRK2A | A0A6J5WDY4 | *Prunus mume* | 40 | | Phosphorylation |
| Serine/threonine-protein phosphatase 2A regulatory subunit A beta isoform | UPI00046E20A0 | *Prunus mume* | 66 | PM/Cell wall | Response to stress and stimulus |
| SH3 domain-containing protein | D7LA99 | *Arabidopsis lyrata* | 128 | Cyt/PM/Chloroplast | |
| SH3 domain-containing protein isoform 3 | A0A061GSY4 | *Theobroma cacao* | 130 | PM/Nucleus | |

| | | | | | |
|---|---|---|---|---|---|
| Stem-specific protein TSJT1-like | A0A3Q7I7U0 | *Solanum lycopersicum* | 27 | Cyt/PM/Nucleus | Response to stress |
| Stress responsive protein | Q10M11 | *Zea mays* | 39 | | |
| Succinate dehydrogenase [ubiquinone] iron-sulfur subunit 2 | A0A1S2YYS3 | *Prunus mume* | 69 | Mitochondria | Metabolic process |
| Succinate dehydrogenase 1-1 isoform 1 | A0A061DFR8 | *Theobroma cacao* | 70 | Mitochondria | Metabolic process |
| Sucrose synthase | P13708 | *Glycine max* | 92 | | Metabolic and biosynthetic process |
| Superoxide dismutase [Cu-Zn] 1 | P93258 | *Mesembryanthemum crystallinum* | 15 | Cyt | Response to salt stress/Metabolic process |
| T-complex protein 1 subunit alpha | P28769 | *Arabidopsis thaliana* | 57 | Cyt | Protein catalitic process/Protein folding |
| T-complex protein 1 subunit beta | Q940P8 | *Arabidopsis thaliana* | 57 | Cyt | Protein folding |
| T-complex protein 1 subunit beta-like isoform X1 | K3XGI3 | *Setaria italica* | 57 | Cyt | Protein folding |
| T-complex protein 1 subunit delta | UPI00046E175D | *Prunus mume* | 58 | Cyt/Anchored to PM | Protein folding/Response to stress/Response to salt stress |
| T-complex protein 1 subunit epsilon-like | A0A3Q7GD89 | *Solanum lycopersicum* | 59 | Cyt | Protein folding |
| T-complex protein 1 subunit gamma-like | A0A1S2Z036 | *Cicer arietinum* | 60 | Cyt | Protein folding |
| T-complex protein 1 subunit theta | M7Z9B8 | *Triticum urartu* | 73 | Membrane/Plasmodesmata/Cyt | Protein catabolic process |
| TCP-1/cpn60 chaperonin family protein | A0A061DJE4 | *Theobroma cacao* | 61 | Cyt | Response to salt stress/Protein folding/ Glycolysis |
| TCP-1/cpn60 chaperonin family protein isoform 1 | A0A061G375 | *Theobroma cacao* | 59 | Cyt | Protein folding |
| Thioredoxin H-type 1 | P29449 | *Nicotiana tabacum* | 14 | Cyt | Metabolic process/Cell redox homeostasis |
| Thiosulfate/3-mercaptopyruvate sulfurtransferase 1, Mitochondrial | UPI00046DDC8F | *Prunus mume* | 35 | | |
| TPR repeat-containing protein | A0A061FDC1 | *Theobroma cacao* | 90 | PM | |
| Transaldolase | Q9FVH1 | *Solanum lycopersicum* | 48 | Mitochondria/Chloroplast | Response to stress/Metabolic process |
| Trans-cinnamate 4-monooxygenase | Q43054 | *Populus sieboldii x Populus grandidentata* | 58 | Cell wall/ER/PM | Response to stress |
| Transducin/WD40 domain-containing protein | F4J0P2 | *Arabidopsis thaliana* | 175 | Cyt/PM | |
| Translation elongation factor-1 alpha 2 | S4SU69 | *Larix kaempferi* | 49 | Cyt | Translation/GTP catabolic process |
| Translational activator GCN1-like | M7ZCJ0 | *Triticum urartu* | 278 | | |
| Translocation protein SEC63 homolog | UPI000221665C | *Zea mays* | 34 | ER/PM/Cyt membrane bounded ves | Post-traduccional protein modification |
| Translocon at the outer envelope membrane of chloroplasts 75-III | A0A061GVS3 | *Theobroma cacao* | 89 | Vacuolar membrane | Protein transport |
| Transmembrane 9 superfamily member 3-like | UPI00046D992A | *Prunus mume* | 67 | Integral to Golgi/Endosome | |
| Transmembrane 9 superfamily member 4-like | A0A1S2YQM7 | *Cicer arietinum* | 75 | Integral to membrane/Endosome | |
| Tubulin alpha-1 chain | Q6VAG1 | *Gossypium hirsutum* | 50 | Microtubules/Cytoskeleton/Cyt | GTP catabolic process/Protein polimerization |
| Tubulin alpha-3 chain | B9R6R7 | *Ricinus communis* | 50 | Microtubulo | Protein polimerization |
| Tubulin alpha-5 | A0A061GR28 | *Theobroma cacao* | 50 | Cell wall/PM/Plasmodesmata/Cyt | Protein polimerization |
| Tubulin beta-5 chain | Q43697 | *Zea mays* | 50 | PM/Cyt | Response to salt stress/Signaling /Golgi organization/Water transport |
| Tubulin beta-7 chain | P37832 | *Oryza sativa* | 50 | Vacuole/Chloroplast/PM | GTP catabolic process/Protein polimerization |
| Tubulin beta-8 chain | P29516 | *Arabidopsis thaliana* | 51 | Golgi/Microtubules/Membrane | Protein polimerization |
| Ubiquitin carboxyl-terminal hydrolase 12-like | UPI0002C35C2C | *Fragaria vesca* | 131 | | Protein catabolic process |
| Ubiquitin-40S ribosomal protein S27a-like | A0A6N2CCA5 | *Solanum lycopersicum* | 18 | Cyt | Translation |
| Ubiquitin-activating enzyme E1 1 like isoform X1 | UPI00046DF20F | *Prunus mume* | 123 | | Protein modification |
| Ubiquitin-specific protease family C19-related protein | A0A061G9T7 | *Theobroma cacao* | 52 | | |
| Uclacyanin 2 | O80517 | *Arabidopsis thaliana* | 20 | Anchored to PM/Plasmodesmata | Response to stress/Metabolic process |
| UDP-glucose 6-dehydrogenase 1-like | A0A3Q7F9B8 | *Solanum lycopersicum* | 53 | | Oxide-reduction |
| UDP-glucuronic acid decarboxylase 1-like | A0A3Q7I5T5 | *Solanum lycopersicum* | 49 | Golgi/Vacuole/PM | Metabolic process |
| Uncharacterized protein LOC101267991 | A0A3Q7GIR6 | *Solanum lycopersicum* | 64 | | PM/ER/Nucleus |
| Vacuolar ATP synthase subunit C (VATC) / V-ATPase C subunit | A0A061ED54 | *Theobroma cacao* | 42 | Proton transportin V type ATPase V1 domain | ATP hydrolysis coupled proteon transport |

| | | | | | |
|---|---|---|---|---|---|
| Vacuolar ATPase subunit d | B6VAX6 | *Vigna radiata* | 41 | Proton transportin V type ATPase V1 domain | ATP hydrolysis coupled proteon transport |
| Vacuolar protein sorting 13 | J7FHV5 | *Mesembryanthemum crystallinum* | 416 | Plasmodesmata/Vacuole/Golgi/PM | Root hair diferentiation |
| Vacuolar-sorting receptor 3-like | UPI0002C2DE9F | *Fragaria vesca* | 70 | Membrane/Golgi/Vacuole | |
| Valine--tRNA ligase-like | UPI00046DED65 | *Prunus mume* | 117 | | Translation |
| Vesicle-associated membrane protein 726 | Q9MAS5 | *Arabidopsis thaliana* | 25 | Plasmodesmata/PM/Endosome | Vesicle mediated transport/Protein trageting to PM |
| VHA-A | O23654 | *Arabidopsis lyrata* | 69 | TP/Golgi/Cyt/Plasmodesmata/PM | Golgi organization/Response to stress/Response to salt stress/ATPase activity |
| Villin 2 isoform 1 | A0A061EX55 | *Theobroma cacao* | 107 | | Actic filament assembly |
| Voltage dependent anion channel 1 | A0A061EEK9 | *Theobroma cacao* | 30 | Mitochondria | Transport |
| Voltage dependent anion channel 1 | A0A061EEK9 | *Theobroma cacao* | 30 | Mitochondria outer membrane | Transport |
| V-type proton ATPase catalytic subunit A | D7EYG6 | *Glycine max* | 69 | TP/Chloroplast/Golgi/PM/Plasmodes | Vacuolar organization/Response to stress/Protein catalitic proces |
| V-type proton ATPase subunit B | K3XWK8 | *Setaria italica* | 54 | Vacuole/PM/Chloroplast | Response to stress/ATP metabolic process |
| V-type proton ATPase subunit E | Q40272 | *Mesembryanthemum crystallinum* | 26 | PM/TP | Response to salt stress/Golgi organization |
| V-type proton ATPase subunit H | A0A498KKZ1 | *Malus domestica* | 51 | Golgi/TP/PM | Transport/Golgi/Response to salt/Golgi vesicle transport |

***Table S2. Proteins identified by LC-MS/MS in each PCM fraction isolate by FFZE from M. crystallinum root.***

*Blue represents presence, green represents absence for that protein in the display fraction in NT or T condition.*

| Uniprot | Name | Not treated (NT) | | | | | | Salt treated (T) | | | | | |
|---|---|---|---|---|---|---|---|---|---|---|---|---|---|
| | | 80 | 81 | 82 | 83 | 84 | 85 | 80 | 81 | 82 | 83 | 84 | 85 |
| Q39247 | Serine/threonine protein phosphatase 2A | Green | Green | Green | Green | Green | Green | Green | Blue | Blue | Green | Green | Green |
| P93852 | S-adenosyl-L-methionine:delta 24-sterol methyltransferase | Green | Green | Blue | Green | Green | Green | Green | Green | Blue | Blue | Blue | Blue |
| C1KG74 | 14-3-3 protein | Blue | Blue | Blue | Blue | Blue | Blue | Blue | Blue | Blue | Blue | Blue | Blue |
| E9KNA4 | 14-3-3 protein SGF14h | Green | Blue | Blue | Blue | Green | Green | Green | Blue | Blue | Green | Blue | Blue |
| Q42134 | 20S proteasome alpha subunit E2 | Green | Green | Green | Green | Green | Green | Green | Green | Green | Green | Blue | Green |
| A0A061ELR7 | 26S proteasome regulatory complex, Rpn2/Psmd1 subunit | Green | Blue | Blue | Blue | Blue | Green | Blue | Blue | Green | Blue | Green | Blue |
| P62302 | 40S ribosomal protein S13 | Green | Green | Blue | Blue | Green | Blue | Blue | Green | Blue | Green | Green | Blue |
| M7YGU5 | 40S ribosomal protein S23 | Green | Green | Green | Blue | Green | Blue | Green | Blue | Blue | Blue | Blue | Blue |
| O64650 | 40S ribosomal protein S27-1 | Blue | Blue | Blue | Blue | Blue | Blue | Green | Green | Blue | Green | Green | Green |
| B4G194 | 40S ribosomal protein SA | Blue | Green | Blue | Green | Green | Blue | Green | Blue | Blue | Blue | Blue | Green |
| A0A061E919 | 51 kDa subunit of complex I | Blue | Blue | Blue | Blue | Blue | Blue | Blue | Blue | Blue | Blue | Blue | Blue |
| K7XKT4 | 60S ribosomal protein L1 | Blue | Blue | Blue | Blue | Green | Green | Blue | Blue | Blue | Blue | Blue | Blue |
| B6SJV1 | 60S ribosomal protein L17 | Blue | Green | Green | Blue | Green | Green | Green | Green | Green | Green | Green | Green |
| K7VP92 | 60S ribosomal protein L37a | Green | Green | Green | Green | Green | Green | Green | Green | Green | Blue | Green | Green |
| F4YFF3 | 60S ribosomal protein L9, partial | Green | Green | Green | Blue | Green | Green | Green | Green | Green | Green | Green | Green |
| F4JCC9 | AAA-type ATPase family protein | Blue | Blue | Blue | Green | Green | Green | Blue | Green | Blue | Green | Green | Green |
| Q42093 | ABC transporter C family member 2 | Blue | Green | Green | Green | Green | Green | Green | Green | Green | Green | Green | Green |
| L0E7F9 | Acetyl-CoA carboxylase beta subunit | Green | Green | Green | Green | Green | Green | Green | Green | Blue | Blue | Green | Blue |
| A0A061GCM9 | Adaptin family protein isoform 1 | Green | Blue | Blue | Blue | Green | Blue | Blue | Blue | Blue | Blue | Blue | Blue |
| P36397 | ADP-ribosylation factor 1 | Blue | Blue | Blue | Blue | Blue | Blue | Blue | Blue | Blue | Blue | Blue | Blue |
| P40940 | ADP-ribosylation factor 3 | Green | Green | Green | Blue | Green | Blue | Blue | Blue | Blue | Green | Blue | Blue |
| H1AD89 | ADP-ribosylation factor-like 8 | Blue | Blue | Blue | Green | Green | Green | Blue | Blue | Green | Green | Green | Green |
| A0A178UD24 | Aldolase-type TIM barrel family protein | Blue | Green | Blue | Blue | Green | Blue | Green | Green | Green | Blue | Green | Green |
| M1TCB6 | Alpha-tubulin 1 | Blue | Blue | Blue | Blue | Blue | Blue | Blue | Blue | Blue | Blue | Blue | Blue |
| S8CDY1 | Alpha-tubulin 3 | Blue | Blue | Blue | Blue | Green | Blue | Blue | Blue | Blue | Blue | Blue | Blue |
| G3E8E1 | Aquaporin 7 | Green | Green | Green | Green | Green | Green | Blue | Blue | Blue | Blue | Blue | Blue |
| P61837 | Aquaporin PIP1;1 | Blue | Blue | Blue | Blue | Blue | Blue | Blue | Blue | Blue | Blue | Blue | Blue |
| M8BEA3 | Aquaporin PIP1;5 | Blue | Blue | Blue | Blue | Blue | Blue | Blue | Blue | Blue | Blue | Blue | Blue |
| M1F3S6 | Aquaporin PIP2;1 | Blue | Green | Blue | Green | Green | Blue | Blue | Blue | Blue | Blue | Blue | Blue |

| | |
|---|---|
| M7YU72 | Aquaporin PIP2;5 |
| B4G031 | Ascorbate peroxidase2 |
| A0A061DZI3 | ATP binding cassette subfamily B4 isoform 1 |
| M7Y940 | ATP synthase subunit beta, mitochondrial |
| A0A061ETA0 | ATPase, AAA-type, CDC48 protein isoform 1 |
| UPI00046DD172 | ATP synthase subunit delta |
| J9RVH4 | Beta-tubulin, partial |
| A0A0D3DFE2 | Mitochondrial pyruvate carrier 2-like |
| A0A061FL71 | BR-signaling kinase 1 |
| UPI0001D464F1 | C2 domain-containing family protein |
| O23959 | Ca+2-binding EF hand protein |
| A0A061F1V7 | Calcium-binding EF-hand family protein isoform 1 |
| F4JTL3 | Calcium-dependent protein kinase 26 |
| M8AMH4 | Calcium-transporting ATPase 9 |
| Q4W5U7 | Calnexin-like protein precursor |
| S8CMU9 | Calnexin-like protein, partial |
| A0A762 | Calreticulin-1 precursor |
| Q9LZQ4 | Carbohydrate-binding-like fold-containing protein |
| O48947 | Cellulose synthase A catalytic subunit 2 [UDP-forming] |
| P29185 | Chaperonin CPN60-1, mitochondrial precursor |
| M8B4A2 | Chemocyanin |
| G3D5B5 | Cinnamate 4-hydroxylase |
| A0A061DXT8 | Coatomer subunit delta |
| A0A061FT37 | Clathrin, heavy chain isoform 1 |
| UPI00046DF541 | Coatomer subunit alpha-1-like |
| A0A061GJB6 | Coatomer, beta subunit isoform 1 |
| A0A061G8M4 | Cullin-associated and neddylation dissociated |
| B6SL40 | D-3-phosphoglycerate dehydrogenase |
| A0A061GK01 | DEAD/DEAH box RNA helicase family protein isoform 1 |
| Q9SVA6 | Developmentally regulated G-protein |
| A0A061DGI3 | Dolichyl-diphosphooligosaccharide-protein glycosyltransferase |
| A0A072TIU5 | DUF3411 domain protein |
| S8CML0 | Elongation factor 1 gamma-like protein, partial |
| B4FNT1 | Elongation factor 1-delta 1 |

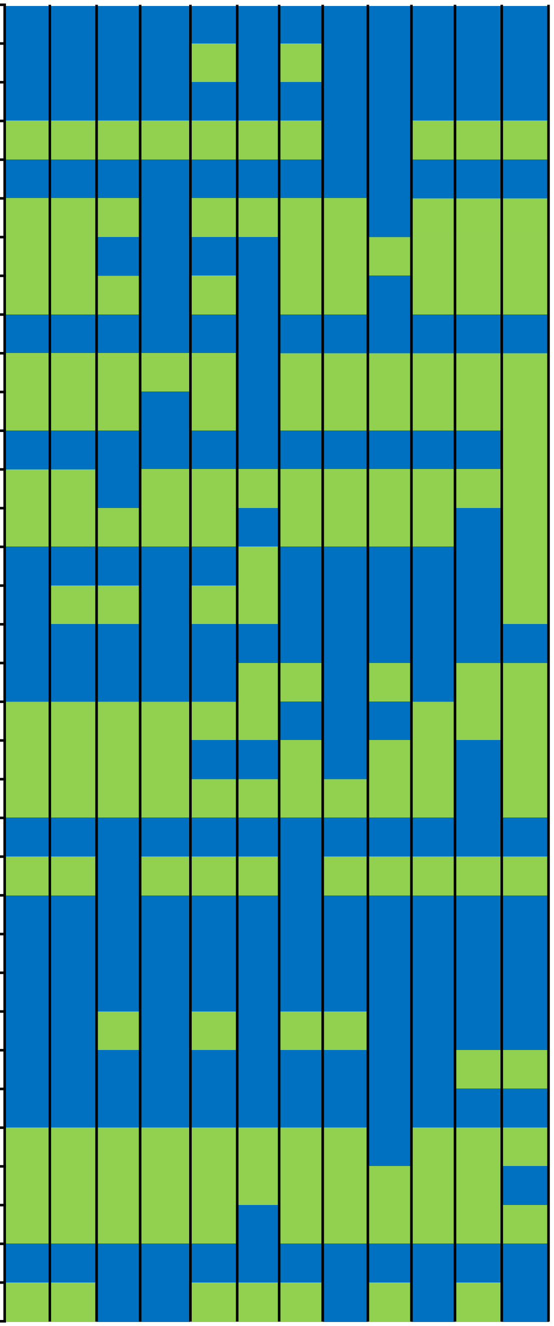

| | | |
|---|---|---|
| A0A061GXS4 | Endoplasmic reticulum [ER]-type calcium ATPase isoform 1 | 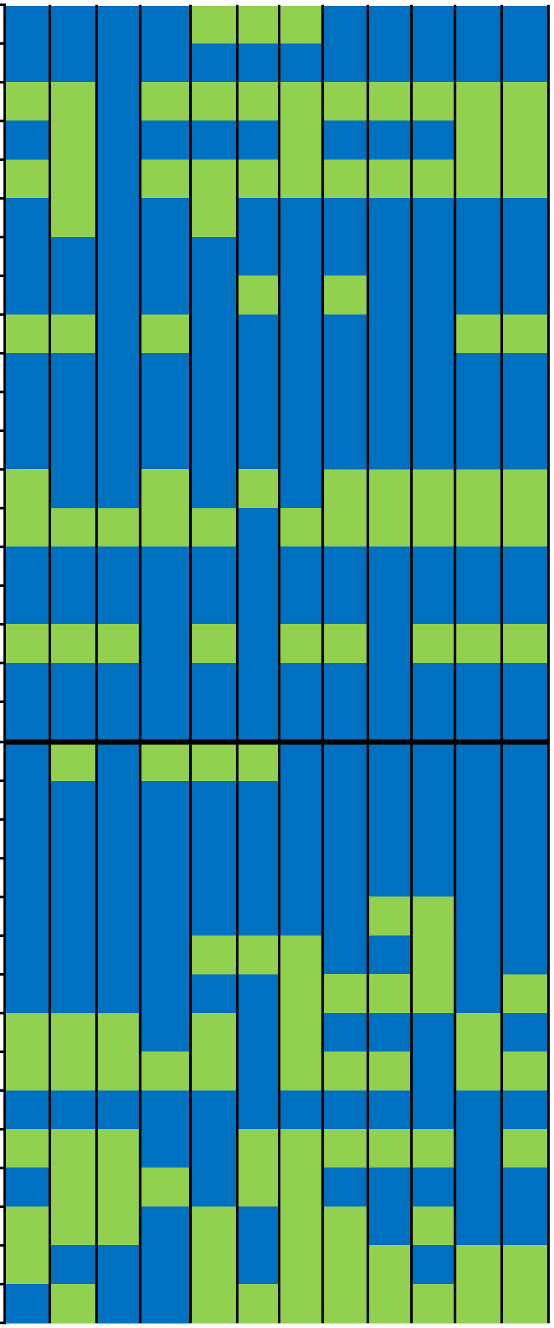 |
| UPI00046E14C3 | Endoplasmin homologo | |
| A0A061E0D5 | Eukaryotic aspartyl protease family protein isoform 1 | |
| B6U1I9 | Eukaryotic peptide chain release factor subunit 1-1 | |
| A0A061DJB3 | Eukaryotic translation initiation factor 2 family protein isoform 1 | |
| O49160 | Eukaryotic translation initiation factor 3 subunit C | |
| A0A061GKK2 | Eukaryotic translation initiation factor 4A1 | |
| Q9SMV6 | Exportin 1A | |
| M8BUY6 | Fatty acyl-CoA synthetase A | |
| M7ZGS6 | Fructose-bisphosphate aldolase cytoplasmic isozyme | |
| Q08069 | 40S ribosomal protein S8 | |
| UPI00046DFF87 | ABC transporter B family member 15 | |
| M4SIP8 | Glyceraldehyde-3-phosphate dehydrogenase | |
| A0A1S3B0B9 | Guanylate binding protein 2 | |
| A0A061H032 | Heat shock cognate protein 70-1 | |
| F4Y5A9 | Heat shock protein 81-3 | |
| F4JFN3 | Heat shock protein 89-1 | |
| K9JHZ2 | Heat shock protein 90-2 | |
| O03986 | Heat shock protein 90-4 | |
| P59259 | Histone H4 | |
| Q9FM19 | Hypersensitive-induced response protein 1 | |
| M0ZPM8 | Hypersensitive-induced response protein 2-like | |
| A0A061EEK9 | Voltage dependent anion channel 1 | |
| O22478 | Importin subunit alpha | |
| M7YB37 | Importin subunit beta-1 | |
| Q945K7 | Isocitrate dehydrogenase [NAD] regulatory subunit 3 | |
| A0A061DS88 | Ketol-acid reductoisomerase | |
| A0A061G6D0 | Kinase protein with tetratricopeptide repeat domain isoform 1 | |
| A0A061GN08 | Leucine-rich repeat protein kinase family protein | |
| Q94F62 | Leu-rich receptor Serine/threonine protein kinase BAK1 | |
| Q9M028 | LMBR1-like membrane protein | |
| S8CZD8 | Long chain acyl-CoA synthetase 9 isoform 1 | |
| Q94AH9 | Mediator of RNA polymerase II transcription subunit 36a | |
| A0A061ER94 | Metallopeptidase M24 family protein isoform 1 | |

| | |
|---|---|
| I6YI03 | Mitochondrial malate dehydrogenase |
| R7WA10 | Mitochondrial-processing peptidase subunit alpha |
| A0A061GTA5 | Myosin-related isoform 1 |
| Q42577 | NADH dehydrogenase (ubiquinone) Fe-S protein 7 |
| G9JLP7 | NADH dehydrogenase subunit 5 |
| G9JLR7 | NADH dehydrogenase subunit 7 |
| R4I225 | NADH dehydrogenase subunit 9 |
| T1RU75 | N-ethylmaleimide sensitive fusion protein |
| M7YQT8 | Nuclease domain-containing protein 1 |
| A0A0R4J410 | Obg-like ATPase 1-like |
| A0A061FEG6 | PAM domain (PCI/PINT associated module) protein isoform 1 |
| A0A178VIS2 | Peptide chain release factor eRF subunit 1 |
| M7ZBJ0 | Peroxidase 1 |
| R7VZF2 | Phosphate carrier protein |
| R0F3C0 | Phosphoenolpyruvate carboxykinase 1 |
| A0A061FI02 | Phosphoglycerate kinase isoform 1 |
| A0A061DGE7 | Phospholipase D delta isoform 1 |
| K4BDP2 | Phosphorus transporter |
| M1EY53 | **Plasma intrinsic protein PIP1.2** |
| M8AIK4 | Plasma membrane ATPase |
| K7TH91 | Plasma membrane H+-ATPase |
| UPI0002215CE2 | Plasma membrane H+-transporting ATPase-like protein |
| A0A061GNG0 | Pleiotropic drug resistance 12 |
| A0A0A0LJT3 | 26S protease regulatory subunit 6A homolog A-like |
| A0A0A0LNP0 | 26S protease regulatory subunit S10B homolog B-like |
| UPI0002C338E5 | 26S proteasome non-ATPase regulatory subunit 2 1A-like |
| I1J8K2 | 26S proteasome non-ATPase regulatory subunit 4 homolog isofor |
| A0A6J5UYR1 | 26S proteasome non-ATPase regulatory subunit 6 homolog |
| A0A3Q7HYJ1 | 26S proteasome regulatory subunit 4 homolog A-like |
| A0A0A0LHQ6 | 40S ribosomal protein S3-3-like |
| A0A0A0L4P6 | 40S ribosomal protein S3a-like |
| A0A1S2XMM1 | 40S ribosomal protein S4-2-like |
| I1L2H4 | 40S ribosomal protein S7 |
| A0A0A0L5R5 | 40S ribosomal protein S6-like |

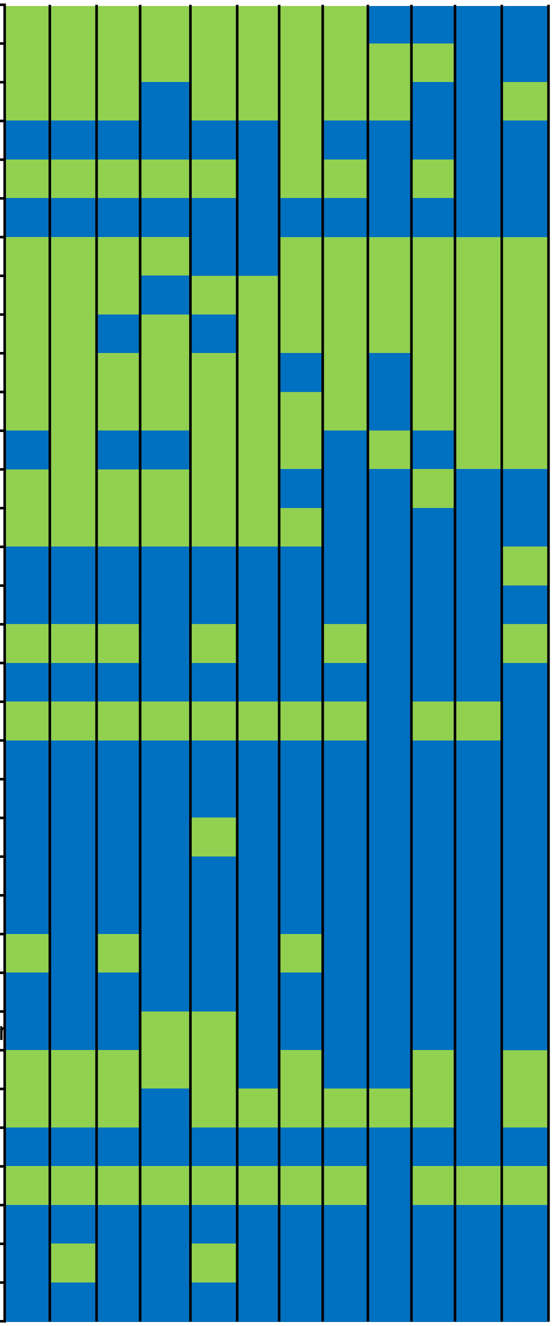

| | |
|---|---|
| UPI00046E0406 | 40S ribosomal protein S8 |
| UPI0002C3142A | 60S ribosomal protein L12-like |
| UPI0002767146 | 60S ribosomal protein L15-like |
| A0A3Q7I5W4 | 60S ribosomal protein L18-like |
| UPI0002B4A10B | 60S ribosomal protein L18a-like |
| I1GLQ1 | 60S ribosomal protein L24-like |
| I1L7D4 | 60S ribosomal protein L30-like |
| A0A0A0L819 | 60S ribosomal protein L37-3-like |
| M1BV98 | 60S ribosomal protein L7-3-like |
| UPI00046DFF87 | 6-phosphofructokinase 6-like |
| UPI0002767F3D | ABC transporter B family member 15-like |
| A0A3Q7I6V8 | ABC transporter C family member 4 |
| A0A2I4FMJ1 | ABC transporter F family member 4 |
| B4FPG2 | Actin-1 |
| K3ZQQ2 | Acyl-CoA dehydrogenase family member 10-like |
| A0A1D6E5Z6 | Adenosine kinase 2-like isoform X2 |
| A0A1S2YWC7 | ADP,ATP carrier protein 1, mitochondrial-like isoform X1 |
| A0A1S2YJ61 | ADP,ATP carrier protein 3, mitochondrial-like |
| UPI00046DBC20 | AP-1 complex subunit mu-1-like |
| UPI0002C30E13 | AP-2 complex subunit alpha-2-like |
| UPI000276C248 | AP-2 complex subunit mu-like |
| UPI0002BC88E7 | Aspartic proteinase-like |
| UPI0002C2F91D | ATP synthase subunit beta, mitochondrial-like |
| A0A0A0KRL4 | ATP synthase subunit d, mitochondrial-like |
| UPI0002767FD0 | ATPase family AAA domain-containing protein 3-B-like |
| A0A6N2C782 | ATP-dependent Clp protease ATP-binding subunit clpA homolog C |
| A0A0A0KH94 | ATP-dependent zinc metalloprotease FTSH 10 |
| A0A6N2BQ61 | Bifunctional polymyxin resistance protein ArnA-like |
| A0A1S2XPV0 | Brefeldin A-inhibited guanine nucleotide-exchange protein 2-like |
| A0A1S2XM13 | Calcium-dependent protein kinase 9-like |
| 460375325 | Calcium-transporting ATPase 2 |
| K3Y6F3 | Calnexin homolog |
| UPI0002C37537 | Calreticulin-like |
| A0A0A0KUY7 | Cation/H(+) antiporter 18-like |

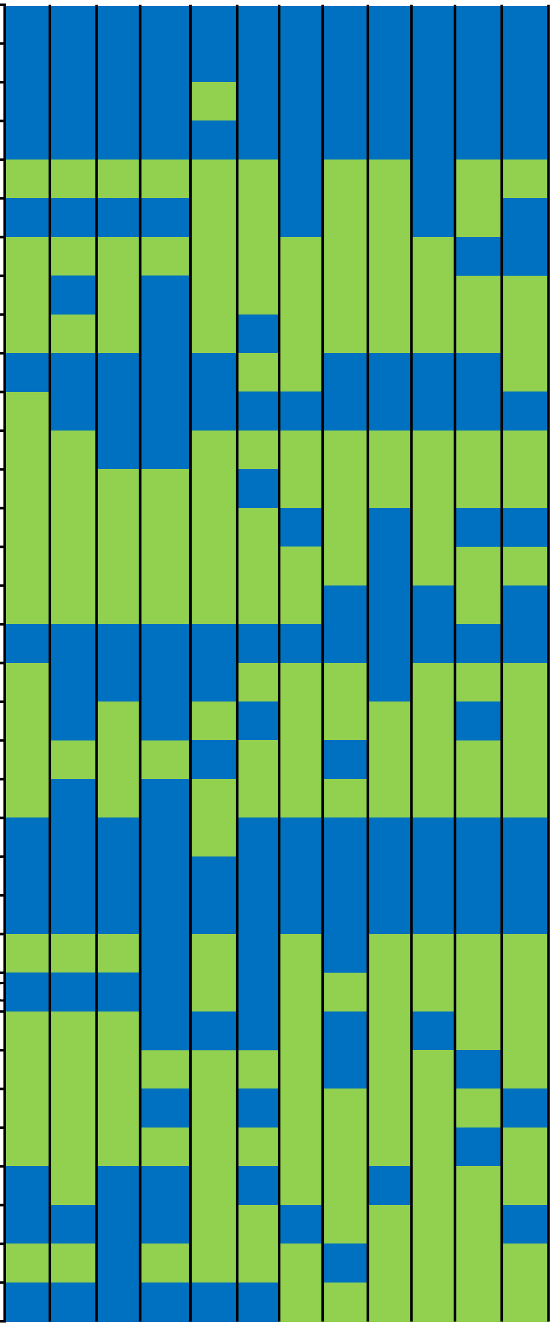

| | |
|---|---|
| UPI0002C3639C | Clathrin heavy chain 1-like |
| A0A3Q7HNN1 | Clathrin heavy chain 1-like |
| A0A0A0K130 | Coatomer subunit alpha-1-like |
| UPI0002C3407C | Coatomer subunit alpha-1-like |
| A0A0A0LCU8 | Coatomer subunit beta-1-like |
| UPI000276943D | Coatomer subunit beta-2-like |
| UPI00027689D6 | Coatomer subunit gamma-2-like |
| UPI0002C32A7C | D-3-phosphoglycerate dehydrogenase, chloroplastic-like |
| A0A1S2YC83 | DEAD-box ATP-dependent RNA helicase 52C-like |
| UPI0002B4BA64 | DEAD-box ATP-dependent RNA helicase 8-like |
| UPI0002C306D2 | Delta(24)-sterol reductase-like |
| A0A3Q7ICL0 | Diacylglycerol kinase iota-like |
| UPI0002C33D25 | DNA damage-inducible protein 1-like |
| UPI0002BC90E9 | DnaJ homolog subfamily C GRV2-like |
| A0A3Q7J3H2 | Dolichyl-diphosphooligosaccharide--protein glycosyltransferase su |
| UPI0002209ABB | DUF21 domain-containing protein At4g14240-like |
| A0A314YPJ6 | E3 ubiquitin-protein ligase UPL2-like |
| UPI00046DD67D | Elongation factor 1-alpha |
| K3XVC0 | Endoplasmin homolog |
| A0A3Q7HMP2 | Eukaryotic initiation factor 4A-2-like |
| K3Y6E4 | Eukaryotic peptide chain release factor subunit ERF3A-like |
| M5X0K8 | Eukaryotic translation initiation factor 2 subunit alpha-like |
| UPI0002C2DDD4 | Eukaryotic translation initiation factor 3 subunit A-like |
| UPI00046E04E2 | Eukaryotic translation initiation factor 3 subunit B-like |
| A0A1S2XCE8 | Eukaryotic translation initiation factor 3 subunit D-like |
| A0A0A0L0F9 | Eukaryotic translation initiation factor 3 subunit E-like |
| M1BYD6 | Eukaryotic translation initiation factor 3 subunit H-like |
| A0A1S2XDH1 | Flavoprotein WrbA-like |
| UPI00046DD6EA | Fructose-bisphosphate aldolase cytoplasmic isozyme |
| A0A0A0KWN9 | Gamma carbonic anhydrase 1, mitochondrial-like |
| A0A0A0LR76 | Gamma carbonic anhydrase-like 2, mitochondrial-like |
| UPI0002768717 | Gamma carbonic anhydrase-like 2, mitochondrial-like |
| UPI00027688C2 | Glucose-6-phosphate 1-dehydrogenase, cytoplasmic isoform 1 |
| UPI00046E1A6C | Glutamate decarboxylase 1 |

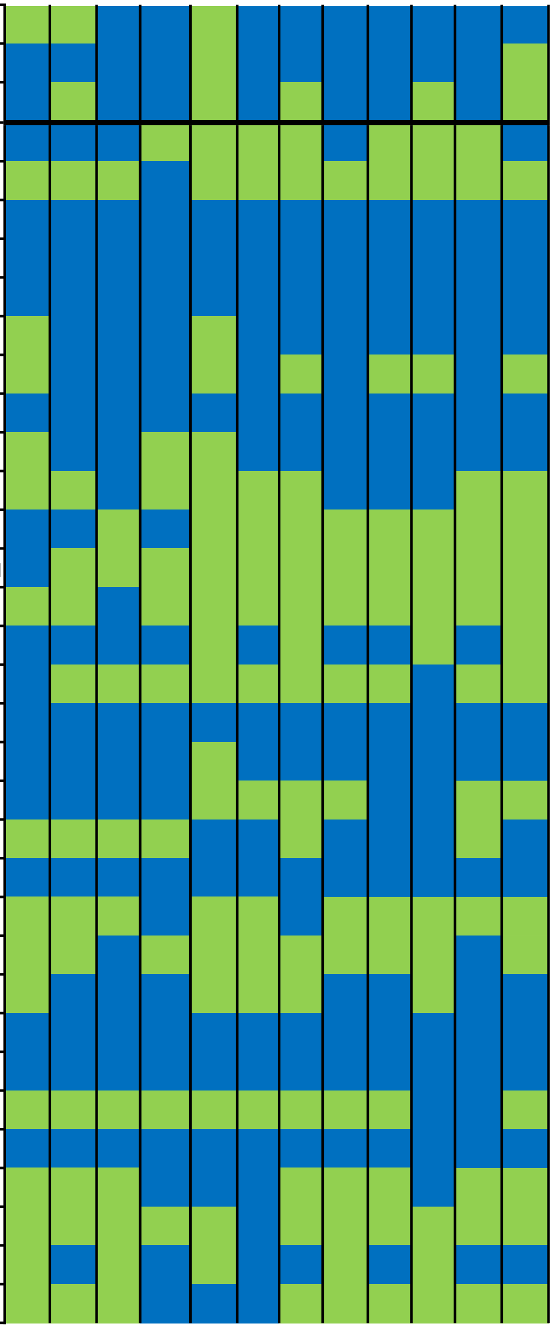

| | |
|---|---|
| UPI0002769AB4 | Glutamine synthetase |
| A0A0A0K8C1 | Glyceraldehyde-3-phosphate dehydrogenase-like isoform 1 |
| A0A3Q7HK84 | GTP-binding protein SAR1A-like |
| A0A3Q7J9R5 | Heat shock 70 kDa protein 15-like |
| A0A1S2Y850 | Heat shock cognate protein 80-like |
| UPI0002C303FE | Heat shock cognate protein 90-2 |
| A0A3Q7EFC3 | Importin subunit alpha-1a-like |
| A0A3Q7EFC3 | Importin subunit alpha |
| K4A4S4 | Intron-binding protein aquarius-like |
| A0A3Q7XWW5 | Leucine-rich repeat receptor-like serine/threonine-protein kinase B |
| K3YGV3 | L-gulonolactone oxidase-like |
| P83970 | Plasma membrane ATPase 4-like |
| A0A1S2Z7E5 | Probable calcium-binding protein CML49-like |
| UPI0002C35E23 | Pyrophosphate-energized vacuolar membrane proton pump-like |
| B6TGD4 | F1F0- ATPase inhibitor protein |
| UPI0002768EA1 | Membrane steroid-binding protein 2-like |
| A0A3Q7EA80 | Mitochondrial outer membrane protein porin of 36 kDa-like |
| A0A0A0LIM9 | Mitochondrial Rho GTPase |
| A0A3Q7J3G4 | Mitochondrial-processing peptidase subunit alpha-like |
| A0A1S2Z641 | Monocopper oxidase-like protein SKU5-like |
| UPI000276C5F7 | NAD(P)H dehydrogenase B1, mitochondrial-like |
| A0A1S2XV81 | NADH dehydrogenase [ubiquinone] 1 alpha subcomplex subunit 9 |
| UPI0002C2E6B3 | NADH dehydrogenase [ubiquinone] iron-sulfur protein 1 |
| A0A5E4FKR1 | Nascent polypeptide-associated complex subunit alpha-like protei |
| A0A1S2YCV8 | Neutral alpha-glucosidase AB-like |
| UPI0002B48A57 | Nicalin-like |
| A0A1S2XML4 | Niemann-Pick C1 protein-like |
| J3N9D2 | Non-specific phospholipase C4-like |
| A0A3Q7J2T2 | Nucleosome assembly protein 1-like 1-like isoform 1 |
| UPI0002C2E42C | Patellin-3-like |
| A0A1S2YYU4 | Peroxidase 72-like |
| UPI00046D98F7 | Phospho-2-dehydro-3-deoxyheptonate aldolase 1 |
| A0A1S2YPD3 | Plasma membrane ATPase 4-like |
| A0A3Q7H652 | Plasma membrane ATPase 4-like |

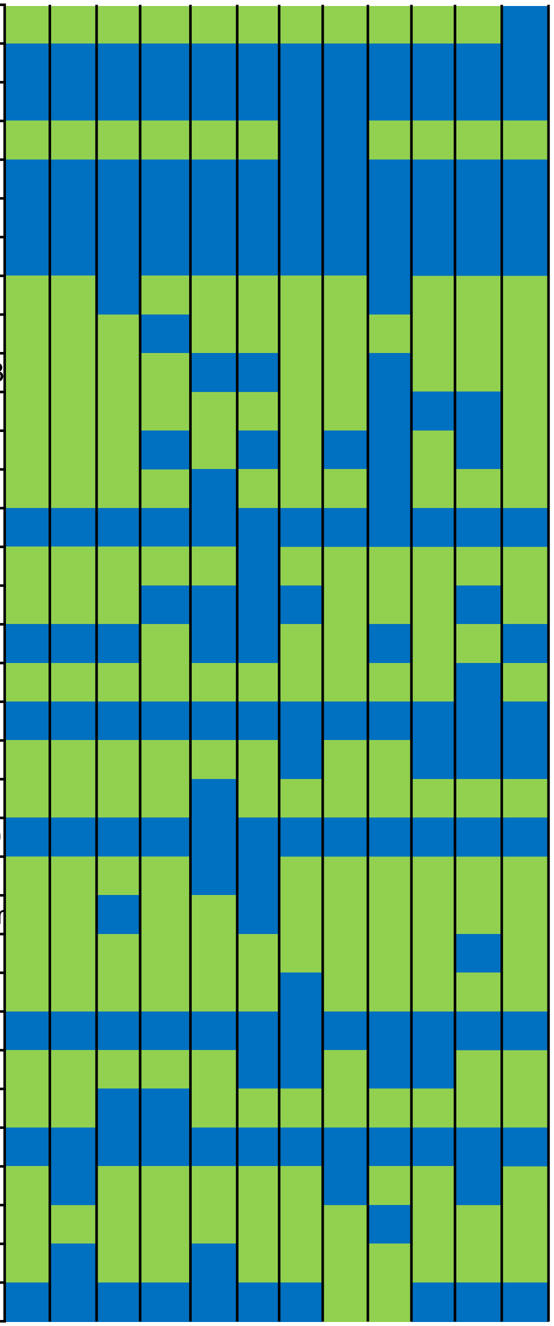

| | |
|---|---|
| A0A1S2YIR6 | Polyadenylate-binding protein |
| A0A3Q7JU30 | Potassium transporter 4-like |
| A0A3Q7I1D8 | Pre-mRNA-processing-splicing factor 8-like |
| UPI00046DC9CC | Alpha,alpha-trehalose-phosphate synthase [UDP-forming] |
| A0A6A3BQM0 | **Aquaporin PIP1;2** |
| UPI00046DDEA9 | **Aquaporin PIP1;4** |
| A0A222C4Z4 | **Aquaporin PIP1;4** |
| UPI000276C169 | Importin-7 homolog |
| A0A1S2XWA4 | Inactive serine/threonine-protein kinase scy1-like isoform X1 |
| UPI000276B496 | LRR receptor-like serine/threonine-protein kinase At1g56130-like |
| A0A1S2XIY9 | Methyltransferase PMT2-like isoform X1 |
| A0A0A0L0I1 | Mitochondrial-processing peptidase subunit beta-like |
| UPI0002B487B6 | NADH dehydrogenase [ubiquinone] 1 alpha subcomplex subunit 5 |
| UPI00046E2350 | Nucleolar protein 5-2 |
| UPI00046DA708 | Protein phosphatase 2C 9 |
| A0A0A0L0D1 | Rhamnose biosynthetic enzyme 1-like |
| A0A3Q7F9B8 | UDP-glucose 6-dehydrogenase 1-like |
| A0A0A0KM77 | Inactive leucine-rich repeat receptor-like protein kinase IMK2-like |
| A0A0R4J321 | Proteasome subunit beta type-6 |
| UPI00046DD765 | Root hair defective 3-like |
| A0A1S3CCL1 | Protein transport protein SEC23-like |
| UPI0002C2DF41 | Protein transport protein Sec24-like At3g07100-like |
| A0A3Q7IAQ0 | Protein transport protein Sec61 subunit alpha-like |
| A0A0A0KZ03 | Mitochondrial 2-oxoglutarate/malate carrier protein-like |
| UPI0002B45BAC | Pyruvate kinase, cytosolic isozyme-like |
| UPI0002C2EE30 | Ras-related protein RABA1f-like |
| A0A3Q7EQ24 | Ras-related protein RABA2a-like |
| A0A0V0HPF5 | Ras-related protein RABG3f-like isoform 1 |
| K3Z3U9 | Receptor-like protein kinase HERK 1-like |
| A0A1S2XTZ4 | Rhicadhesin receptor-like |
| A0A0A0LEI3 | Serine/arginine-rich splicing factor RSZ22-like |
| UPI0002C35CEB | Serine/threonine-protein kinase HT1-like |
| A0A6J5WDY4 | Serine/threonine-protein kinase SRK2A |
| A0A3Q7I7U0 | Stem-specific protein TSJT1-like |

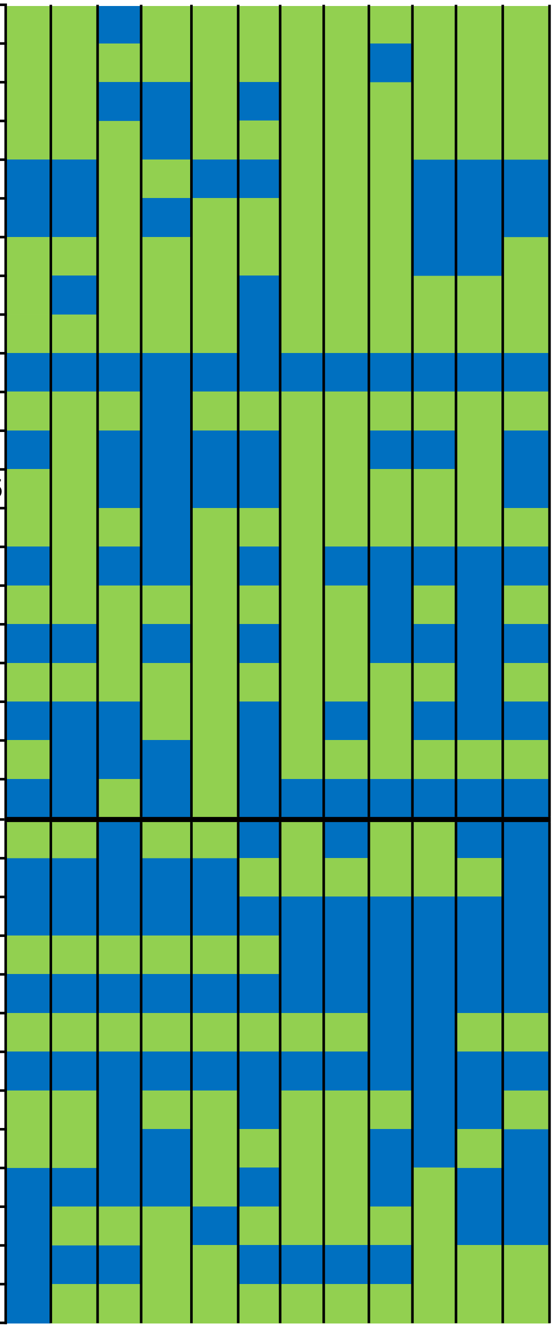

| | |
|---|---|
| A0A1S2YYS3 | Succinate dehydrogenase [ubiquinone] iron-sulfur subunit 2 |
| Q940P8 | T-complex protein 1 subunit beta |
| K3XGI3 | T-complex protein 1 subunit beta-like isoform X1 |
| UPI00046E175D | T-complex protein 1 subunit delta |
| A0A3Q7GD89 | T-complex protein 1 subunit epsilon-like |
| A0A1S2Z036 | T-complex protein 1 subunit gamma-like |
| UPI00046DDC8F | Thiosulfate/3-mercaptopyruvate sulfurtransferase 1, Mitochondrial |
| M7ZCJ0 | Translational activator GCN1-like |
| UPI000221665C | Translocation protein SEC63 homolog |
| UPI00046D992A | Transmembrane 9 superfamily member 3-like |
| A0A1S2YQM7 | Transmembrane 9 superfamily member 4-like |
| UPI0002C35C2C | Ubiquitin carboxyl-terminal hydrolase 12-like |
| A0A6N2CCA5 | Ubiquitin-40S ribosomal protein S27a-like |
| A0A3Q7I5T5 | UDP-glucuronic acid decarboxylase 1-like |
| A0A3Q7GIR6 | Uncharacterized protein LOC101267991 |
| UPI0002C2DE9F | Vacuolar-sorting receptor 3-like |
| UPI00046DED65 | Valine--tRNA ligase-like |
| K3XWK8 | V-type proton ATPase subunit B |
| A0A498KKZ1 | V-type proton ATPase subunit H |
| Q9SID0 | Fructokinase-1 |
| A0A061F4N1 | Proteasome activating protein 200 isoform 1 |
| S8C641 | Pto kinase interactor 1 |
| K7WJW2 | 40S ribosomal protein S9 |
| Q9ZT91 | Elongation factor Tu |
| M7ZR94 | Glycerophosphoryl diester phosphodiesterase 1 |
| N1QWM1 | LRR receptor-like serine/threonine-protein kinase |
| A0A061FDC1 | TPR repeat-containing protein |
| O65744 | Rab GDP dissociation inhibitor alpha-like |
| P92963 | RAB GTPase homolog B1C |
| Q38902 | Rac-like GTP-binding protein ARAC1 |
| P93214 | 14-3-3 protein 9 |
| O65352 | 14-3-3-like protein |
| O49996 | 14-3-3-like protein D |
| P93259 | 14-3-3-like protein |

| | |
|---|---|
| Q42908 | 2,3-bisphosphoglycerate-independent phosphoglycerate mutase |
| P17093 | 40S ribosomal protein S11 |
| A0A6P5TX25 | 40S ribosomal protein S14-3 |
| P93263 | 5-methyltetrahydropteroyltriglutamate--homocysteine methyltransfe |
| P50346 | 60S acidic ribosomal protein P0 |
| Q0DK10 | 60S ribosomal protein L11 |
| I1N527 | 60S ribosomal protein L23a |
| A0A498JM53 | 60S ribosomal protein L26-1 |
| Q0JGY1 | 60S ribosomal protein L5-1 |
| P34091 | 60S ribosomal protein L6 |
| P29766 | 60S ribosomal protein L8 |
| P30164 | Actin-1 |
| Q10DV7 | Actin-1 |
| P93253 | S-adenosyl-L-homocysteine hydrolase |
| Q08480 | Adenylate kinase 4 |
| P13547 | ATP synthase subunit 9 |
| P05492 | ATP synthase subunit alpha, mitochondrial |
| P17614 | ATP synthase subunit beta, mitochondrial |
| P26360 | ATP synthase subunit gamma, mitochondrial |
| P04464 | Calmodulin |
| Q01297 | Catalase isozyme 1 |
| Q2RBN7 | Clathrin heavy chain 1 |
| Q0WNJ6 | Clathrin heavy chain 1 |
| O65361 | Delta-1-pyrroline-5-carboxylate synthase |
| P43644 | DnaJ protein homolog ANJ1 |
| M5VHG2 | Dynamin-2B |
| O23755 | Elongation factor 2 |
| Q43130 | Enolase |
| P24922 | Eukaryotic translation initiation factor 5A-2 |
| P29356 | Fructose-bisphosphate aldolase, cytoplasmic isozyme |
| P17878 | Glyceraldehyde-3-phosphate dehydrogenase, cytosolic |
| K3XLR2 | GTP-binding nuclear protein Ran-3 |
| P93398 | Guanine nucleotide-binding protein subunit beta-2 |
| O24076 | Guanine nucleotide-binding protein subunit beta-like |

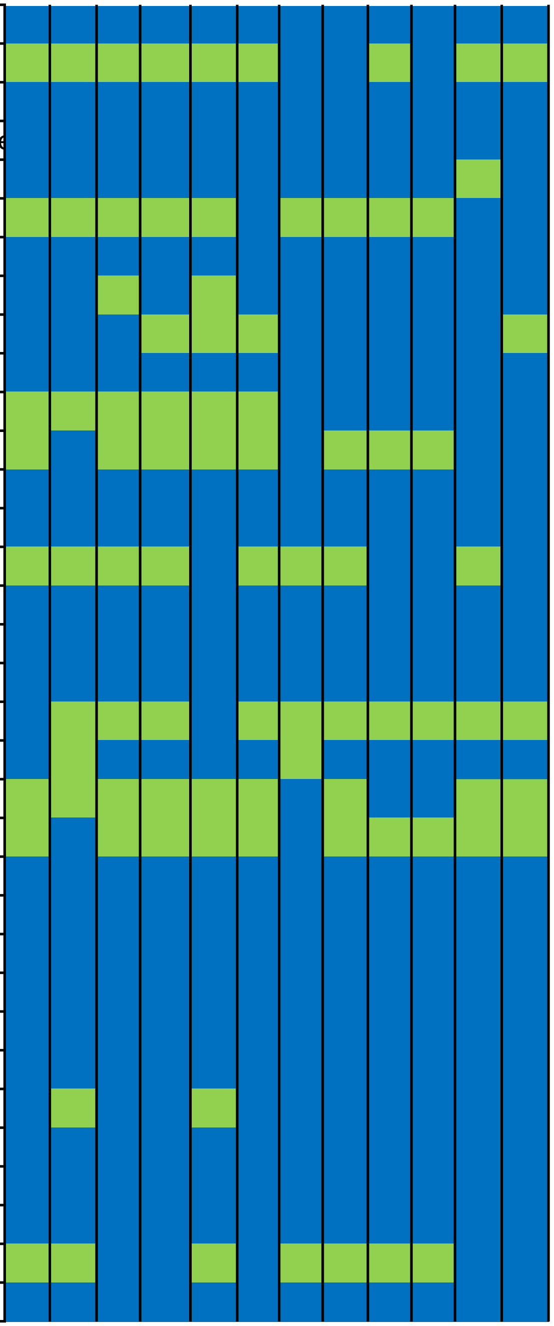

| | |
|---|---|
| A2WKT1 | Histone H2B.6 |
| Q40345 | Isocitrate dehydrogenase [NADP], chloroplastic |
| Q03684 | Luminal-binding protein 4 |
| Q42434 | Luminal-binding protein |
| O24047 | Malate dehydrogenase, cytoplasmic |
| Q42711 | Monodehydroascorbate reductase |
| Q43644 | NADH dehydrogenase [ubiquinone] iron-sulfur protein 1 |
| Q01148 | NADH-ubiquinone oxidoreductase chain 1 |
| O81372 | Nucleoside diphosphate kinase 1 |
| P16097 | Phosphoenolpyruvate carboxylase 2 |
| P93262 | Phosphoglucomutase, cytoplasmic |
| Q76CU2 | Pleiotropic drug resistance protein 1 |
| B6SHX9 | Histone H2A.6 |
| K4B7G2 | L-type lectin-domain containing receptor kinase S.5 |
| UPI00046DB4A1 | Probable methyltransferase PMT8 |
| Q9LEF0 | Probable phospholipid hydroperoxide glutathione peroxidase |
| P22200 | Pyruvate kinase, cytosolic isozyme |
| Q39433 | Ras-related protein RAB1BV |
| P16032 | S-adenosylmethionine synthase |
| O48660 | Putrescine aminopropyltransferase |
| P93258 | Superoxide dismutase [Cu-Zn] 1 |
| P28769 | T-complex protein 1 subunit alpha |
| P29449 | Thioredoxin H-type 1 |
| Q43054 | Trans-cinnamate 4-monooxygenase |
| Q6VAG1 | Tubulin alpha-1 chain |
| P37832 | Tubulin beta-7 chain |
| UPI00046DF20F | Ubiquitin-activating enzyme E1 1 like isoform X1 |
| Q40272 | V-type proton ATPase subunit E |
| A0A061GC06 | Regulatory particle triple-A 1A |
| Q9SEI4 | Regulatory particle triple-A ATPase 3 |
| A0A0R4J2Q7 | Ribosomal protein L16p/L10e family protein |
| Q6SKP4 | Ribosomal protein L3 |
| A0A061F8V4 | Ribosomal protein S5 |
| B7ZZP2 | SAR-like protein |

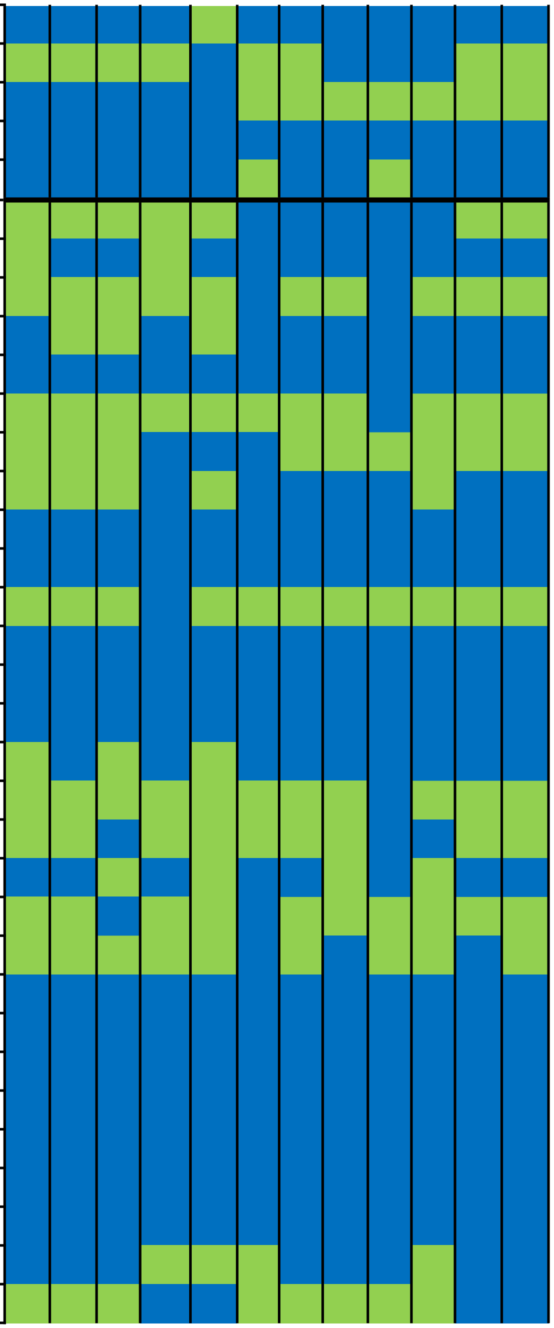

| | |
|---|---|
| Q9LUG1 | Sec23/Sec24 protein transport family protein |
| C6ZJY7 | Serine hydroxymethyltransferase 2 |
| A0A061FCG1 | Serine protease inhibitor (SERPIN) family protein |
| UPI00046E20A0 | Serine/threonine-protein phosphatase 2A regulatory subunit A beta |
| D7LA99 | SH3 domain-containing protein |
| A0A061GSY4 | SH3 domain-containing protein isoform 3 |
| Q10M11 | Stress responsive protein |
| A0A061DFR8 | Succinate dehydrogenase 1-1 isoform 1 |
| P13708 | Sucrose synthase |
| M7Z9B8 | T-complex protein 1 subunit theta |
| A0A061DJE4 | TCP-1/cpn60 chaperonin family protein |
| A0A061G375 | TCP-1/cpn60 chaperonin family protein isoform 1 |
| Q9FVH1 | Transaldolase |
| F4J0P2 | Transducin/WD40 domain-containing protein |
| S4SU69 | Translation elongation factor-1 alpha 2 |
| A0A061GVS3 | Translocon at the outer envelope membrane of chloroplasts 75-III |
| B9R6R7 | Tubulin alpha-3 chain |
| A0A061GR28 | Tubulin alpha-5 |
| Q43697 | Tubulin beta-5 chain |
| P29516 | Tubulin beta-8 chain |
| A0A061G9T7 | Ubiquitin-specific protease family C19-related protein |
| A0A061ED54 | Vacuolar ATP synthase subunit C (VATC) / V-ATPase C subunit |
| B6VAX6 | Vacuolar ATPase subunit d |
| J7FHV5 | Vacuolar protein sorting 13 |
| Q9MAS5 | Vesicle-associated membrane protein 726 |
| O23654 | VHA-A |
| A0A061EX55 | Villin 2 isoform 1 |
| A0A061EEK9 | Voltage dependent anion channel 1 |
| D7EYG6 | V-type proton ATPase catalytic subunit A |
| P26518 | Glyceraldehyde-3-phosphate dehydrogenase, cytosolic |
| O80517 | Uclacyanin 2 |

***Table S3. Salt up-regulated proteins in each PCM fractions from M. crystallinum root.***

*Protein name and  accesion numer are display.*

*p-Value and Fold change was obtain by a student's T-test according to the emPAI of each protein when comparing T/NT protein abundance n=4*

*Blue represents presence increase abundance, red decrease abundance, purple respresents proteins that appear for salt treatment and green proteins that disappear in the correponding analyze fraction.*

**Proteins that change in abundace in Fraction T80**

| Protein name | Uniprot Accession | p-Value | Fold change |
|---|---|---|---|
| Ca+2-binding EF hand protein | O23959 | 0.00068 | **INF** |
| T-complex protein 1 subunit epsilon-like | A0A3Q7GD89 | 0.033 | 5.6 |
| **Ras-related protein RABA1f-like** | UPI0002C2EE30 | 0.025 | 3.3 |
| Coatomer subunit delta | A0A061DXT8 | 0.031 | 2.8 |
| Serine hydroxymethyltransferase 2 | C6ZJY7 | 0.042 | 2.2 |
| Sucrose synthase | P13708 | 0.0033 | 2.1 |
| Enolase | Q43130 | 0.042 | 2.1 |
| Monodehydroascorbate reductase | Q42711 | 0.037 | 2 |
| TCP-1/cpn60 chaperonin family protein | A0A061DJE4 | 0.052 | 1.8 |
| **Clathrin heavy chain 1-like** | A0A3Q7HNN1 | 0.016 | 1.3 |
| **Aquaporin PIP2;5** | M7YU72 | 0.053 | 1.2 |
| 14-3-3 protein | C1KG74 | 0.044 | 0.8 |
| Glyceraldehyde-3-phosphate dehydrogenase | M4SIP8 | 0.013 | 0.7 |
| UDP-glucose 6-dehydrogenase 1-like | A0A3Q7F9B8 | 0.019 | 0.6 |
| Phosphoglycerate kinase isoform 1 | A0A061FI02 | 0.041 | 0.6 |
| Protein transport protein Sec61 subunit alpha-like | A0A3Q7IAQ0 | 0.019 | 0.3 |
| Beta-tubulin, partial | J9RVH4 | 0.026 | 0.3 |
| Nuclease domain-containing protein 1 | M7YQT8 | 0.027 | 0.3 |
| Superoxide dismutase [Cu-Zn] 1 | P93258 | 0.0005 | 0 |
| Cation/H(+) antiporter 18-like | A0A0A0KUY7 | 0.019 | 0 |

**Proteins that change in abundace in Fraction T81**

| Protein name | Uniprot Accession | p-Value | Fold change |
|---|---|---|---|
| Actin-1 | B4FPG2 | 0.00061 | **INF** |
| Histone H2A.6 | B6SHX9 | 0.011 | 3 |
| 2,3-bisphosphoglycerate-independent phosphoglycerate mu | Q42908 | 0.0056 | 2 |
| 14-3-3-like protein | P93259 | 0.043 | 1.9 |
| **Aquaporin PIP2;5** | M7YU72 | 0.025 | 1.6 |
| Heat shock protein 90-2 | K9JHZ2 | 0.004 | 0.8 |
| Heat shock protein 90-4 | O03986 | 0.042 | 0.7 |
| Elongation factor 1 gamma-like protein, partial | S8CML0 | 0.045 | 0.7 |
| RAB GTPase homolog B1C | P92963 | 0.014 | 0.6 |
| Transaldolase | Q9FVH1 | 0.02 | 0.5 |
| Pyrophosphate-energized vacuolar membrane proton pump | UPI0002C35E23 | 0.027 | 0.5 |
| Calcium-transporting ATPase 2 | 460375325 | 0.039 | 0.5 |
| TCP-1/cpn60 chaperonin family protein | A0A061DJE4 | 0.024 | 0.2 |
| Nuclease domain-containing protein 1 | M7YQT8 | 0.025 | 0.2 |

**Proteins that change in abundace in Fraction T82**

| Protein name | Uniprot Accession | p-Value | Fold change |
|---|---|---|---|
| Ubiquitin carboxyl-terminal hydrolase 12-like | UPI0002C35C2C | 0.0022 | **INF** |

| | | | |
|---|---|---|---|
| Myosin-related isoform 1 | A0A061GTA5 | 0.047 | 6.7 |
| Serine hydroxymethyltransferase 2 | C6ZJY7 | 0.0074 | 3.3 |
| Putrescine aminopropyltransferase | O48660 | 0.012 | 2.5 |
| Aspartic proteinase-like | UPI0002BC88E7 | 0.031 | 1.5 |
| Sucrose synthase | P13708 | 0.027 | 1.3 |
| Heat shock protein 90-4 | O03986 | 0.013 | 0.7 |
| Pto kinase interactor 1 | S8C641 | 0.0083 | 0.6 |
| ATP binding cassette subfamily B4 isoform 1 | A0A061DZI3 | 0.021 | 0.2 |

**Proteins that change in abundace in Fraction T83**

| Protein name | Uniprot Accession | p-Value | Fold change |
|---|---|---|---|
| Catalase isozyme 1 | Q01297 | 0.0097 | **INF** |
| Glycerophosphoryl diester phosphodiesterase 1 | M7ZR94 | 0.01 | **INF** |
| Mitochondrial malate dehydrogenase | I6YI03 | 0.036 | 5.8 |
| Plasma membrane ATPase 4-like | A0A1S2YPD3 | 0.018 | 4.4 |
| 20S proteasome alpha subunit E2 | Q42134 | 0.026 | 3.8 |
| Monocopper oxidase-like protein SKU5-like | A0A1S2Z641 | 0.006 | 3.3 |
| Voltage dependent anion channel 1 | A0A061EEK9 | 0.039 | 3 |
| Monodehydroascorbate reductase | Q42711 | 0.032 | 2.9 |
| Fructose-bisphosphate aldolase cytoplasmic isozyme | UPI00046DD6EA | 0.014 | 2.7 |
| **Aquaporin PIP2;5** | M7YU72 | 0.0039 | 2.5 |
| Serine hydroxymethyltransferase 2 | C6ZJY7 | 0.024 | 2.5 |
| **Aquaporin PIP1;2** | A0A6A3BQM0 | 0.021 | 2.3 |
| **Aquaporin PIP1;4** | A0A222C4Z4 | 0.053 | 2.3 |
| 60S ribosomal protein L37a | K7VP92 | 0.038 | 2.2 |
| Protein phosphatase 2C 9 | UPI00046DA708 | 0.049 | 2.2 |
| **Aquaporin PIP2;1** | M1F3S6 | 0.039 | 2.1 |
| **Ras-related protein RABG3f-like isoform 1** | A0A0V0HPF5 | 0.044 | 1.9 |
| Calnexin-like protein precursor | Q4W5U7 | 0.016 | 1.4 |
| Tubulin alpha-5 | A0A061GR28 | 0.036 | 0.8 |
| Alpha-tubulin 1 | M1TCB6 | 0.0046 | 0.7 |
| Coatomer subunit beta-1-like | A0A0A0LCU8 | 0.016 | 0.7 |
| Alpha-tubulin 3 | S8CDY1 | 0.034 | 0.7 |
| Uncharacterized protein LOC101267991 | A0A3Q7GIR6 | 0.046 | 0.7 |
| Tubulin beta-7 chain | P37832 | 0.017 | 0.6 |
| Tubulin beta-8 chain | P29516 | 0.029 | 0.6 |
| TCP-1/cpn60 chaperonin family protein | A0A061DJE4 | 0.038 | 0.6 |
| Eukaryotic translation initiation factor 3 subunit E-like | A0A0A0L0F9 | 0.041 | 0.6 |
| ATPase, AAA-type, CDC48 protein isoform 1 | A0A061ETA0 | 0.047 | 0.6 |
| NADH dehydrogenase [ubiquinone] 1 alpha subcomplex su | A0A1S2XV81 | 0.0029 | 0.5 |
| NADH-ubiquinone oxidoreductase chain 1 | Q01148 | 0.0068 | 0.5 |
| Phosphoenolpyruvate carboxykinase 1 | R0F3C0 | 0.0081 | 0.4 |
| Pleiotropic drug resistance protein 1 | Q76CU2 | 0.013 | 0.4 |
| T-complex protein 1 subunit beta-like isoform X1 | K3XGI3 | 0.025 | 0.4 |
| Dynamin-2B | M5VHG2 | 0.018 | 0.3 |
| Glutamate decarboxylase 1 | UPI00046E1A6C | 0.0022 | 0.1 |
| Calcium-transporting ATPase 2 | 460375325 | 0.017 | 0.1 |

| | | | |
|---|---|---|---|
| Cullin-associated and neddylation dissociated | A0A061G8M4 | 0.0025 | 0 |
| N-ethylmaleimide sensitive fusion protein | T1RU75 | 0.0071 | 0 |

**Proteins that change in abundace in Fraction T84**

| Protein name | Uniprot Accession | p-Value | Fold change |
|---|---|---|---|
| Histone H2A.6 | B6SHX9 | 0.012 | 5 |
| Catalase isozyme 1 | Q01297 | 0.041 | 5 |
| T-complex protein 1 subunit epsilon-like | A0A3Q7GD89 | 0.024 | 4.9 |
| Carbohydrate-binding-like fold-containing protein | Q9LZQ4 | 0.028 | 4.6 |
| Serine hydroxymethyltransferase 2 | C6ZJY7 | 0.0072 | 4.3 |
| 60S ribosomal protein L15-like | UPI0002767146 | 0.012 | 3.9 |
| 14-3-3-like protein | P93259 | 0.03 | 3.6 |
| Eukaryotic translation initiation factor 3 subunit C | O49160 | 0.005 | 3.3 |
| Thioredoxin H-type 1 | P29449 | 0.039 | 2.7 |
| 14-3-3-like protein | O65352 | 0.014 | 2 |
| Actin-1 | B4FPG2 | 0.029 | 1.9 |
| Ribosomal protein S5 | A0A061F8V4 | 0.0053 | 1.6 |
| Elongation factor 2 | O23755 | 0.0062 | 1.6 |
| ATP synthase subunit alpha, mitochondrial | P05492 | 0.01 | 0.3 |
| ATP synthase subunit alpha | P05492 | 0.018 | 0.3 |
| Gamma carbonic anhydrase-like 2, mitochondrial-like | A0A0A0LR76 | 0.027 | 0.3 |
| Mitochondrial-processing peptidase subunit beta-like | A0A0A0L0I1 | 0.013 | 0.2 |
| NADH dehydrogenase subunit 7 | G9JLR7 | 0.046 | 0.2 |
| Fatty acyl-CoA synthetase A | M8BUY6 | 0.0035 | 0 |

**Proteins that change in abundace in Fraction T85**

| Protein name | Uniprot Accession | p-Value | Fold change |
|---|---|---|---|
| Monocopper oxidase-like protein SKU5-like | A0A1S2Z641 | 0.007 | **INF** |
| Mitochondrial malate dehydrogenase | I6YI03 | 0.014 | **INF** |
| Elongation factor 1-delta 1 | B4FNT1 | 0.018 | 2.9 |
| Vacuolar protein sorting 13 | J7FHV5 | 0.028 | 0.7 |
| Phosphorus transporter | K4BDP2 | 0.039 | 0.7 |
| Ribosomal protein L3 | I1L7D4 | 0.012 | 0.6 |
| Gamma carbonic anhydrase-like 2 | UPI0002768717 | 0.014 | 0.4 |
| Phosphoenolpyruvate carboxykinase 1 | R0F3C0 | 0.024 | 0.3 |
| AAA-type ATPase family protein | F4JCC9 | 0.0032 | 0 |
| 40S ribosomal protein S27-1 | O64650 | 0.0044 | 0 |
| Cation/H(+) antiporter 18-like | A0A0A0KUY7 | 0.011 | 0 |

**Supplementary Material and Methods Table.** Oligonucleotides use for gene cloning.

| *Gene* | *Forward primer* | *Reverse primer* |
|---|---|---|
| *McTIP1;2* | CACCATGCCGATCAGAGACATCAG | TTAGGCAGAGAGTCTCTGGTA |
| McPIP1;4 | GGGGACAAGTTTGTACAAAAAAGCAGGCTTAATG GAGGGGAAGGAAGAGGATG | GGGGACCACTTTGTACAAGAAAGCTGGGTGCTTGG ATTTGAATGGAATTGCCCT |
| McPIP2;1 | GGGGACAAGTTTGTACAAAAAAGCAGGCTTAATG ACTAAGGACGTGGAGGTG | GGGGACAACTTTGTATAGAAAAGTTGGGTGTGCAG AGCTCCTGAAGGAAC |
| McCLC | GTACAAAAAAGCAGGCTTCATGTCTTCATTTCCCG ACTCTTT | GTACAAGAAAGCTGGGTCCGCAGCCAATACGGCCT CTG |
| McAP1µ | CACCATGTCGGGAGCGGCCTC | TCACATTAGCCTCAGCTCGTACT |
| McAP2µ | CACCATGCCGGTGGCTGCTTC | ATCACACCTGATCTCATATGAAC |
